# Lung expression of genes encoding SARS-CoV-2 cell entry molecules and antiviral restriction factors: interindividual differences are associated with age and germline variants

**DOI:** 10.1101/2020.06.24.168534

**Authors:** Chiara E. Cotroneo, Nunzia Mangano, Tommaso A. Dragani, Francesca Colombo

**Author notes:** Corresponding author: Tommaso A. Dragani, Department of Research, Fondazione IRCCS Istituto Nazionale dei Tumori, Via G.A. Amadeo 42, 20133 Milan, Italy.

## Abstract

Germline variants in genes involved in SARS-CoV-2 cell entry (i.e. *ACE2* and *TMPRSS2*) may influence the susceptibility to infection, as may polymorphisms in genes involved in the innate host response to viruses (e.g. APOBEC3 family). We searched for polymorphisms acting, in lung tissue, as expression quantitative trait loci (eQTLs) for 15 candidate COVID-19 susceptibility genes, selected for their roles in virus cell entry and host antiviral responses. No significant eQTLs were identified for *ACE2* and *TMPRSS2* genes, whose expression levels did not associate with either sex or age of the 408 patients whose non-diseased lung tissue was analyzed. Instead, we identified seven *cis*-eQTLs (FDR<0.05) for *APOBEC3D* and *APOBEC3G* (rs139296, rs9611092, rs139331, rs8177832, rs17537581, rs61362448, and rs738469). The genetic control of the expression of APOBEC3 genes, which encode enzymes that interfere with virus replication, may explain interindividual differences in risk or severity of viral infections. Future studies should investigate the role of host genetics in COVID-19 patients using a genome-wide approach, to identify other genes whose expression levels are associated with susceptibility to SARS-CoV-2 infection or COVID-19 severity.

**Author summary:** Identification of expression quantitative trait loci (eQTLs) has become commonplace in functional studies on the role of individual genetic variants in susceptibility to diseases. In COVID-19, it has been proposed that individual variants in SARS-CoV-2 cell entry and innate host response genes may influence the susceptibility to infection. We searched for polymorphisms acting, in non-diseased lung tissue of 408 patients, as eQTLs for 15 candidate COVID-19 susceptibility genes, selected for their roles in virus cell entry and host antiviral responses. Seven *cis*-eQTLs were detected for *APOBEC3D* and *APOBEC3G* genes, which encode enzymes that interfere with virus replication. No significant eQTLs were identified for *ACE2* and *TMPRSS2* genes. Therefore, the identified eQTLs may represent candidate loci modulating interindividual differences in risk or severity of SARS-CoV-2 virus infection.

## Introduction

SARS-CoV-2 infection has a wide range of clinical presentations, from asymptomatic to severe or critical, leading in some cases to death [1, 2]. Data from Italy exemplify the great variability in prognosis [3]. For example, in a sample of over 40,000 persons who tested positive for SARS-CoV-2 genome, 25.2% were asymptomatic, 55.7% had few, mild or unspecified symptoms, and 19.1% had severe symptoms requiring hospitalization. Considering all 230,000 PCR-confirmed cases in Italy, the case-fatality rate is 13.7%. Prognosis depends on age, with ∼85% of deaths registered in persons ≥70 years old. Prognosis also depends on sex, with a higher case-fatality rate for men than women (17.6% vs. 10.4%, respectively). According to a meta-analysis of more than 3,000 patients, male sex and age >65 years are poor prognostic factors [4]. Why SARS-CoV-2 infection has such variable presentations and why COVID-19 is more fatal in men than women are currently are largely unknown. Pre-existing medical conditions such as diabetes and hypertension and a smoking habit have been reported to increase the risk of severe disease [4, 5], and there is emerging evidence for the existence of at least two strains of the virus that vary in virulence [6]. Individual genetic constitution may also play a role in determining the severity of a SARS-CoV-2 infection.

The influence of genetics on pulmonary infections has already been observed. For example, susceptibility to and severity of influenza vary according to the genotype of single nucleotide polymorphisms (SNPs) in several genes involved in interactions with the influenza A virus, in inflammatory processes, and in other host responses (reviewed in [7]). These genes include *IFITM3*, a member of the interferon-induced transmembrane protein (IFITM) family, and *TMPRSS*2, which encodes a transmembrane protease whose activity is essential for viral entry. Both coding and non-coding polymorphisms in these genes have been associated with the risk or severity of influenza A infection [8-10]. These results suggest that not only genetic variants affecting protein function but also those affecting gene expression modulate the response to viral infection. For severe acute respiratory syndrome (SARS), caused by the SARS-CoV virus, the risk or severity of infection has been associated with polymorphisms in genes involved in inflammatory and immune responses [11-13] or viral entry [14].

For the new coronavirus SARS-CoV-2, two human proteins so far have been identified as being involved in viral entry and replication in cells. Angiotensin I converting enzyme 2 (ACE2), the main receptor for viral attachment [15, 16], is a transmembrane protein encoded by the *ACE2* gene. According to molecular modeling studies, two coding variants may alter the binding affinity between ACE2 and the viral spike protein (protein S) [17]. SARS-CoV-2 entry also depends on the protease encoded by *TMPRSS2*, which cleaves the viral spike protein, an essential step for adsorption of the virus to the cell [15].

In addition to the human proteins needed for viral entry, other proteins involved in viral infection are the so-called antiviral restriction factors. These components of the innate immune response system target viruses and interfere with their life cycle, for instance by modifying their proteins or RNA (reviewed in [18]). Antiviral restriction factors include members of the IFITM family, the APOBEC3 family of RNA editing enzymes, the adenosine deaminase ADAR, and members of the poly(ADP-Ribose) polymerase (PARP) family [19-21]. Coding variants in these genes may also explain the different responses to SARS-CoV-2 infection in different people. As already observed for influenza, risk or severity of infection may also depend on non-coding variants that influence gene expression, for instance by modifying the binding sites for transcription factors or altering enhancer sequences. These regulatory genetic variants are termed expression quantitative trait loci (eQTLs) because they alter gene expression in a quantitative manner [22].

To test the hypothesis that individual genetic constitution plays a role in SARS-CoV-2 infection, we examined the expression in lung of two genes known to be involved in SARS-CoV-2 infection susceptibility (i.e. *ACE2* and *TMPRSS2)* and of other genes involved in host antiviral responses. Taking advantage of an existing dataset of genotype and transcriptome data from non-involved lung tissue from patients with lung adenocarcinoma, we investigated whether the expression of these candidate genes reflected clinical characteristics known to be prognostic of SARS-CoV-2 outcome (i.e. age and sex) or was genetically controlled by *cis or trans* regulatory polymorphisms.

## Materials and Methods

### Study design

We selected 16 genes for their known involvement in SARS-CoV-2 virus adsorption or entry into human cells (2 genes) and their suspected involvement in the innate immune response to this virus (14 genes). The expression of these genes and the genotypes of germline variants were studied in apparently normal lung tissue from 408 Italian patients who had undergone lobectomy for lung adenocarcinoma.

### Ethics Statement

This case series is part of a broader research program on lung adenocarcinoma, for which a biobank of lung tissue and extracted nucleic acids is maintained at Fondazione IRCCS Istituto Nazionale dei Tumori, Milan (previously described in [23]). Methods for tissue collection, storage and nucleic acid extraction, and information about ethical approval and patients’ informed consent have already been reported [23].

### Transcriptome profiling

Lung transcriptome data, obtained using Illumina HumanHT-12 v4 Expression BeadChips, were already available in our lab (GEO database accession numbers: GSE71181 and GSE123352). Microarray raw data were subjected to pre-processing and quality checking using R version 3.6.3. Raw data were log2-transformed and normalized using the robust spline normalization method implemented in the lumi Bioconductor package [24]. Corrected data were then collapsed from probe level to gene level by selecting, for each gene, the probe with the highest mean intensity across samples.

After pre-processing, measured intensities for genes of interest were extracted and then merged after adjusting for batch effects using the ComBat function from the sva R Bioconductor package [25], using sex and age at surgery of each patient as covariates for the normalization step.

### Correlation of gene expression with sex and age

Sex and age of the patients were tested as possible linear predictors of the observed log2-transformed transcript level of each gene using a linear regression model, by the `glm` function in R version 3.6.3. Significant correlations were considered when *P*<0.05.

### SNP genotyping

Genotyping data of a subset of 201 patients were already available (described in [23]). Genome-wide genotyping of 200 ng genomic DNA from the remaining 207 patients was carried out using Infinium HumanOmni2.5-8 v1.2 BeadChip microarrays (Illumina). After completion of the assay, the BeadChips were scanned with the two-color confocal Illumina HIscan System at a 0.375 μm pixel resolution. Image intensities were extracted and genotypes were determined using Illumina’s BeadStudio software.

Genotype data were subjected to quality control using PLINK v1.90b6.16 [26] following the protocol described in [27]. All individuals whose heterozygosity rates for markers on chromosome X were inconsistent with their biological sex were discarded. Other discarded samples corresponded to individuals where 3 % of the total genotyped SNPs were missing and whose heterozygosity rates deviated by three or more standard deviations from the average heterozygosity rate of the whole study group. In the remaining samples, all genotyped SNPs with a call rate <95 %, a minor allele frequency (MAF) <2 %, or a Hardy-Weinberg equilibrium P < 1.0×10^−7^ were excluded from further analyses. Probes mapping to multiple regions were also removed, as were mitochondrial SNPs and duplicate SNPs (i.e., those with duplicate IDs or genomic positions). The resulting dataset was then converted to a tabular format with the option `--recode A-transpose`.

### eQTL analyses

To test whether gene expression in lung tissue varied with the genotype of SNPs, we carried out an eQTL analysis by standard additive linear regression model, using the MatrixEQTL package [28] in R environment, assuming that genotype has an additive effect on gene expression. Briefly, genotypes were expressed as integers (0, 1, or 2) according to the number of minor alleles at each SNP. Sex and age at surgery of each patient were used as covariates. eQTLs were considered *in cis* where the corresponding SNP was located within one megabase of the gene mapping coordinates, *and in trans* for all other cases. Correction of *P*-values for multiple testing was carried out using the Benjamini-Hochberg method [29] to obtain the false discovery rate (FDR). Genome-wide significance threshold for the analysis was set at FDR<0.05. Differences in expression levels (log2-transformed values) among the three genotype groups were tested for significance by one-way ANOVA followed by Tukey’s test for multiple comparisons (*P*<0.05 was considered as statistically significant).

To understand the tissue specificity of eQTLs, we searched the Genotype-Tissue Expression (GTEx) database [https://www.gtexportal.org/home/, GTEx Analysis Release V8, dbGaP Accession phs000424.v8.p2, accessed on 06/05/2020]. All identified cis-eQTLs were singularly searched using the variant browser, and we looked for our target genes among the single-tissue eQTL results. In particular, data (normalized effect size and *P*-value) of lung eQTLs were reported.

Linkage disequilibrium data and minor allele frequencies in the different populations were retrieved from the Ensembl genome browser. To investigate potential functional roles of the eQTLs, we used the SNP Function Prediction tool at the SNPinfo Web Server [30], with the default settings. All identified cis-eQTL SNPs were checked.

## Results

This study considered transcriptomic and genotype data from samples of apparently normal lung tissue from 408 patients with lung adenocarcinoma (**Supplementary Table 1**). The study group had a median age of 65 years (range, 36-84) and a predominance of men (67.2 %). The vast majority (88.2 %) were ever smokers and 64.6 % had pathological stage I disease.

**Table 1.** Effects of sex and age on the lung expression of genes known or suspected to be involved in COVID-19 susceptibility or severity

| Gene | Location (Chr: bp)<sup>a</sup> | Correlation with sex<sup>b</sup> |  | Correlation with age |  |
| --- | --- | --- | --- | --- | --- |
|  |  | β | *P* | β | *P* |
| SARS-CoV-2 cell entry |  |  |  |  |  |
| ACE2 | X: 15,561,033-15,602,148 | -0.025 | 0.391 | 0.0009 | 0.531 |
| TMPRSS2 | 21: 41,464,305-41,531,116 | -0.095 | 0.060 | -0.0007 | 0.801 |
| Innate immune response |  |  |  |  |  |
| ADAR | 1: 154,582,057-154,628,013 | 0.014 | 0.655 | -0.004 | 0.022 |
| APOBEC3A | 22: 38,952,741-38,992,778 | -0.065 | 0.249 | 0.004 | 0.186 |
| APOBEC3B | 22: 38,982,347-38,992,804 | 0.026 | 0.466 | 0.0002 | 0.906 |
| APOBEC3C | 22: 39,014,257-39,020,352 | 0.021 | 0.581 | -0.0002 | 0.928 |
| APOBEC3D | 22: 39,021,113-39,033,277 | 0.032 | 0.336 | 0.0002 | 0.900 |
| APOBEC3F | 22: 39,040,604-39,055,972 | -0.007 | 0.853 | 0.001 | 0.504 |
| APOBEC3G | 22: 39,077,067-39,087,743 | 0.060 | 0.229 | 0.001 | 0.690 |
| APOBEC3H | 22: 39,097,224-39,104,067 | 0.024 | 0.491 | 0.007 | <0.001 |
| IFITM2 | 11: 307,631-315,272 | -0.031 | 0.420 | 0.004 | 0.068 |
| IFITM3 | 11: 319,676-327,537 | -0.010 | 0.790 | 0.004 | 0.049 |
| PARP12 | 7: 140,023,749-140,062,951 | 0.0279 | 0.462 | -0.007 | <0.001 |
| PARP14 | 3: 122,680,839-122,730,840 | 0.065 | 0.068 | -0.004 | 0.035 |
| ST3GAL1 | 8: 133,454,848-133,571,940 | 0.082 | 0.057 | 0.002 | 0.339 |

<sup>a</sup> According to GRCh38 genome assembly; <sup>b</sup> Men as reference. Chr, chromosome.

### Effect of sex and age on gene expression

Of the selected genes, all were expressed at detectable levels in lung tissue with the exception of *IFITM1*, which did not pass pre-processing and quality checking filters. A first analysis assessed whether the expression levels of the remaining 15 genes expressed in lung associated with two clinical variables known to affect prognosis in COVID-19, i.e. sex and age (**Table 1**). No gene was differently expressed between men and women. Three genes, i.e. *ADAR* (beta = −0.004), *PARP12* (beta = −0.007) *and PARP14* (beta = −0.004), showed decreased expression in older patients. Higher expression levels of two genes, *APOBEC3H* (beta = 0.007) and *IFITM3* (beta = 0.004), were instead associated with increasing age.

### Germline variations associated with gene expression

A genome-wide eQTL analysis was carried out for the 15 genes, considering 736,210 SNPs. This analysis identified 16 *cis*-eQTLs at a nominal *P* < 1.0 × 10^−3^ associated with the expression levels of five genes (**Table 2**) and 1,170 *trans*-eQTLs at *P* < 1.0 × 10^−4^ associated with the expression levels of 15 genes (**Supplementary Table 2**).

**Table 2.**
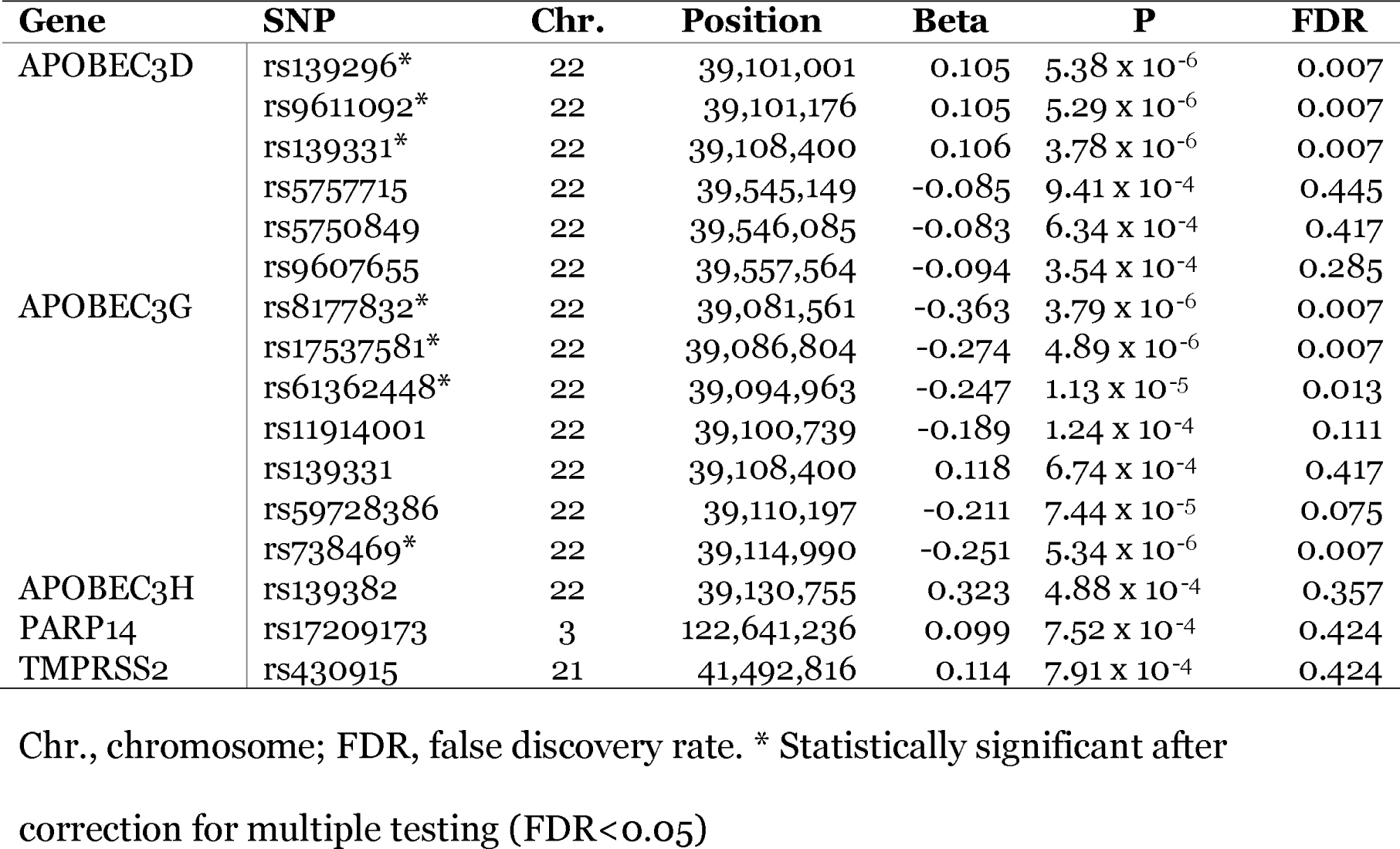
*cis*-eQTLs identified for COVID-19-related genes, in non-involved lung tissue of 408 patients with lung adenocarcinoma (*P*<10×10^−3^)

Of the 16 cis-eQTLs, after correcting for multiple testing, seven continued to meet the genome-wide significance threshold for the analysis (i.e. FDR < 0.05). These seven eQTLs were involved in the *in cis* modulation of two genes from the APOBEC3 family, i.e. *APOBEC3D* and *APOBEC3G* (**Figure 1**). For *APOBEC3D*, subjects homozygous for the major alleles of rs139296, rs9611092 and rs139331 had lower transcript levels than did heterozygotes or homozygotes for the minor allele (for all of them, *P* < 0.01, ANOVA followed by Tukey’s test for multiple comparisons; **Figure 1A-C**). For *APOBEC3G*, homozygotes for the major alleles of rs8177832, rs17537581, rs61362448 and rs738469 had higher expression levels than heterozygotes (*P* < 0.01) or homozygotes for the minor alleles (for all but rs61362448, *P* < 0.05, ANOVA followed by Tukey’s test for multiple comparisons; **Figure 1D-G**).

**Figure 1.**
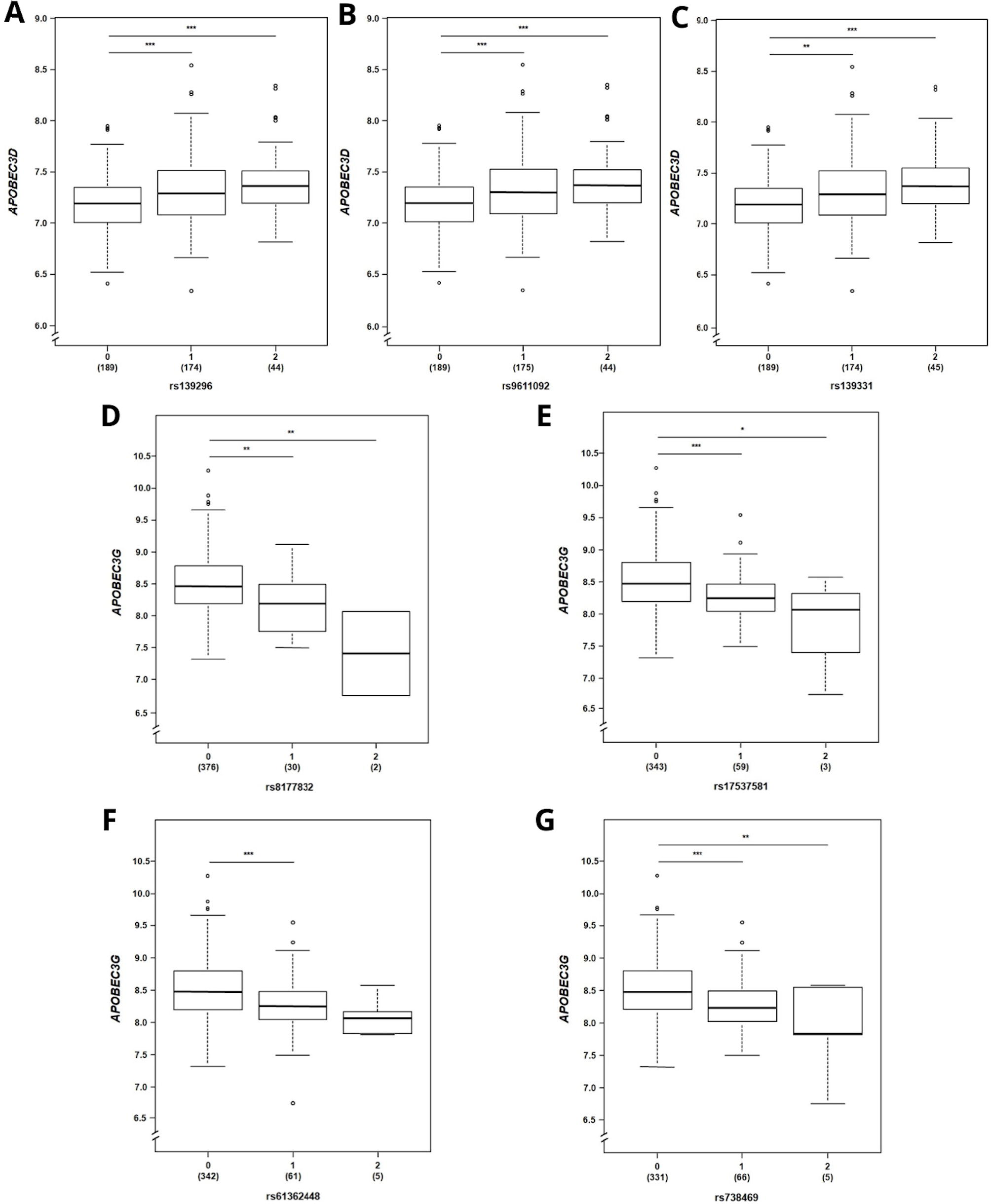
Effect of genotype on *APOBEC3D* and *APOBEC3G* transcript levels. (A-C) Expression levels (log2-transformed probe intensity) of *APOBEC3D*, according to rs139296, rs9611092, or rs139331 genotypes, respectively. (D-G) Expression levels (log2-transformed probe intensity) of *APOBEC3G*, according to rs8177832, rs17537581, rs61362448, or rs738469 genotypes, respectively. Genotypes were expressed as integers (0, 1, or 2) according to the number of minor alleles at each SNP. Numbers in parentheses are the individuals carrying the indicated genotype. The line within each box represents the median; upper and lower edges of each box are 75^th^ and 25^th^ percentiles, respectively; top and bottom whiskers indicate the greatest and least values, respectively; circles denote outliers (>1.5 times the interquartile range above the upper quartile and below the lower quartile). *** *P* < 0.001, ** *P* < 0.01, * *P* < 0.05, ANOVA followed by Tukey’s test for multiple comparisons.

The three SNPs associated with the expression of *APOBEC3D* were in tight linkage disequilibrium with each other, suggesting that they comprise a unique locus. They all map ∼80 kb downstream of *APOBEC3D*. The first two are intronic of *APOBEC3H*, and the third is intergenic. These three SNPs are frequent in all populations (**Supplementary Table 3**). In this series, the minor allele frequency (MAF) was 32 % for all of them. Regarding the eQTLs that were excluded after correction for multiple testing, two other loci (the first including the linked SNPs rs5757715 and rs5750849, and the second marked by rs9607655) associated with levels of *APOBEC3D*. Their MAFs vary widely in different populations (**Supplementary Table 3**). Both these eQTLs map more than 500 kb downstream of *APOBEC3D*. SNPs rs5757715 and rs9607655 map in regulatory regions, i.e. in an enhancer and in a CTCF binding site, respectively.

The four *APOBEC3G* eQTLs with genome-wide significance are rare variants in our series and other populations, but not in the African population (**Supplementary Table 3**). The first two variants, rs8177832 and rs17537581, map inside the *APOBEC3G* gene: the first is a missense variant in the third exon and the second is intronic. The SNP rs61362448 is an intergenic variation, located ∼7 kb downstream of *APOBEC3G*, whereas rs738469 maps more than 20 kb downstream of its target gene. Seven eQTLs for *APOBEC3G* were excluded after correction for multiple testing, including rs139331, identified here as an eQTL of *APOBEC3D* at FDR < 0.05.

Finally, 1,170 *trans*-eQTLs were identified (**Supplementary Table 2**), including 16 *trans* eQTLs for eight genes at nominal *P* < 1.0 × 10^−6^. Three of these SNPs (rs62552897, rs17612218, and rs11792875) were associated with the levels of two genes (i.e. *IFITM2* and *IFITM3*). However, none of them met the genome-wide significance threshold for the analysis (i.e. FDR < 0.05).

### Tissue specificity and possible roles of eQTLs

Four of the *cis*-eQTLs for *APOBEC3G* identified here have already been reported. According to the GTEx database, rs17537581 (intronic) and rs61362448 (intergenic) are eQTLs for *APOBEC3G* in 16 and 18 tissues, respectively, including lung. The effects of genotype at both SNPs in our study confirm those reported, with lower levels of *APOBEC3G* associated with an increasing number of minor alleles (GTEx data: rs17537581 normalized effect size, −0.30, *P* = 2.2 × 10^−12^; rs61362448 normalized effect size, −0.29, *P* = 1.1 × 10^−10^). The downstream SNP rs738469 has been reported to be an *APOBEC3G* eQTL in four tissues (*P* < 1.0 × 10^−3^), but not in lung. The SNP rs8177832 has been reported to be an *APOBEC3G* eQTL only in thyroid. The SNP found to be a *cis*-eQTL for both *APOBEC3G* and *APOBEC3D*, rs139331, has already been reported for *APOBEC3G* in 38 tissues, including lung. The effect of genotype observed in this study confirms earlier reports (GTEx data: normalized effect size, 0.25, *P* = 3.1 × 10^−20^). Neither rs119914001 nor rs59728386 have been previously reported as eQTLs of *APOBEC3G*. Also the three genome-wide significant eQTLs of *APOBEC3D* have not yet been reported in lung tissue, but two of them (rs139296 and rs139331) were previously identified as *APOBEC3D* eQTLs in 11 other tissues. Instead, the rs9611092 variant was not found in the GTEx database. Finally, rs430915, a *cis*-eQTL for *TMPRSS2* (without genome-wide significance), had already been reported, but with an opposite effect of genotype in lung tissue (GTEx data: normalized effect size, −0.11, *P* = 6.7 × 10^−8^). The other SNPs identified here as eQTLs of *APOBEC3D, APOBEC3H* and *PARP14* are novel.

According to the SNP Function Prediction tool (**Supplementary Table 3**), rs17537581 and rs61362448 are predicted to affect some transcription factor binding sites. On the other hand, the missense variant rs8177832 is predicted to be a benign non-synonymous SNP (according to Polyphen) and to affect splicing activity, being located in an exonic splicing enhancer. No other predictions were made by the SNP Function Prediction tool.

## Discussion

This study investigated the regulation of expression, in non-involved lung tissue of surgically treated lung adenocarcinoma patients, of 15 genes selected for possible involvement in SARS-CoV-2 cell entry or in the host innate immune response. First, we did not observe significant differences in the expression of these genes between sexes, and only small effects of age on expression levels were identified for five genes. From the eQTL analysis, genome-wide significant associations between seven SNPs and the expression levels of two genes were identified.

The main result of this study is the identification of *cis*-eQTLs, in lung tissue, for two members of the APOBEC3 family, i.e. *APOBEC3D* and *APOBEC3G*. This indicates that individuals with different genotypes at the identified eQTL SNPs express different mRNA levels of the *APOBEC3D* and *APOBEC3G* genes. Of note, for two of the genome-wide significant *APOBEC3G* eQTLs (rs17537581, rs61362448), our results are in agreement with those already present in the GTEx database. APOBEC3 enzymes are involved in the host innate response against retroviruses, and they interfere with the replication of RNA viruses, including some coronaviruses [31-33]. Therefore, if these enzymes were also involved in the innate response against SARS-CoV-2, individuals expressing different levels of these enzymes could respond differently to the infection and develop more or less severe forms of COVID-19. Our findings are also of relevance in other viral infections for which a role of APOBEC enzymes in the host response has already been identified, e.g. HIV-1, HBV, HCV and HCoV-NL63 [32, 34].

*ACE2* and *TMPRSS2* are the best candidates for differences in COVID-19 susceptibility, since they are directly involved in SARS-Cov-2 cell entry [15, 16]. Understanding whether or not the level of *ACE2* transcripts in lung tissue is under genetic control is important. Indeed, a genetic control of *ACE2* expression would imply that different individuals express different levels of the protein encoded by this gene and, in turn, be differently susceptible to SARS-CoV-2 infection. However, we found that *ACE2* mRNA levels were not significantly associated with any genetic variants, either in *cis* or in *trans*. This result implies either that *ACE2* expression in the non-involved lung tissue is not genetically regulated or, alternatively, that *ACE2* levels are subject to a complex genetic regulation that our study was unable to detect. Although the statistical power of our series was sufficient to detect relatively strong effects of single polymorphisms, it was under-powered to detect SNP interactions.

TMPRSS2 has been shown to be involved in SARS-CoV-2 spike protein priming and has been suggested as a target for pharmaceutically blocking viral cell entry [15]. Genetic variants affecting the expression of this gene could be expected to influence the individual susceptibility to SARS-CoV-2 infection. However, we did not identify any significant *TMPRSS2* eQTL. In the lung, *TMPRSS2* has been observed to be co-expressed with *ACE2* in type II pneumocytes [35]. Since we did not observe a genetic control of *ACE2*, it is not surprising that even the expression of *TMPRSS2* in lung tissue is not subject to strict genetic control.

The results of our eQTL analysis indicate that the expression in lung tissue of the two best candidates for COVID-19 susceptibility are not controlled by germline variants. Additionally, we did not observe any significant difference in the levels of these two transcripts either according to patient age or between males and females. Therefore, the different COVID-19 severity observed in older or male patients than younger or female patients cannot be due to sex- or age-specific expression of *ACE2* and *TMPRSS2* genes. Our findings, though excluding a possible role of *ACE2* and *TMPRSS2* regulatory variants in COVID-19 susceptibility, do not rule out the possibility of coding variants affecting the functionality of the proteins coded by these two genes. However, a study of exome sequences from almost 50,000 subjects in the UK Biobank, including 74 SARS-CoV-2-positive cases, did not find significant differences between cases and controls in genotype frequencies of coding polymorphisms in either *ACE2* or *TMPRSS2* gene [36]. These results suggest that genetic variants affecting the structure of *ACE2* and *TMPRSS2* proteins are not associated with COVID-19 risk or severity.

Therefore, to understand the genetic basis of differences in susceptibility and severity of COVID-19, other genes should be investigated. For instance, future studies could examine the genetic control of genes whose encoded proteins have been suggested, by in *silico* analyses, to interact with viral proteins [37, 38]. Also of interest is the genetic control of immunogenes coding for proteins involved in the host responses to the virus. In conclusion, a genome-wide approach, considering both coding and regulatory variants, is preferable for investigating the genetic basis of SARS-CoV-2 infection susceptibility and of inter-individual differences in COVID-19 prognosis.

## Funding

This work was supported in part by a grant from the Fondazione Regionale Ricerca Biomedica (FRRB) of the Lombardy Region, Italy, to TAD.

## Disclosure of conflicts of interest

All authors have completed the ICMJE uniform disclosure form at www.icmje.org/coi_disclosure.pdf and declare: TAD had financial support (grant) from FRRB, Italy. All other authors had no support from any organization for the submitted work. All authors had no financial relationships with any organizations that might have an interest in the submitted work in the previous three years; no other relationships or activities that could appear to have influenced the submitted work.

## Acknowledgements

The authors acknowledge the contributions of Valerie Matarese, PhD, who provided scientific editing.

## Notes

### Competing Interest Statement

The authors have declared no competing interest.

## References

1. Huang C, Wang Y, Li X, Ren L, Zhao J, et al. (2020) Clinical features of patients infected with 2019 novel coronavirus in wuhan, china. Lancet 395(10223): 497–506.

2. Guan WJ, Ni ZY, Hu Y, Liang WH, Ou CQ, et al. (2020) Clinical characteristics of coronavirus disease 2019 in china. N Engl J Med 382(18): 1708–1720.

3. Riccardo F. (2020) Epidemia COVID-19. aggiornamento nazionale. 14 maggio 2020 – ore 16:00. 2020(June/09).

4. Zheng Z, Peng F, Xu B, Zhao J, Liu H, et al. (2020) Risk factors of critical & mortal COVID-19 cases: A systematic literature review and meta-analysis. J Infect.

5. Wang B, Li R, Lu Z, Huang Y. (2020) Does comorbidity increase the risk of patients with COVID-19: Evidence from meta-analysis. Aging (Albany NY) 12(7): 6049–6057.

6. Becerra-Flores M, Cardozo T. (2020) SARS-CoV-2 viral spike G614 mutation exhibits higher case fatality rate. Int J Clin Pract : e13525.

7. Nogales A, L DeDiego M. (2019) Host single nucleotide polymorphisms modulating influenza A virus disease in humans. Pathogens 8(4): 168. doi: 10.3390/pathogens8040168.

8. Allen EK, Randolph AG, Bhangale T, Dogra P, Ohlson M, et al. (2017) SNP-mediated disruption of CTCF binding at the IFITM3 promoter is associated with risk of severe influenza in humans. Nat Med 23(8): 975–983.

9. Cheng Z, Zhou J, To KK, Chu H, Li C, et al. (2015) Identification of TMPRSS2 as a susceptibility gene for severe 2009 pandemic A(H1N1) influenza and A(H7N9) influenza. J Infect Dis 212(8): 1214–1221.

10. Everitt AR, Clare S, Pertel T, John SP, Wash RS, et al. (2012) IFITM3 restricts the morbidity and mortality associated with influenza. Nature 484(7395): 519–523.

11. Tu X, Chong WP, Zhai Y, Zhang H, Zhang F, et al. (2015) Functional polymorphisms of the CCL2 and MBL genes cumulatively increase susceptibility to severe acute respiratory syndrome coronavirus infection. J Infect 71(1): 101–109.

12. Zhu X, Wang Y, Zhang H, Liu X, Chen T, et al. (2011) Genetic variation of the human alpha-2-heremans-schmid glycoprotein (AHSG) gene associated with the risk of SARS-CoV infection. PLoS One 6(8): e23730.

13. Chan KY, Xu MS, Ching JC, So TM, Lai ST, et al. (2010) CD209 (DC-SIGN) - 336A>G promoter polymorphism and severe acute respiratory syndrome in hong kong chinese. Hum Immunol 71(7): 702–707.

14. Itoyama S, Keicho N, Quy T, Phi NC, Long HT, et al. (2004) ACE1 polymorphism and progression of SARS. Biochem Biophys Res Commun 323(3): 1124–1129.

15. Hoffmann M, Kleine-Weber H, Schroeder S, KrÃ¼ger N, Herrler T, et al. (2020) SARS-CoV-2 cell entry depends on ACE2 and TMPRSS2 and is blocked by a clinically proven protease inhibitor. Cell.

16. Zhou P, Yang XL, Wang XG, Hu B, Zhang L, et al. (2020) A pneumonia outbreak associated with a new coronavirus of probable bat origin. Nature 579(7798): 270–273.

17. Hussain M, Jabeen N, Raza F, Shabbir S, Baig AA, et al. (2020) Structural variations in human ACE2 may influence its binding with SARS-CoV-2 spike protein. J Med Virol.

18. Chemudupati M, Kenney AD, Bonifati S, Zani A, McMichael TM, et al. (2019) From APOBEC to ZAP: Diverse mechanisms used by cellular restriction factors to inhibit virus infections. Biochim Biophys Acta Mol Cell Res 1866(3): 382–394.

19. Bailey CC, Zhong G, Huang IC, Farzan M. (2014) IFITM-family proteins: The cell’s first line of antiviral defense. Annu Rev Virol 1: 261–283.

20. Heraud-Farlow JE, Walkley CR. (2016) The role of RNA editing by ADAR1 in prevention of innate immune sensing of self-RNA. J Mol Med (Berl) 94(10): 1095–1102.

21. Grunewald ME, Chen Y, Kuny C, Maejima T, Lease R, et al. (2019) The coronavirus macrodomain is required to prevent PARP-mediated inhibition of virus replication and enhancement of IFN expression. PLoS Pathog 15(5): e1007756.

22. Nica AC, Dermitzakis ET. (2013) Expression quantitative trait loci: Present and future. Philos Trans R Soc Lond B Biol Sci 368(1620): 20120362.

23. Maspero D, Dassano A, Pintarelli G, Noci S, De Cecco L, et al. (2020) Read-through transcripts in lung: Germline genetic regulation and correlation with the expression of other genes. Carcinogenesis.

24. Du P, Kibbe WA, Lin SM. (2008) Lumi: A pipeline for processing illumina microarray. Bioinformatics 24(13): 1547–1548.

25. Leek JT, Johnson WE, Parker HS, Jaffe AE, Storey JD. (2012) The sva package for removing batch effects and other unwanted variation in high-throughput experiments. Bioinformatics 28(6): 882–883.

26. Purcell S, Neale B, Todd-Brown K, Thomas L, Ferreira MA, et al. (2007) PLINK: A tool set for whole-genome association and population-based linkage analyses. Am J Hum Genet 81(3): 559–575.

27. Anderson CA, Pettersson FH, Clarke GM, Cardon LR, Morris AP, et al. (2010) Data quality control in genetic case-control association studies. Nat Protoc 5(9): 1564–1573.

28. Shabalin AA. (2012) Matrix eQTL: Ultra fast eQTL analysis via large matrix operations. Bioinformatics 28(10): 1353–1358.

29. Benjamini Y, Drai D, Elmer G, Kafkafi N, Golani I. (2001) Controlling the false discovery rate in behavior genetics research. Behav Brain Res 125(1-2): 279–284.

30. Xu Z, Taylor JA. (2009) SNPinfo: Integrating GWAS and candidate gene information into functional SNP selection for genetic association studies. Nucleic Acids Res 37(Web Server issue): W600–5.

31. Krisko JF, Begum N, Baker CE, Foster JL, Garcia JV. (2016) APOBEC3G and APOBEC3F act in concert to extinguish HIV-1 replication. J Virol 90(9): 4681–4695.

32. Milewska A, Kindler E, Vkovski P, Zeglen S, Ochman M, et al. (2018) APOBEC3-mediated restriction of RNA virus replication. Sci Rep 8(1): 5960–018-24448-2.

33. Li J, Chen Y, Li M, Carpenter MA, McDougle RM, et al. (2014) APOBEC3 multimerization correlates with HIV-1 packaging and restriction activity in living cells. J Mol Biol 426(6): 1296–1307.

34. Siriwardena SU, Chen K, Bhagwat AS. (2016) Functions and malfunctions of mammalian DNA-cytosine deaminases. Chem Rev 116(20): 12688–12710.

35. Ziegler CGK, Allon SJ, Nyquist SK, Mbano IM, Miao VN, et al. (2020) SARS-CoV-2 receptor ACE2 is an interferon-stimulated gene in human airway epithelial cells and is detected in specific cell subsets across tissues. Cell.

36. Curtis D. (2020) Coding variants in ACE2 and TMPRSS2 are not major drivers of COVID-19 severity in UK biobank subjects. MedRxiv : 2020.05.01.20085860. 10.1101/2020.05.01.20085860.

37. Srinivasan S, Cui H, Gao Z, Liu M, Lu S, et al. (2020) Structural genomics of SARS-CoV-2 indicates evolutionary conserved functional regions of viral proteins. Viruses 12(4): E360. doi: 10.3390/v12040360.

38. Guzzi PH, Mercatelli D, Ceraolo C, Giorgi FM. (2020) Master regulator analysis of the SARS-CoV-2/Human interactome. J Clin Med 9(4): E982. doi: 10.3390/jcm9040982.

