## Supplemental Table 1 for "Lung expression of genes encoding SARS-CoV-2 cell entry molecules and antiviral restriction factors: interindividual differences are associated with age and germline variants"

**Supplementary Table 1.** Clinical characteristics of the 408 lung adenocarcinoma patients whose genotype and transcriptome data were used in the study

| Characteristic | All cases (n = 408) | Women (n = 134) | Men (n = 274) |
|---|---|---|---|
| Age, years, median (range)[a] | 65 (36-84) | 64 (40-84) | 66 (36-83) |
| Smoking habit, n (%) | | | |
| Ever | 344 (88.2) | 95 (74.8) | 249 (94.7) |
| Never | 46 (11.8) | 32 (25.2) | 14 (5.3) |
| Unknown | 18 (4.4) | 7 (5.2) | 11 (4.0) |
| Pathological stage, n (%) | | | |
| I | 263 (64.6) | 101 (75.4) | 162 (59.3) |
| II | 51 (12.5) | 10 (7.5) | 41 (15.0) |
| III/IV | 93 (22.9) | 23 (17.2) | 70 (25.6) |
| Unknown | 1 (0.2) | 0 (0) | 1 (0.4) |

[a] At diagnosis
