## Supplemental Table 2 for "Lung expression of genes encoding SARS-CoV-2 cell entry molecules and antiviral restriction factors: interindividual differences are associated with age and germline variants"

**Supplementary Table 2a.** *Trans*-eQTL identified for the *ACE2* in non-involved lung tissue of 408 Italian lung adenocarcinoma patients ($P < 1.0 \times 10^{-4}$), listed in order of *P*-value.

| gene | gene chr | SNP | SNP chr | SNP position | beta | p-value | FDR |
|---|---|---|---|---|---|---|---|
| ACE2 | X | rs56003339 | 6 | 53159523 | 0.214961279 | 4.13E-06 | 0.84453778 |
| ACE2 | X | rs4713360 | 6 | 30785369 | -0.086589982 | 4.21E-06 | 0.84453778 |
| ACE2 | X | rs66539021 | 2 | 222261912 | 0.164224566 | 4.30E-06 | 0.84453778 |
| ACE2 | X | rs12617763 | 2 | 222262248 | 0.165459158 | 4.82E-06 | 0.84453778 |
| ACE2 | X | rs17673883 | 18 | 43671105 | 0.096451226 | 6.69E-06 | 0.84453778 |
| ACE2 | X | rs2834860 | 21 | 35240877 | 0.127920928 | 9.79E-06 | 0.84453778 |
| ACE2 | X | rs73192905 | 21 | 35272341 | 0.100015543 | 1.01E-05 | 0.84453778 |
| ACE2 | X | rs28780116 | 6 | 30787891 | -0.082716055 | 1.02E-05 | 0.84453778 |
| ACE2 | X | rs16897900 | 6 | 30788116 | -0.082716055 | 1.02E-05 | 0.84453778 |
| ACE2 | X | rs13201769 | 6 | 30788289 | -0.082716055 | 1.02E-05 | 0.84453778 |
| ACE2 | X | rs4711228 | 6 | 30788497 | -0.082716055 | 1.02E-05 | 0.84453778 |
| ACE2 | X | rs4713367 | 6 | 30789045 | -0.082716055 | 1.02E-05 | 0.84453778 |
| ACE2 | X | rs9295914 | 6 | 30790930 | -0.082716055 | 1.02E-05 | 0.84453778 |
| ACE2 | X | rs113137063 | 19 | 23427912 | 0.227511839 | 1.03E-05 | 0.84453778 |
| ACE2 | X | rs71356423 | 18 | 43680659 | 0.094275228 | 1.11E-05 | 0.84453778 |
| ACE2 | X | rs12526481 | 6 | 30778856 | -0.08277187 | 1.22E-05 | 0.84453778 |
| ACE2 | X | rs13206009 | 6 | 30789411 | -0.081763394 | 1.44E-05 | 0.848481351 |
| ACE2 | X | rs13201901 | 6 | 30788363 | -0.081307639 | 1.53E-05 | 0.848481351 |
| ACE2 | X | rs10831258 | 11 | 94653557 | -0.084439941 | 1.62E-05 | 0.853417593 |
| ACE2 | X | rs577388 | 6 | 85391114 | 0.092915659 | 1.65E-05 | 0.856932848 |
| ACE2 | X | rs7537969 | 1 | 98868754 | 0.234443304 | 1.75E-05 | 0.863488543 |
| ACE2 | X | rs60810174 | 1 | 188799084 | 0.259021574 | 1.78E-05 | 0.863488543 |
| ACE2 | X | rs75281696 | 20 | 6495987 | 0.190596859 | 1.88E-05 | 0.863488543 |
| ACE2 | X | rs12661731 | 6 | 30094568 | -0.093650029 | 2.00E-05 | 0.863488543 |
| ACE2 | X | rs7765810 | 6 | 30095719 | -0.093432588 | 2.12E-05 | 0.863488543 |
| ACE2 | X | rs80046975 | 19 | 23393158 | 0.226850384 | 2.13E-05 | 0.863488543 |
| ACE2 | X | rs45565532 | 2 | 222285331 | 0.141263722 | 2.41E-05 | 0.863488543 |
| ACE2 | X | rs7503195 | 17 | 39566262 | -0.081827789 | 2.48E-05 | 0.863488543 |
| ACE2 | X | rs79777402 | 4 | 81770669 | 0.152500894 | 3.30E-05 | 0.875040765 |
| ACE2 | X | rs80233133 | 15 | 38982089 | 0.276675703 | 3.46E-05 | 0.875040765 |
| ACE2 | X | rs10753872 | 1 | 200231063 | -0.077736369 | 3.61E-05 | 0.875040765 |
| ACE2 | X | rs9556258 | 13 | 93088735 | 0.087470805 | 3.62E-05 | 0.875040765 |
| ACE2 | X | rs11781000 | 8 | 29113069 | 0.077488192 | 3.95E-05 | 0.894226918 |
| ACE2 | X | rs77134256 | 16 | 79036504 | 0.129245838 | 4.49E-05 | 0.90811064 |
| ACE2 | X | rs13293756 | 9 | 135649146 | 0.10204389 | 4.56E-05 | 0.909284759 |
| ACE2 | X | rs3755154 | 2 | 215999834 | 0.082379387 | 4.66E-05 | 0.912369094 |
| ACE2 | X | rs6697889 | 1 | 168756476 | -0.07709516 | 4.74E-05 | 0.912369094 |
| ACE2 | X | rs35376774 | 7 | 18228289 | 0.089878013 | 4.96E-05 | 0.914201928 |
| ACE2 | X | rs302153 | 7 | 18227606 | 0.089747682 | 5.04E-05 | 0.914201928 |
| ACE2 | X | rs17063610 | 6 | 134292794 | 0.17481732 | 5.17E-05 | 0.914201928 |
| ACE2 | X | rs4656792 | 1 | 170634669 | 0.125098397 | 5.19E-05 | 0.914201928 |
| ACE2 | X | rs8063170 | 16 | 27123448 | -0.104450056 | 5.20E-05 | 0.914201928 |
| ACE2 | X | rs2100471 | X | 122636628 | 0.064077005 | 5.34E-05 | 0.91575718 |
| ACE2 | X | rs2285688 | 7 | 24834840 | 0.089082587 | 5.42E-05 | 0.91575718 |
| ACE2 | X | rs4787405 | 16 | 27115394 | -0.113772995 | 5.52E-05 | 0.915968818 |
| ACE2 | X | rs1277220 | 1 | 108830824 | 0.093753409 | 5.60E-05 | 0.918110223 |
| ACE2 | X | rs9421581 | 10 | 87091942 | 0.116820598 | 6.27E-05 | 0.921112474 |
| ACE2 | X | rs34978805 | 13 | 29592749 | 0.155443265 | 6.39E-05 | 0.924166007 |

**Supplementary Table 2a.** *Trans*-eQTL identified for the *ACE2* in non-involved lung tissue of 408 Italian lung adenocarcinoma patients ($P < 1.0 \times 10^{-4}$), listed in order of *P*-value.

| gene | gene chr | SNP | SNP chr | SNP position | beta | p-value | FDR |
|---|---|---|---|---|---|---|---|
| ACE2 | X | rs1926184 | 10 | 88882644 | 0.34452858 | 6.60E-05 | 0.924166007 |
| ACE2 | X | rs28416625 | 8 | 72228508 | -0.120029103 | 6.89E-05 | 0.924166007 |
| ACE2 | X | rs8068277 | 17 | 73265683 | 0.090649097 | 6.90E-05 | 0.924166007 |
| ACE2 | X | rs2080658 | 9 | 94231316 | -0.091049304 | 7.10E-05 | 0.924166007 |
| ACE2 | X | rs9355642 | 6 | 167403361 | -0.154852991 | 7.36E-05 | 0.924166007 |
| ACE2 | X | rs12935391 | 16 | 6775364 | 0.076414806 | 7.65E-05 | 0.924166007 |
| ACE2 | X | rs1178313 | 7 | 18204417 | 0.086525452 | 7.72E-05 | 0.924166007 |
| ACE2 | X | rs10955984 | 8 | 120654601 | 0.07947808 | 7.73E-05 | 0.924166007 |
| ACE2 | X | rs13382096 | 19 | 23293628 | 0.20414605 | 7.90E-05 | 0.924166007 |
| ACE2 | X | rs16863631 | 2 | 222272133 | 0.154719499 | 8.01E-05 | 0.924166007 |
| ACE2 | X | rs7191315 | 16 | 6763911 | 0.075614281 | 8.21E-05 | 0.924166007 |
| ACE2 | X | rs9572568 | 13 | 70919642 | -0.086912241 | 8.93E-05 | 0.924166007 |
| ACE2 | X | rs2014978 | 7 | 71261393 | 0.079457961 | 9.08E-05 | 0.924166007 |
| ACE2 | X | rs4936490 | 11 | 119636039 | 0.108696156 | 9.21E-05 | 0.924166007 |
| ACE2 | X | rs898227 | 18 | 77091880 | 0.082265457 | 9.21E-05 | 0.924166007 |
| ACE2 | X | rs4526031 | 4 | 55015818 | 0.153070704 | 9.25E-05 | 0.924166007 |
| ACE2 | X | rs17738628 | 4 | 143360852 | -0.097450606 | 9.27E-05 | 0.924166007 |
| ACE2 | X | rs919123 | 2 | 64326306 | 0.07356368 | 9.38E-05 | 0.924166007 |
| ACE2 | X | rs9988929 | 12 | 10605929 | -0.075873504 | 9.80E-05 | 0.924166007 |
| ACE2 | X | rs1189830 | 14 | 57076414 | 0.091133183 | 9.87E-05 | 0.924166007 |
| ACE2 | X | rs1189837 | 14 | 57081434 | 0.091133183 | 9.87E-05 | 0.924166007 |

**Supplementary Table 2b.** *Trans*-eQTL identified for the *ADAR* in non-involved lung tissue of 408 Italian lung adenocarcinoma patients ($P < 1.0 \times 10^{-4}$), listed in order of P-value.

| gene | gene chr | SNP | SNP chr | SNP position | beta | p-value | FDR |
|---|---|---|---|---|---|---|---|
| ADAR | 1 | rs60428151 | 6 | 161889295 | 0.178300798 | 1.99E-06 | 0.807441011 |
| ADAR | 1 | rs2446649 | 6 | 39620342 | -0.114219213 | 2.29E-06 | 0.812807829 |
| ADAR | 1 | rs115466749 | 5 | 180747782 | -0.311908685 | 2.88E-06 | 0.837668746 |
| ADAR | 1 | rs2749391 | 20 | 13064789 | 0.110498905 | 3.95E-06 | 0.84453778 |
| ADAR | 1 | rs78389024 | 13 | 51732860 | 0.153791533 | 7.14E-06 | 0.84453778 |
| ADAR | 1 | rs61453118 | 5 | 180772256 | -0.252884728 | 7.48E-06 | 0.84453778 |
| ADAR | 1 | rs17423839 | 7 | 32100914 | 0.085658081 | 8.47E-06 | 0.84453778 |
| ADAR | 1 | rs6006849 | 22 | 44708486 | 0.094833647 | 8.74E-06 | 0.84453778 |
| ADAR | 1 | rs7716977 | 5 | 178878049 | 0.270528744 | 1.07E-05 | 0.84453778 |
| ADAR | 1 | rs73020577 | 6 | 161885906 | 0.164006374 | 1.10E-05 | 0.84453778 |
| ADAR | 1 | rs17602954 | 13 | 58685368 | 0.178391969 | 1.12E-05 | 0.84453778 |
| ADAR | 1 | rs76459215 | 5 | 180781343 | -0.246888432 | 1.22E-05 | 0.84453778 |
| ADAR | 1 | rs61783240 | 1 | 76835373 | 0.244300865 | 1.47E-05 | 0.848481351 |
| ADAR | 1 | rs678278 | 6 | 153157422 | 0.08745639 | 1.53E-05 | 0.848481351 |
| ADAR | 1 | rs2162289 | 12 | 108289279 | 0.084764954 | 1.85E-05 | 0.863488543 |
| ADAR | 1 | rs2305339 | 4 | 61934836 | -0.094729067 | 2.06E-05 | 0.863488543 |
| ADAR | 1 | rs112797886 | 6 | 39581823 | -0.124721227 | 2.29E-05 | 0.863488543 |
| ADAR | 1 | rs1124016 | 6 | 153162468 | 0.085732602 | 2.34E-05 | 0.863488543 |
| ADAR | 1 | rs10810942 | 9 | 18280145 | -0.107869857 | 2.38E-05 | 0.863488543 |
| ADAR | 1 | rs1609127 | 5 | 18353145 | -0.090497431 | 2.97E-05 | 0.875040765 |
| ADAR | 1 | rs2814142 | 6 | 65732603 | 0.107673928 | 3.44E-05 | 0.875040765 |
| ADAR | 1 | rs7569382 | 2 | 45253003 | 0.109548409 | 3.56E-05 | 0.875040765 |
| ADAR | 1 | rs86857 | 1 | 171626947 | -0.095018826 | 3.61E-05 | 0.875040765 |
| ADAR | 1 | rs235861 | 1 | 171630087 | -0.095018826 | 3.61E-05 | 0.875040765 |
| ADAR | 1 | rs608994 | 20 | 13000237 | -0.090877442 | 3.64E-05 | 0.875040765 |
| ADAR | 1 | rs10859482 | 12 | 93314030 | 0.094694405 | 3.78E-05 | 0.879948067 |
| ADAR | 1 | rs13043334 | 20 | 50374678 | -0.092820821 | 3.99E-05 | 0.894226918 |
| ADAR | 1 | rs61879374 | 11 | 19834357 | -0.12862756 | 4.13E-05 | 0.896458005 |
| ADAR | 1 | rs76988909 | 6 | 39598877 | -0.120710663 | 4.23E-05 | 0.896458005 |
| ADAR | 1 | rs75176168 | 1 | 193431230 | 0.20178629 | 4.57E-05 | 0.909284759 |
| ADAR | 1 | rs80177306 | 9 | 38411686 | 0.254530434 | 4.79E-05 | 0.913006272 |
| ADAR | 1 | rs75570271 | 2 | 173402962 | 0.114779802 | 4.90E-05 | 0.914201928 |
| ADAR | 1 | rs73100658 | 7 | 32107879 | 0.078656656 | 5.16E-05 | 0.914201928 |
| ADAR | 1 | rs17339606 | 7 | 32117504 | 0.078656656 | 5.16E-05 | 0.914201928 |
| ADAR | 1 | rs7573219 | 2 | 45254043 | 0.107426066 | 5.44E-05 | 0.91575718 |
| ADAR | 1 | rs10498743 | 6 | 39633412 | -0.125866951 | 5.47E-05 | 0.91575718 |
| ADAR | 1 | rs111594086 | 6 | 39651187 | -0.125866951 | 5.47E-05 | 0.91575718 |
| ADAR | 1 | rs112747422 | 6 | 39658381 | -0.125866951 | 5.47E-05 | 0.91575718 |
| ADAR | 1 | rs755486 | 9 | 77583675 | 0.080983289 | 5.63E-05 | 0.918110223 |
| ADAR | 1 | rs2287958 | 19 | 422543 | -0.084112381 | 5.70E-05 | 0.918110223 |
| ADAR | 1 | rs6597061 | 6 | 495636 | -0.185868805 | 5.87E-05 | 0.918110223 |
| ADAR | 1 | rs4695002 | 4 | 44354115 | -0.1141655 | 5.99E-05 | 0.918110223 |
| ADAR | 1 | rs6447330 | 4 | 44356395 | -0.1141655 | 5.99E-05 | 0.918110223 |
| ADAR | 1 | rs17599823 | 4 | 44361470 | -0.1141655 | 5.99E-05 | 0.918110223 |
| ADAR | 1 | rs6456021 | 6 | 165852618 | 0.113415597 | 6.10E-05 | 0.918110223 |
| ADAR | 1 | rs111955783 | 5 | 18255787 | -0.080812922 | 6.18E-05 | 0.919202897 |
| ADAR | 1 | rs79107604 | 8 | 16394573 | -0.190225921 | 6.19E-05 | 0.919360784 |
| ADAR | 1 | rs6849081 | 4 | 69144840 | 0.089365229 | 6.37E-05 | 0.924166007 |

**Supplementary Table 2b.** *Trans*-eQTL identified for the *ADAR* in non-involved lung tissue of 408 Italian lung adenocarcinoma patients ($P < 1.0 \times 10^{-4}$), listed in order of P-value.

| gene | gene chr | SNP | SNP chr | SNP position | beta | p-value | FDR |
|---|---|---|---|---|---|---|---|
| ADAR | 1 | rs243888 | 20 | 13072387 | 0.08359091 | 6.45E-05 | 0.924166007 |
| ADAR | 1 | rs2937039 | 5 | 18315689 | -0.086914222 | 6.61E-05 | 0.924166007 |
| ADAR | 1 | rs2044577 | 3 | 126271174 | 0.080879201 | 6.74E-05 | 0.924166007 |
| ADAR | 1 | rs4756261 | 11 | 35943608 | -0.100454615 | 7.37E-05 | 0.924166007 |
| ADAR | 1 | rs2756018 | 10 | 33057769 | 0.087871992 | 7.40E-05 | 0.924166007 |
| ADAR | 1 | rs922628 | 12 | 66001055 | -0.088455977 | 7.56E-05 | 0.924166007 |
| ADAR | 1 | rs6796443 | 3 | 28840396 | 0.080422046 | 7.74E-05 | 0.924166007 |
| ADAR | 1 | rs12516366 | 5 | 180779397 | -0.215976726 | 8.14E-05 | 0.924166007 |
| ADAR | 1 | rs76370846 | 9 | 38403199 | 0.236880992 | 8.14E-05 | 0.924166007 |
| ADAR | 1 | rs7421861 | 2 | 241853198 | -0.085453061 | 8.58E-05 | 0.924166007 |
| ADAR | 1 | rs2814140 | 6 | 65733257 | 0.101703614 | 8.76E-05 | 0.924166007 |
| ADAR | 1 | rs724101 | 6 | 39638859 | -0.121992895 | 8.83E-05 | 0.924166007 |
| ADAR | 1 | rs113484329 | 6 | 39713438 | -0.121992895 | 8.83E-05 | 0.924166007 |
| ADAR | 1 | rs375807 | 2 | 203825857 | 0.114306985 | 8.91E-05 | 0.924166007 |
| ADAR | 1 | rs55874361 | 2 | 45252452 | 0.103890366 | 8.97E-05 | 0.924166007 |
| ADAR | 1 | rs6572744 | 14 | 51303147 | -0.084155984 | 9.36E-05 | 0.924166007 |
| ADAR | 1 | rs7843820 | 8 | 16397864 | -0.183716878 | 9.37E-05 | 0.924166007 |
| ADAR | 1 | rs12462724 | 19 | 417556 | -0.079027524 | 9.54E-05 | 0.924166007 |
| ADAR | 1 | rs2938462 | 5 | 18262936 | -0.078094525 | 9.79E-05 | 0.924166007 |
| ADAR | 1 | rs8018254 | 14 | 80543909 | 0.174169011 | 9.91E-05 | 0.924166007 |

**Supplementary Table 2c.** *Trans*-eQTL identified for the *APOBEC3A* in non-involved lung tissue of 408 Italian lung adenocarcinoma patients ($P < 1.0 \times 10^{-4}$), listed in order of *P*-value.

| gene | gene chr | SNP | SNP chr | SNP position | beta | p-value | FDR |
|---|---|---|---|---|---|---|---|
| APOBEC3A | 22 | rs741216 | 4 | 13765889 | 0.693297265 | 1.11E-07 | 0.493509947 |
| APOBEC3A | 22 | rs74459109 | 14 | 46237744 | 0.444407178 | 3.83E-07 | 0.65901749 |
| APOBEC3A | 22 | rs17116722 | 14 | 45905968 | 0.428509725 | 4.37E-07 | 0.65901749 |
| APOBEC3A | 22 | rs1508335 | 4 | 76297087 | 0.188546904 | 1.43E-06 | 0.716570532 |
| APOBEC3A | 22 | rs10483595 | 14 | 46239055 | 0.404438372 | 2.48E-06 | 0.812807829 |
| APOBEC3A | 22 | rs74371034 | 14 | 45993630 | 0.378504611 | 2.80E-06 | 0.834550446 |
| APOBEC3A | 22 | rs11564636 | 19 | 48062226 | 0.367316752 | 5.09E-06 | 0.84453778 |
| APOBEC3A | 22 | rs1037855 | 4 | 43925817 | 0.169958657 | 5.29E-06 | 0.84453778 |
| APOBEC3A | 22 | rs17091067 | 10 | 113860349 | 0.292515187 | 5.88E-06 | 0.84453778 |
| APOBEC3A | 22 | rs4692427 | 4 | 26189744 | -0.168442144 | 1.09E-05 | 0.84453778 |
| APOBEC3A | 22 | rs116788727 | 3 | 22559606 | 0.553268037 | 1.45E-05 | 0.848481351 |
| APOBEC3A | 22 | rs58664467 | 17 | 79180216 | 0.292808109 | 1.46E-05 | 0.848481351 |
| APOBEC3A | 22 | rs114542428 | 6 | 33397947 | 0.356188423 | 1.46E-05 | 0.848481351 |
| APOBEC3A | 22 | rs7330056 | 13 | 32141665 | 0.169114157 | 1.63E-05 | 0.855831907 |
| APOBEC3A | 22 | rs36099140 | 4 | 152282280 | 0.329672195 | 1.74E-05 | 0.863488543 |
| APOBEC3A | 22 | rs12633890 | 3 | 86160127 | -0.161919563 | 1.92E-05 | 0.863488543 |
| APOBEC3A | 22 | rs10034002 | 4 | 26194433 | -0.162953845 | 1.97E-05 | 0.863488543 |
| APOBEC3A | 22 | rs10031048 | 4 | 41478873 | 0.278002493 | 2.25E-05 | 0.863488543 |
| APOBEC3A | 22 | rs12644632 | 4 | 43911448 | 0.160685974 | 2.28E-05 | 0.863488543 |
| APOBEC3A | 22 | rs935865 | 2 | 174781109 | -0.158319873 | 2.31E-05 | 0.863488543 |
| APOBEC3A | 22 | rs13093222 | 3 | 86155717 | -0.160489005 | 2.40E-05 | 0.863488543 |
| APOBEC3A | 22 | rs115019778 | 1 | 165167580 | 0.514043741 | 2.41E-05 | 0.863488543 |
| APOBEC3A | 22 | rs12402150 | 1 | 4690207 | 0.402950634 | 2.51E-05 | 0.863488543 |
| APOBEC3A | 22 | rs7624 | 19 | 34733565 | 0.16247694 | 2.69E-05 | 0.868532829 |
| APOBEC3A | 22 | rs387220 | 19 | 56009656 | 0.348466401 | 2.76E-05 | 0.875040765 |
| APOBEC3A | 22 | rs78271556 | 12 | 124202322 | 0.305000036 | 2.78E-05 | 0.875040765 |
| APOBEC3A | 22 | rs896615 | 15 | 86769490 | 0.156274397 | 2.96E-05 | 0.875040765 |
| APOBEC3A | 22 | rs75247713 | 4 | 152288388 | 0.397632253 | 3.01E-05 | 0.875040765 |
| APOBEC3A | 22 | rs74238506 | 12 | 93300272 | 0.296315444 | 3.03E-05 | 0.875040765 |
| APOBEC3A | 22 | rs12476824 | 2 | 45186255 | -0.198639944 | 3.03E-05 | 0.875040765 |
| APOBEC3A | 22 | rs58871612 | 2 | 45280150 | 0.547497189 | 3.07E-05 | 0.875040765 |
| APOBEC3A | 22 | rs73119682 | 7 | 46495224 | 0.56271102 | 3.23E-05 | 0.875040765 |
| APOBEC3A | 22 | rs73090281 | 1 | 208732043 | 0.35704064 | 3.25E-05 | 0.875040765 |
| APOBEC3A | 22 | rs896614 | 15 | 86764614 | 0.153792025 | 3.34E-05 | 0.875040765 |
| APOBEC3A | 22 | rs6841440 | 4 | 26044075 | 0.321677046 | 3.37E-05 | 0.875040765 |
| APOBEC3A | 22 | rs2498335 | X | 87358832 | 0.129765152 | 3.50E-05 | 0.875040765 |
| APOBEC3A | 22 | rs61879374 | 11 | 19834357 | 0.240154822 | 3.75E-05 | 0.879948067 |
| APOBEC3A | 22 | rs10017756 | 4 | 26196251 | -0.157634047 | 3.77E-05 | 0.879948067 |
| APOBEC3A | 22 | rs28711142 | 19 | 56011602 | 0.337955836 | 4.13E-05 | 0.896458005 |
| APOBEC3A | 22 | rs59108675 | 4 | 152257801 | 0.270073602 | 4.16E-05 | 0.896458005 |
| APOBEC3A | 22 | rs5909133 | X | 19867530 | 0.203195766 | 4.44E-05 | 0.90811064 |
| APOBEC3A | 22 | rs866062 | 3 | 23056593 | 0.154133681 | 4.46E-05 | 0.90811064 |
| APOBEC3A | 22 | rs17284977 | X | 102867283 | 0.140949864 | 4.75E-05 | 0.912369094 |
| APOBEC3A | 22 | rs2064302 | 6 | 556106 | 0.414132512 | 4.75E-05 | 0.912369094 |
| APOBEC3A | 22 | rs2689694 | 10 | 94178187 | 0.155689921 | 4.87E-05 | 0.914201928 |
| APOBEC3A | 22 | rs878581 | 11 | 19506456 | -0.162225219 | 4.90E-05 | 0.914201928 |
| APOBEC3A | 22 | rs73562055 | 11 | 99702254 | 0.337670686 | 4.94E-05 | 0.914201928 |
| APOBEC3A | 22 | rs60896611 | X | 92092718 | 0.461380115 | 5.13E-05 | 0.914201928 |

**Supplementary Table 2c.** *Trans*-eQTL identified for the *APOBEC3A* in non-involved lung tissue of 408 Italian lung adenocarcinoma patients ($P < 1.0 \times 10^{-4}$), listed in order of *P*-value.

| gene | gene chr | SNP | SNP chr | SNP position | beta | p-value | FDR |
|---|---|---|---|---|---|---|---|
| APOBEC3A | 22 | rs72747142 | 15 | 56822035 | 0.228903499 | 5.15E-05 | 0.914201928 |
| APOBEC3A | 22 | rs6844857 | 4 | 41481527 | 0.268061302 | 5.25E-05 | 0.91575718 |
| APOBEC3A | 22 | rs16957160 | 17 | 8091107 | 0.243057851 | 5.27E-05 | 0.91575718 |
| APOBEC3A | 22 | rs1031800 | 6 | 113828821 | -0.177922899 | 5.31E-05 | 0.91575718 |
| APOBEC3A | 22 | rs73988067 | 17 | 31445416 | 0.341664719 | 5.47E-05 | 0.91575718 |
| APOBEC3A | 22 | rs8046892 | 16 | 31603872 | 0.238483793 | 5.69E-05 | 0.918110223 |
| APOBEC3A | 22 | rs12069043 | 1 | 208695321 | 0.356873972 | 5.76E-05 | 0.918110223 |
| APOBEC3A | 22 | rs114429968 | 6 | 18561457 | 0.458745349 | 5.78E-05 | 0.918110223 |
| APOBEC3A | 22 | rs837870 | 2 | 129336630 | 0.35514119 | 5.79E-05 | 0.918110223 |
| APOBEC3A | 22 | rs60350712 | 3 | 40560563 | 0.390527925 | 5.89E-05 | 0.918110223 |
| APOBEC3A | 22 | rs4871742 | 8 | 126843320 | 0.212943329 | 5.91E-05 | 0.918110223 |
| APOBEC3A | 22 | rs3812358 | 7 | 29271851 | 0.234764413 | 6.02E-05 | 0.918110223 |
| APOBEC3A | 22 | rs881183 | 11 | 32068647 | 0.513941741 | 6.10E-05 | 0.918110223 |
| APOBEC3A | 22 | rs6713970 | 2 | 212972582 | 0.347410585 | 6.27E-05 | 0.921112474 |
| APOBEC3A | 22 | rs2276979 | 4 | 163494582 | 0.152835953 | 6.55E-05 | 0.924166007 |
| APOBEC3A | 22 | rs4822921 | 22 | 27592781 | 0.320048643 | 6.57E-05 | 0.924166007 |
| APOBEC3A | 22 | rs59218805 | 9 | 130429625 | 0.210429756 | 6.63E-05 | 0.924166007 |
| APOBEC3A | 22 | rs11805514 | 1 | 208745317 | 0.346535667 | 6.65E-05 | 0.924166007 |
| APOBEC3A | 22 | rs848194 | 1 | 15958847 | -0.154211267 | 6.89E-05 | 0.924166007 |
| APOBEC3A | 22 | rs72807066 | 2 | 56813532 | 0.370860001 | 6.89E-05 | 0.924166007 |
| APOBEC3A | 22 | rs72804425 | 10 | 58796818 | 0.301192554 | 6.91E-05 | 0.924166007 |
| APOBEC3A | 22 | rs55881012 | 2 | 19519299 | 0.149929971 | 6.93E-05 | 0.924166007 |
| APOBEC3A | 22 | rs1344653 | 2 | 19531084 | 0.149929971 | 6.93E-05 | 0.924166007 |
| APOBEC3A | 22 | rs9952006 | 18 | 9135889 | 0.154598252 | 6.95E-05 | 0.924166007 |
| APOBEC3A | 22 | rs12431886 | 14 | 76342591 | 0.180748332 | 6.98E-05 | 0.924166007 |
| APOBEC3A | 22 | rs1364614 | 9 | 21115003 | -0.180009907 | 7.03E-05 | 0.924166007 |
| APOBEC3A | 22 | rs76344454 | 6 | 41433670 | 0.197587386 | 7.12E-05 | 0.924166007 |
| APOBEC3A | 22 | rs61979383 | 14 | 76365871 | 0.185191805 | 7.17E-05 | 0.924166007 |
| APOBEC3A | 22 | rs6816526 | 4 | 163528121 | 0.152189813 | 7.17E-05 | 0.924166007 |
| APOBEC3A | 22 | rs12500221 | 4 | 163514951 | 0.152253239 | 7.25E-05 | 0.924166007 |
| APOBEC3A | 22 | rs2032060 | 9 | 21114203 | -0.179310439 | 7.38E-05 | 0.924166007 |
| APOBEC3A | 22 | rs56086333 | 17 | 79172582 | 0.261117679 | 7.38E-05 | 0.924166007 |
| APOBEC3A | 22 | rs7092312 | 10 | 29320342 | -0.187157614 | 7.39E-05 | 0.924166007 |
| APOBEC3A | 22 | rs79785209 | 22 | 23904378 | 0.304905836 | 7.40E-05 | 0.924166007 |
| APOBEC3A | 22 | rs9533759 | 13 | 44064518 | -0.15692391 | 7.45E-05 | 0.924166007 |
| APOBEC3A | 22 | rs8058507 | 16 | 11384988 | 0.217362369 | 7.47E-05 | 0.924166007 |
| APOBEC3A | 22 | rs75634332 | 4 | 152285364 | 0.363985791 | 7.53E-05 | 0.924166007 |
| APOBEC3A | 22 | rs4757815 | 11 | 19503509 | -0.156287847 | 7.53E-05 | 0.924166007 |
| APOBEC3A | 22 | rs848210 | 1 | 15933318 | -0.152857982 | 7.71E-05 | 0.924166007 |
| APOBEC3A | 22 | rs7374138 | 3 | 38564241 | 0.170997553 | 7.77E-05 | 0.924166007 |
| APOBEC3A | 22 | rs12450227 | 17 | 65102508 | 0.145366006 | 7.79E-05 | 0.924166007 |
| APOBEC3A | 22 | rs11824559 | 11 | 106069633 | -0.236449243 | 7.87E-05 | 0.924166007 |
| APOBEC3A | 22 | rs28647489 | 4 | 186609868 | 0.21175693 | 7.88E-05 | 0.924166007 |
| APOBEC3A | 22 | rs17704933 | 7 | 13254217 | 0.244832035 | 8.01E-05 | 0.924166007 |
| APOBEC3A | 22 | rs12973146 | 19 | 652739 | 0.149003964 | 8.03E-05 | 0.924166007 |
| APOBEC3A | 22 | rs57433885 | 8 | 54560987 | -0.14847931 | 8.23E-05 | 0.924166007 |
| APOBEC3A | 22 | rs12487032 | 3 | 151832171 | -0.147922223 | 8.51E-05 | 0.924166007 |
| APOBEC3A | 22 | rs1927369 | 13 | 102483093 | 0.194030578 | 8.70E-05 | 0.924166007 |

**Supplementary Table 2c.** *Trans*-eQTL identified for the *APOBEC3A* in non-involved lung tissue of 408 Italian lung adenocarcinoma patients ($P < 1.0 \times 10^{-4}$), listed in order of *P*-value.

| gene | gene chr | SNP | SNP chr | SNP position | beta | p-value | FDR |
|---|---|---|---|---|---|---|---|
| APOBEC3A | 22 | rs1044722 | 4 | 163526824 | 0.150309663 | 9.35E-05 | 0.924166007 |
| APOBEC3A | 22 | rs6585251 | 10 | 113891494 | 0.177866244 | 9.36E-05 | 0.924166007 |
| APOBEC3A | 22 | rs11795225 | 9 | 21106902 | -0.176914341 | 9.37E-05 | 0.924166007 |
| APOBEC3A | 22 | rs4844721 | 1 | 208744446 | 0.316956683 | 9.54E-05 | 0.924166007 |
| APOBEC3A | 22 | rs7494445 | 14 | 98984579 | 0.172055935 | 9.67E-05 | 0.924166007 |
| APOBEC3A | 22 | rs13257199 | 8 | 86785345 | 0.150876062 | 9.70E-05 | 0.924166007 |
| APOBEC3A | 22 | rs12970386 | 18 | 9722052 | 0.178208712 | 9.77E-05 | 0.924166007 |
| APOBEC3A | 22 | rs848214 | 1 | 15938927 | -0.151080088 | 9.83E-05 | 0.924166007 |
| APOBEC3A | 22 | rs1349654 | 15 | 86765704 | 0.144294534 | 9.86E-05 | 0.924166007 |
| APOBEC3A | 22 | rs11264610 | 1 | 157075932 | -0.193572501 | 9.91E-05 | 0.924166007 |

**Supplementary Table 2d.** *Trans*-eQTL identified for the *APOBEC3B* in non-involved lung tissue of 408 Italian lung adenocarcinoma patients ($P < 1.0 \times 10^{-4}$), listed in order of *P*-value.

| gene | gene chr | SNP | SNP chr | SNP position | beta | p-value | FDR |
|---|---|---|---|---|---|---|---|
| APOBEC3B | 22 | rs13010450 | 2 | 109685950 | -0.12713293 | 1.04E-06 | 0.65901749 |
| APOBEC3B | 22 | rs73233037 | 12 | 126586236 | 0.185129543 | 5.05E-06 | 0.84453778 |
| APOBEC3B | 22 | rs17114816 | 1 | 57016210 | 0.171978176 | 9.55E-06 | 0.84453778 |
| APOBEC3B | 22 | rs5923964 | X | 87325842 | 0.107636489 | 9.82E-06 | 0.84453778 |
| APOBEC3B | 22 | rs5922424 | X | 87331924 | 0.107636489 | 9.82E-06 | 0.84453778 |
| APOBEC3B | 22 | rs117683188 | 12 | 132002353 | 0.206835105 | 9.98E-06 | 0.84453778 |
| APOBEC3B | 22 | rs16857234 | 1 | 166513516 | -0.141666534 | 1.02E-05 | 0.84453778 |
| APOBEC3B | 22 | rs1078756 | 14 | 103595309 | 0.125040232 | 1.09E-05 | 0.84453778 |
| APOBEC3B | 22 | rs73166216 | 3 | 159235555 | 0.355053778 | 1.10E-05 | 0.84453778 |
| APOBEC3B | 22 | rs73229843 | 8 | 22643896 | 0.361909197 | 1.28E-05 | 0.848481351 |
| APOBEC3B | 22 | rs58533102 | 2 | 109680644 | -0.116273919 | 1.30E-05 | 0.848481351 |
| APOBEC3B | 22 | rs7037327 | 9 | 122541481 | 0.150234295 | 1.32E-05 | 0.848481351 |
| APOBEC3B | 22 | rs61875405 | 11 | 24354813 | -0.151889458 | 1.36E-05 | 0.848481351 |
| APOBEC3B | 22 | rs502611 | 11 | 132720691 | 0.112410251 | 1.51E-05 | 0.848481351 |
| APOBEC3B | 22 | rs9863882 | 3 | 187463294 | -0.097940197 | 1.74E-05 | 0.863488543 |
| APOBEC3B | 22 | rs12965511 | 18 | 50034470 | 0.230032298 | 1.88E-05 | 0.863488543 |
| APOBEC3B | 22 | rs17495782 | 17 | 41476088 | -0.110707065 | 2.17E-05 | 0.863488543 |
| APOBEC3B | 22 | rs6571647 | 14 | 34297878 | 0.195280927 | 2.25E-05 | 0.863488543 |
| APOBEC3B | 22 | rs10766627 | 11 | 20113323 | 0.138115624 | 2.30E-05 | 0.863488543 |
| APOBEC3B | 22 | rs35577195 | 6 | 89180699 | 0.283206969 | 2.44E-05 | 0.863488543 |
| APOBEC3B | 22 | rs201621994 | X | 47107377 | 0.231566371 | 2.45E-05 | 0.863488543 |
| APOBEC3B | 22 | rs6668180 | 1 | 56974569 | 0.162971687 | 2.56E-05 | 0.863488543 |
| APOBEC3B | 22 | rs62213555 | 21 | 45792616 | -0.115506938 | 2.60E-05 | 0.863488543 |
| APOBEC3B | 22 | rs11601279 | 11 | 81289288 | -0.145473117 | 2.72E-05 | 0.872704818 |
| APOBEC3B | 22 | rs6426676 | 1 | 21227826 | 0.331535291 | 2.83E-05 | 0.875040765 |
| APOBEC3B | 22 | rs7048825 | 9 | 68981049 | 0.150712244 | 2.99E-05 | 0.875040765 |
| APOBEC3B | 22 | rs969968 | 11 | 83923187 | 0.248146235 | 3.23E-05 | 0.875040765 |
| APOBEC3B | 22 | rs7091633 | 10 | 14428921 | 0.163516049 | 3.28E-05 | 0.875040765 |
| APOBEC3B | 22 | rs6736742 | 2 | 217412426 | 0.108743094 | 3.50E-05 | 0.875040765 |
| APOBEC3B | 22 | rs2839005 | 21 | 45820448 | -0.113349116 | 3.66E-05 | 0.875640476 |
| APOBEC3B | 22 | rs11600036 | 11 | 81290049 | -0.144626105 | 3.75E-05 | 0.879948067 |
| APOBEC3B | 22 | rs12389354 | X | 116234513 | 0.093400585 | 3.98E-05 | 0.894226918 |
| APOBEC3B | 22 | rs17429397 | 1 | 79951789 | 0.301619525 | 4.01E-05 | 0.894226918 |
| APOBEC3B | 22 | rs73054250 | 1 | 193662371 | 0.197536113 | 4.05E-05 | 0.896458005 |
| APOBEC3B | 22 | rs10042679 | 5 | 179199759 | 0.165922851 | 4.20E-05 | 0.896458005 |
| APOBEC3B | 22 | rs1118150 | 2 | 217415545 | 0.10769048 | 4.23E-05 | 0.896458005 |
| APOBEC3B | 22 | rs858956 | 2 | 50651874 | -0.139908886 | 4.25E-05 | 0.896458005 |
| APOBEC3B | 22 | rs7280173 | 21 | 45841792 | -0.118919664 | 4.30E-05 | 0.901453344 |
| APOBEC3B | 22 | rs11144476 | 9 | 68951931 | 0.123481734 | 4.39E-05 | 0.907207573 |
| APOBEC3B | 22 | rs7737765 | 5 | 16794671 | 0.124193926 | 4.48E-05 | 0.90811064 |
| APOBEC3B | 22 | rs35846970 | 3 | 187453022 | 0.094071479 | 4.74E-05 | 0.912369094 |
| APOBEC3B | 22 | rs112588597 | 7 | 62765576 | 0.33652925 | 4.92E-05 | 0.914201928 |
| APOBEC3B | 22 | rs2120844 | 4 | 142901190 | -0.111606265 | 4.95E-05 | 0.914201928 |
| APOBEC3B | 22 | rs9978740 | 21 | 45748600 | -0.110530363 | 5.06E-05 | 0.914201928 |
| APOBEC3B | 22 | rs188417 | 3 | 139590036 | 0.206532655 | 5.31E-05 | 0.91575718 |
| APOBEC3B | 22 | rs6746221 | 2 | 209290834 | 0.098376603 | 5.34E-05 | 0.91575718 |
| APOBEC3B | 22 | rs716878 | 4 | 142914153 | -0.110803263 | 5.50E-05 | 0.91575718 |
| APOBEC3B | 22 | rs10833249 | 11 | 20119868 | 0.11898825 | 5.60E-05 | 0.918110223 |

**Supplementary Table 2d.** *Trans*-eQTL identified for the *APOBEC3B* in non-involved lung tissue of 408 Italian lung adenocarcinoma patients ($P < 1.0 \times 10^{-4}$), listed in order of *P*-value.

| gene | gene chr | SNP | SNP chr | SNP position | beta | p-value | FDR |
|---|---|---|---|---|---|---|---|
| APOBEC3B | 22 | rs2183596 | 21 | 45755537 | -0.109014316 | 5.65E-05 | 0.918110223 |
| APOBEC3B | 22 | rs139616402 | X | 97776015 | 0.25731924 | 5.87E-05 | 0.918110223 |
| APOBEC3B | 22 | rs7630583 | 3 | 187482581 | -0.091602179 | 6.20E-05 | 0.919360784 |
| APOBEC3B | 22 | rs1381266 | X | 20554574 | 0.107644258 | 6.25E-05 | 0.920613194 |
| APOBEC3B | 22 | rs1381266 | X | 20554574 | 0.107644258 | 6.25E-05 | 0.920613194 |
| APOBEC3B | 22 | rs579775 | 9 | 110109584 | -0.105297211 | 6.34E-05 | 0.924166007 |
| APOBEC3B | 22 | rs3788220 | 21 | 45868078 | -0.10466247 | 6.42E-05 | 0.924166007 |
| APOBEC3B | 22 | rs17284515 | 18 | 71370731 | 0.123742886 | 6.52E-05 | 0.924166007 |
| APOBEC3B | 22 | rs7959966 | 12 | 3799056 | 0.110767328 | 6.53E-05 | 0.924166007 |
| APOBEC3B | 22 | rs3367 | 21 | 45763982 | -0.109592762 | 6.64E-05 | 0.924166007 |
| APOBEC3B | 22 | rs613968 | 11 | 79227617 | -0.122011993 | 6.64E-05 | 0.924166007 |
| APOBEC3B | 22 | rs4980694 | 11 | 69796059 | -0.119137345 | 6.77E-05 | 0.924166007 |
| APOBEC3B | 22 | rs11027861 | 11 | 24385254 | -0.132680254 | 6.83E-05 | 0.924166007 |
| APOBEC3B | 22 | rs1403589 | 15 | 26052770 | 0.157871752 | 6.86E-05 | 0.924166007 |
| APOBEC3B | 22 | rs17004799 | 21 | 45709551 | -0.107759068 | 6.99E-05 | 0.924166007 |
| APOBEC3B | 22 | rs66867540 | 4 | 25029056 | 0.098063795 | 7.06E-05 | 0.924166007 |
| APOBEC3B | 22 | rs76476993 | 7 | 55548382 | 0.260861959 | 7.12E-05 | 0.924166007 |
| APOBEC3B | 22 | rs7682354 | 4 | 130974982 | 0.228596473 | 7.38E-05 | 0.924166007 |
| APOBEC3B | 22 | rs9509982 | 13 | 22044277 | 0.108104081 | 7.50E-05 | 0.924166007 |
| APOBEC3B | 22 | rs10018936 | 4 | 174657671 | -0.098457787 | 7.75E-05 | 0.924166007 |
| APOBEC3B | 22 | rs12511536 | 4 | 174658291 | -0.098457787 | 7.75E-05 | 0.924166007 |
| APOBEC3B | 22 | rs4425356 | 4 | 174661833 | -0.098457787 | 7.75E-05 | 0.924166007 |
| APOBEC3B | 22 | rs5922425 | X | 87337504 | 0.125880669 | 8.04E-05 | 0.924166007 |
| APOBEC3B | 22 | rs1381266 | X | 20554574 | 0.106015098 | 8.07E-05 | 0.924166007 |
| APOBEC3B | 22 | rs1381266 | X | 20554574 | 0.106015098 | 8.07E-05 | 0.924166007 |
| APOBEC3B | 22 | rs2159604 | 12 | 3796249 | 0.108579018 | 8.14E-05 | 0.924166007 |
| APOBEC3B | 22 | rs3788216 | 21 | 45867312 | -0.113044043 | 8.66E-05 | 0.924166007 |
| APOBEC3B | 22 | rs4863838 | 4 | 131019478 | 0.218804322 | 8.75E-05 | 0.924166007 |
| APOBEC3B | 22 | rs11190944 | 10 | 101344089 | -0.098071845 | 9.13E-05 | 0.924166007 |
| APOBEC3B | 22 | rs4908343 | 1 | 27605187 | -0.104177071 | 9.26E-05 | 0.924166007 |
| APOBEC3B | 22 | rs75188236 | 15 | 68607384 | -0.301882995 | 9.30E-05 | 0.924166007 |
| APOBEC3B | 22 | rs480859 | 11 | 132726883 | 0.099983343 | 9.38E-05 | 0.924166007 |
| APOBEC3B | 22 | rs9935088 | 16 | 78945618 | 0.14140165 | 9.56E-05 | 0.924166007 |
| APOBEC3B | 22 | rs1441413 | 4 | 142902686 | -0.099754738 | 9.73E-05 | 0.924166007 |
| APOBEC3B | 22 | rs62559212 | 9 | 75897049 | -0.138936056 | 9.92E-05 | 0.924166007 |

**Supplementary Table 2e.** *Trans*-eQTL identified for the *APOBEC3C* in non-involved lung tissue of 408 Italian lung adenocarcinoma patients ($P < 1.0 \times 10^{-4}$), listed in order of *P*-value.

| gene | gene chr | SNP | SNP chr | SNP position | beta | p-value | FDR |
|---|---|---|---|---|---|---|---|
| APOBEC3C | 22 | rs1496248 | 11 | 133682217 | -0.133222854 | 1.20E-06 | 0.667197673 |
| APOBEC3C | 22 | rs203849 | 1 | 167880176 | -0.111335839 | 3.76E-06 | 0.84453778 |
| APOBEC3C | 22 | rs10072177 | 5 | 26927395 | 0.225141481 | 4.92E-06 | 0.84453778 |
| APOBEC3C | 22 | rs10064442 | 5 | 26931897 | 0.225141481 | 4.92E-06 | 0.84453778 |
| APOBEC3C | 22 | rs4740196 | 9 | 130577940 | -0.181115672 | 6.76E-06 | 0.84453778 |
| APOBEC3C | 22 | rs3087943 | 6 | 24650533 | 0.138136234 | 1.10E-05 | 0.84453778 |
| APOBEC3C | 22 | rs74688188 | 12 | 32984196 | 0.25985477 | 1.13E-05 | 0.84453778 |
| APOBEC3C | 22 | rs2242342 | 11 | 92913088 | 0.139968824 | 1.22E-05 | 0.84453778 |
| APOBEC3C | 22 | rs10732252 | 11 | 133661733 | -0.127364669 | 1.36E-05 | 0.848481351 |
| APOBEC3C | 22 | rs2832658 | 21 | 30217096 | 0.117219477 | 1.53E-05 | 0.848481351 |
| APOBEC3C | 22 | rs28513267 | 7 | 156608761 | -0.188577837 | 1.98E-05 | 0.863488543 |
| APOBEC3C | 22 | rs2353005 | 19 | 49869505 | -0.152169822 | 1.99E-05 | 0.863488543 |
| APOBEC3C | 22 | rs7278769 | 21 | 30221383 | 0.124190298 | 2.14E-05 | 0.863488543 |
| APOBEC3C | 22 | rs12495745 | 3 | 20367532 | -0.17808296 | 2.20E-05 | 0.863488543 |
| APOBEC3C | 22 | rs76588334 | 8 | 78110813 | 0.188054571 | 2.46E-05 | 0.863488543 |
| APOBEC3C | 22 | rs9376611 | 6 | 141371496 | 0.10619474 | 2.61E-05 | 0.863488543 |
| APOBEC3C | 22 | rs870643 | 20 | 61731779 | 0.149302701 | 2.63E-05 | 0.863488543 |
| APOBEC3C | 22 | rs201066567 | 21 | 30249726 | 0.103442162 | 3.04E-05 | 0.875040765 |
| APOBEC3C | 22 | rs201768560 | 10 | 24315764 | 0.105810124 | 3.04E-05 | 0.875040765 |
| APOBEC3C | 22 | rs11583747 | 1 | 195526131 | 0.115772645 | 3.07E-05 | 0.875040765 |
| APOBEC3C | 22 | rs882308 | 4 | 114025712 | -0.140301407 | 3.09E-05 | 0.875040765 |
| APOBEC3C | 22 | rs7031913 | 9 | 24112126 | 0.107958255 | 3.15E-05 | 0.875040765 |
| APOBEC3C | 22 | rs17677852 | 4 | 114035524 | -0.139392534 | 3.36E-05 | 0.875040765 |
| APOBEC3C | 22 | rs11020104 | 11 | 92929571 | 0.131543516 | 3.50E-05 | 0.875040765 |
| APOBEC3C | 22 | rs201074054 | 4 | 114024126 | -0.137569973 | 3.52E-05 | 0.875040765 |
| APOBEC3C | 22 | rs12644731 | 4 | 114087335 | -0.139644721 | 3.77E-05 | 0.879948067 |
| APOBEC3C | 22 | rs7801996 | 7 | 156613217 | -0.160753945 | 3.91E-05 | 0.89286517 |
| APOBEC3C | 22 | rs79124404 | 20 | 50436002 | 0.316349455 | 4.01E-05 | 0.894226918 |
| APOBEC3C | 22 | rs9376609 | 6 | 141350595 | 0.102974057 | 4.68E-05 | 0.912369094 |
| APOBEC3C | 22 | rs11791133 | 9 | 20303920 | -0.102819997 | 4.76E-05 | 0.912369094 |
| APOBEC3C | 22 | rs1023369 | 21 | 30245254 | 0.101093815 | 4.83E-05 | 0.914201928 |
| APOBEC3C | 22 | rs2974277 | 8 | 87102930 | -0.124574833 | 4.91E-05 | 0.914201928 |
| APOBEC3C | 22 | rs2374223 | 4 | 114021753 | -0.133296746 | 5.14E-05 | 0.914201928 |
| APOBEC3C | 22 | rs2012484 | 21 | 30244148 | 0.100347553 | 5.51E-05 | 0.91575718 |
| APOBEC3C | 22 | rs7689486 | 4 | 7333853 | -0.114791056 | 5.73E-05 | 0.918110223 |
| APOBEC3C | 22 | rs1452991 | 6 | 141152226 | 0.099786769 | 5.82E-05 | 0.918110223 |
| APOBEC3C | 22 | rs6020495 | 20 | 50429241 | 0.303142295 | 5.86E-05 | 0.918110223 |
| APOBEC3C | 22 | rs4780315 | 16 | 10623710 | 0.107000965 | 6.00E-05 | 0.918110223 |
| APOBEC3C | 22 | rs12380144 | 9 | 24083441 | 0.111040297 | 6.22E-05 | 0.91987342 |
| APOBEC3C | 22 | rs1037442 | 6 | 141275602 | 0.099080622 | 6.23E-05 | 0.920157798 |
| APOBEC3C | 22 | rs12038043 | 1 | 195536987 | 0.150009228 | 6.45E-05 | 0.924166007 |
| APOBEC3C | 22 | rs7927604 | 11 | 92896433 | 0.109023094 | 6.56E-05 | 0.924166007 |
| APOBEC3C | 22 | rs505480 | 8 | 13021334 | -0.258158827 | 6.62E-05 | 0.924166007 |
| APOBEC3C | 22 | rs10047182 | 1 | 4474261 | -0.144315802 | 6.69E-05 | 0.924166007 |
| APOBEC3C | 22 | rs28874125 | 4 | 114070032 | -0.132432152 | 6.90E-05 | 0.924166007 |
| APOBEC3C | 22 | rs1721282 | 2 | 224440872 | -0.104298168 | 7.70E-05 | 0.924166007 |
| APOBEC3C | 22 | rs17204200 | 20 | 61284434 | -0.149024385 | 7.78E-05 | 0.924166007 |
| APOBEC3C | 22 | rs62022281 | 15 | 42353976 | -0.146932407 | 7.84E-05 | 0.924166007 |

**Supplementary Table 2e.** *Trans*-eQTL identified for the *APOBEC3C* in non-involved lung tissue of 408 Italian lung adenocarcinoma patients ($P < 1.0 \times 10^{-4}$), listed in order of *P*-value.

| gene | gene chr | SNP | SNP chr | SNP position | beta | p-value | FDR |
|---|---|---|---|---|---|---|---|
| APOBEC3C | 22 | rs13097492 | 3 | 197427636 | -0.100200754 | 8.20E-05 | 0.924166007 |
| APOBEC3C | 22 | rs4326079 | 4 | 114021047 | -0.131951052 | 8.27E-05 | 0.924166007 |
| APOBEC3C | 22 | rs12769905 | 10 | 119977002 | 0.108095699 | 8.33E-05 | 0.924166007 |
| APOBEC3C | 22 | rs13229349 | 7 | 158731839 | 0.131879456 | 8.41E-05 | 0.924166007 |
| APOBEC3C | 22 | rs3763406 | 7 | 158732072 | 0.131879456 | 8.41E-05 | 0.924166007 |
| APOBEC3C | 22 | rs3763410 | 7 | 158732370 | 0.131879456 | 8.41E-05 | 0.924166007 |
| APOBEC3C | 22 | rs1061739 | 7 | 158733663 | 0.131879456 | 8.41E-05 | 0.924166007 |
| APOBEC3C | 22 | rs10520529 | 4 | 182394766 | -0.10318458 | 8.42E-05 | 0.924166007 |
| APOBEC3C | 22 | rs17798635 | 6 | 155079485 | 0.240130084 | 8.55E-05 | 0.924166007 |
| APOBEC3C | 22 | rs78967251 | 20 | 50446298 | 0.286029084 | 8.59E-05 | 0.924166007 |
| APOBEC3C | 22 | rs16903972 | 5 | 38442014 | 0.352237981 | 8.62E-05 | 0.924166007 |
| APOBEC3C | 22 | rs4833439 | 4 | 114017322 | -0.131111743 | 8.93E-05 | 0.924166007 |
| APOBEC3C | 22 | rs13112787 | 4 | 114037387 | -0.133452133 | 9.08E-05 | 0.924166007 |
| APOBEC3C | 22 | rs6533743 | 4 | 114036049 | -0.130198748 | 9.19E-05 | 0.924166007 |
| APOBEC3C | 22 | rs6707963 | 2 | 82255386 | -0.120418651 | 9.21E-05 | 0.924166007 |
| APOBEC3C | 22 | rs57598153 | 7 | 156617411 | -0.174030619 | 9.26E-05 | 0.924166007 |
| APOBEC3C | 22 | rs9403295 | 6 | 141384021 | 0.096577253 | 9.51E-05 | 0.924166007 |
| APOBEC3C | 22 | rs11046811 | 12 | 23125015 | -0.113463696 | 9.58E-05 | 0.924166007 |

**Supplementary Table 2f.** *Trans*-eQTL identified for the *APOBEC3D* in non-involved lung tissue of 408 Italian lung adenocarcinoma patients ($P < 1.0 \times 10^{-4}$), listed in order of *P*-value.

| gene | gene chr | SNP | SNP chr | SNP position | beta | p-value | FDR |
|---|---|---|---|---|---|---|---|
| APOBEC3D | 22 | rs1454856 | 1 | 85525400 | -0.108471586 | 8.86E-07 | 0.65901749 |
| APOBEC3D | 22 | rs1424106 | 16 | 79182896 | -0.104727869 | 1.72E-06 | 0.767785475 |
| APOBEC3D | 22 | rs719830 | 6 | 72722245 | 0.142545343 | 3.09E-06 | 0.84453778 |
| APOBEC3D | 22 | rs55936668 | 22 | 36146759 | 0.189739209 | 4.49E-06 | 0.84453778 |
| APOBEC3D | 22 | rs11161636 | 1 | 85544565 | -0.104188538 | 4.77E-06 | 0.84453778 |
| APOBEC3D | 22 | rs17730134 | 16 | 79185763 | 0.10227855 | 4.93E-06 | 0.84453778 |
| APOBEC3D | 22 | rs12200468 | 6 | 72642142 | 0.136400131 | 6.01E-06 | 0.84453778 |
| APOBEC3D | 22 | rs10943051 | 6 | 72664188 | 0.137727724 | 7.53E-06 | 0.84453778 |
| APOBEC3D | 22 | rs17332991 | 5 | 60883533 | -0.155226652 | 7.88E-06 | 0.84453778 |
| APOBEC3D | 22 | rs77647293 | 5 | 40619770 | 0.307617784 | 1.03E-05 | 0.84453778 |
| APOBEC3D | 22 | rs11709953 | 3 | 34840944 | 0.128264217 | 1.22E-05 | 0.84453778 |
| APOBEC3D | 22 | rs2906721 | 7 | 101823718 | 0.120926558 | 1.44E-05 | 0.848481351 |
| APOBEC3D | 22 | rs76093199 | 4 | 188978777 | 0.271993408 | 1.59E-05 | 0.853417593 |
| APOBEC3D | 22 | rs79458507 | 3 | 10667662 | 0.157778218 | 1.60E-05 | 0.853417593 |
| APOBEC3D | 22 | rs74735032 | 1 | 27277650 | 0.235793579 | 1.77E-05 | 0.863488543 |
| APOBEC3D | 22 | rs1336197 | 10 | 54737899 | 0.099376245 | 1.78E-05 | 0.863488543 |
| APOBEC3D | 22 | rs62303602 | 4 | 81467404 | 0.135939628 | 1.85E-05 | 0.863488543 |
| APOBEC3D | 22 | rs7752777 | 6 | 23202625 | -0.092238705 | 1.96E-05 | 0.863488543 |
| APOBEC3D | 22 | rs7540868 | 1 | 70861679 | -0.095801909 | 1.97E-05 | 0.863488543 |
| APOBEC3D | 22 | rs7530345 | 1 | 70861409 | -0.095795532 | 1.97E-05 | 0.863488543 |
| APOBEC3D | 22 | rs12205374 | 6 | 72727632 | 0.134618595 | 2.18E-05 | 0.863488543 |
| APOBEC3D | 22 | rs8050297 | 16 | 53020579 | -0.121380578 | 2.30E-05 | 0.863488543 |
| APOBEC3D | 22 | rs12212268 | 6 | 72736359 | 0.124804582 | 2.35E-05 | 0.863488543 |
| APOBEC3D | 22 | rs750459 | 5 | 171514968 | -0.101261212 | 2.55E-05 | 0.863488543 |
| APOBEC3D | 22 | rs13278849 | 8 | 26857357 | 0.101051278 | 2.66E-05 | 0.863488543 |
| APOBEC3D | 22 | rs2644295 | 7 | 2820583 | 0.092438715 | 2.66E-05 | 0.863488543 |
| APOBEC3D | 22 | rs10069885 | 5 | 14064971 | -0.095628814 | 2.82E-05 | 0.875040765 |
| APOBEC3D | 22 | rs9567446 | 13 | 32226289 | 0.150190977 | 3.07E-05 | 0.875040765 |
| APOBEC3D | 22 | rs1108663 | 16 | 79187995 | -0.092806005 | 3.12E-05 | 0.875040765 |
| APOBEC3D | 22 | rs2941940 | 16 | 78544162 | 0.09680439 | 3.13E-05 | 0.875040765 |
| APOBEC3D | 22 | rs9906006 | 17 | 13718644 | -0.091824729 | 3.24E-05 | 0.875040765 |
| APOBEC3D | 22 | rs12483846 | 22 | 30724727 | 0.174782116 | 3.28E-05 | 0.875040765 |
| APOBEC3D | 22 | rs5753290 | 22 | 30725659 | 0.174782116 | 3.28E-05 | 0.875040765 |
| APOBEC3D | 22 | rs34667146 | 13 | 99820429 | 0.107694845 | 3.61E-05 | 0.875040765 |
| APOBEC3D | 22 | rs62411975 | 6 | 72617245 | 0.12506865 | 3.62E-05 | 0.875040765 |
| APOBEC3D | 22 | rs116821300 | 5 | 40345272 | 0.287001084 | 3.86E-05 | 0.890332882 |
| APOBEC3D | 22 | rs7802881 | 7 | 68856585 | 0.134960197 | 4.45E-05 | 0.90811064 |
| APOBEC3D | 22 | rs117892826 | 15 | 55962224 | 0.248225041 | 4.79E-05 | 0.913006272 |
| APOBEC3D | 22 | rs1691327 | 21 | 17572543 | 0.174004598 | 4.86E-05 | 0.914201928 |
| APOBEC3D | 22 | rs6099648 | 20 | 57482875 | -0.091463461 | 4.96E-05 | 0.914201928 |
| APOBEC3D | 22 | rs11078173 | 17 | 13672839 | -0.090459393 | 5.00E-05 | 0.914201928 |
| APOBEC3D | 22 | rs76546184 | 15 | 62821752 | -0.284140972 | 5.03E-05 | 0.914201928 |
| APOBEC3D | 22 | rs10846013 | 12 | 14511661 | 0.096597988 | 5.26E-05 | 0.91575718 |
| APOBEC3D | 22 | rs4753437 | 11 | 93200725 | 0.109490684 | 5.50E-05 | 0.91575718 |
| APOBEC3D | 22 | rs56342921 | 16 | 78577374 | 0.120558181 | 5.89E-05 | 0.918110223 |
| APOBEC3D | 22 | rs7777484 | 7 | 2774637 | 0.087621715 | 6.04E-05 | 0.918110223 |
| APOBEC3D | 22 | rs2527684 | 7 | 2786288 | 0.088940434 | 6.10E-05 | 0.918110223 |
| APOBEC3D | 22 | rs2293837 | 14 | 79841514 | 0.092702074 | 6.34E-05 | 0.924166007 |

**Supplementary Table 2f.** *Trans*-eQTL identified for the *APOBEC3D* in non-involved lung tissue of 408 Italian lung adenocarcinoma patients ($P < 1.0 \times 10^{-4}$), listed in order of *P*-value.

| gene | gene chr | SNP | SNP chr | SNP position | beta | p-value | FDR |
|---|---|---|---|---|---|---|---|
| APOBEC3D | 22 | rs113587473 | X | 6206125 | 0.06777677 | 6.58E-05 | 0.924166007 |
| APOBEC3D | 22 | rs10845808 | 12 | 13589558 | 0.120193131 | 6.60E-05 | 0.924166007 |
| APOBEC3D | 22 | rs73247924 | 4 | 26510638 | 0.178973686 | 6.72E-05 | 0.924166007 |
| APOBEC3D | 22 | rs9431929 | 1 | 231112867 | 0.265097016 | 7.33E-05 | 0.924166007 |
| APOBEC3D | 22 | rs706503 | 12 | 25852812 | 0.085814309 | 7.36E-05 | 0.924166007 |
| APOBEC3D | 22 | rs11924964 | 3 | 139343002 | 0.12924716 | 7.42E-05 | 0.924166007 |
| APOBEC3D | 22 | rs2729923 | 11 | 19418908 | 0.08877859 | 7.42E-05 | 0.924166007 |
| APOBEC3D | 22 | rs4515943 | 11 | 19419975 | 0.08877859 | 7.42E-05 | 0.924166007 |
| APOBEC3D | 22 | rs11783456 | 8 | 74005936 | -0.094181438 | 7.77E-05 | 0.924166007 |
| APOBEC3D | 22 | rs72709232 | 9 | 29475181 | 0.214890044 | 7.80E-05 | 0.924166007 |
| APOBEC3D | 22 | rs1185614 | 12 | 25848138 | 0.085119403 | 8.01E-05 | 0.924166007 |
| APOBEC3D | 22 | rs62235878 | 22 | 30690485 | 0.170620207 | 8.03E-05 | 0.924166007 |
| APOBEC3D | 22 | rs1636255 | 7 | 2853170 | 0.094438203 | 8.33E-05 | 0.924166007 |
| APOBEC3D | 22 | rs79348617 | 1 | 183029118 | 0.22390514 | 8.60E-05 | 0.924166007 |
| APOBEC3D | 22 | rs17673749 | 10 | 73850659 | 0.247627975 | 8.64E-05 | 0.924166007 |
| APOBEC3D | 22 | rs77285730 | 21 | 17579089 | 0.197080063 | 8.87E-05 | 0.924166007 |
| APOBEC3D | 22 | rs11110252 | 12 | 100129718 | 0.162915118 | 9.10E-05 | 0.924166007 |
| APOBEC3D | 22 | rs11625881 | 14 | 79837823 | 0.090634561 | 9.26E-05 | 0.924166007 |
| APOBEC3D | 22 | rs7864125 | 9 | 29500533 | 0.114997709 | 9.29E-05 | 0.924166007 |
| APOBEC3D | 22 | rs35215795 | 5 | 133357758 | -0.13131257 | 9.34E-05 | 0.924166007 |
| APOBEC3D | 22 | rs17166628 | 5 | 133354540 | -0.12468497 | 9.46E-05 | 0.924166007 |
| APOBEC3D | 22 | rs17634230 | 15 | 94791216 | 0.152671223 | 9.48E-05 | 0.924166007 |
| APOBEC3D | 22 | rs990824 | 12 | 13575007 | 0.188654977 | 9.74E-05 | 0.924166007 |
| APOBEC3D | 22 | rs1150476 | 5 | 5959971 | -0.135903899 | 9.81E-05 | 0.924166007 |
| APOBEC3D | 22 | rs10504646 | 8 | 77158158 | -0.188722082 | 9.81E-05 | 0.924166007 |
| APOBEC3D | 22 | rs11110259 | 12 | 100151891 | 0.162197202 | 9.82E-05 | 0.924166007 |

**Supplementary Table 2g.** *Trans*-eQTL identified for the *APOBEC3F* in non-involved lung tissue of 408 Italian lung adenocarcinoma patients ($P < 1.0 \times 10^{-4}$), listed in order of *P*-value.

| gene | gene chr | SNP | SNP chr | SNP position | beta | p-value | FDR |
|---|---|---|---|---|---|---|---|
| APOBEC3F | 22 | rs199678960 | 16 | 3990775 | -0.124886977 | 6.02E-07 | 0.65901749 |
| APOBEC3F | 22 | rs488035 | 19 | 56081053 | 0.218417307 | 9.74E-07 | 0.65901749 |
| APOBEC3F | 22 | rs2444217 | 16 | 3988386 | -0.120139922 | 1.36E-06 | 0.71466926 |
| APOBEC3F | 22 | rs3949938 | 14 | 64924765 | 0.199741443 | 3.87E-06 | 0.84453778 |
| APOBEC3F | 22 | rs28375296 | 15 | 86488955 | -0.155890146 | 6.52E-06 | 0.84453778 |
| APOBEC3F | 22 | rs142021956 | 3 | 11884291 | 0.280537067 | 1.44E-05 | 0.848481351 |
| APOBEC3F | 22 | rs4583504 | 2 | 1172485 | -0.137271817 | 1.50E-05 | 0.848481351 |
| APOBEC3F | 22 | rs4971407 | 2 | 1188160 | -0.136688025 | 1.75E-05 | 0.863488543 |
| APOBEC3F | 22 | rs117981161 | 9 | 35928074 | 0.386656494 | 1.79E-05 | 0.863488543 |
| APOBEC3F | 22 | rs34550784 | 5 | 80949849 | 0.123263018 | 1.80E-05 | 0.863488543 |
| APOBEC3F | 22 | rs2124432 | 2 | 128426592 | -0.117347595 | 1.88E-05 | 0.863488543 |
| APOBEC3F | 22 | rs2124432 | 2 | 128426592 | -0.117347595 | 1.88E-05 | 0.863488543 |
| APOBEC3F | 22 | rs2124432 | 2 | 128426592 | -0.117347595 | 1.88E-05 | 0.863488543 |
| APOBEC3F | 22 | rs2124432 | 2 | 128426592 | -0.117347595 | 1.88E-05 | 0.863488543 |
| APOBEC3F | 22 | rs10815124 | 9 | 4933391 | -0.111258945 | 2.18E-05 | 0.863488543 |
| APOBEC3F | 22 | rs11588699 | 1 | 47524527 | 0.181745223 | 2.25E-05 | 0.863488543 |
| APOBEC3F | 22 | rs11242536 | 5 | 104774418 | 0.110144306 | 2.45E-05 | 0.863488543 |
| APOBEC3F | 22 | rs56202519 | 5 | 80943930 | 0.123483166 | 2.56E-05 | 0.863488543 |
| APOBEC3F | 22 | rs10974859 | 9 | 4930487 | -0.109358404 | 2.89E-05 | 0.875040765 |
| APOBEC3F | 22 | rs72904183 | 18 | 7261259 | -0.174954458 | 2.93E-05 | 0.875040765 |
| APOBEC3F | 22 | rs62192926 | 2 | 233240091 | -0.111759137 | 3.16E-05 | 0.875040765 |
| APOBEC3F | 22 | rs10873179 | 14 | 64960345 | 0.157836113 | 3.19E-05 | 0.875040765 |
| APOBEC3F | 22 | rs57432155 | 6 | 82063851 | 0.138084307 | 3.51E-05 | 0.875040765 |
| APOBEC3F | 22 | rs8130517 | 21 | 36298159 | -0.107164347 | 3.51E-05 | 0.875040765 |
| APOBEC3F | 22 | rs6634815 | X | 131221163 | 0.11812761 | 3.61E-05 | 0.875040765 |
| APOBEC3F | 22 | rs7149828 | 14 | 94629284 | -0.113310095 | 3.62E-05 | 0.875040765 |
| APOBEC3F | 22 | rs4820102 | 22 | 33355076 | 0.185370837 | 3.66E-05 | 0.875640476 |
| APOBEC3F | 22 | rs67802170 | 1 | 239574707 | 0.224426942 | 3.67E-05 | 0.875640476 |
| APOBEC3F | 22 | rs79123033 | 7 | 54049501 | 0.268626523 | 3.75E-05 | 0.879948067 |
| APOBEC3F | 22 | rs11875530 | 18 | 57078925 | -0.167690321 | 3.76E-05 | 0.879948067 |
| APOBEC3F | 22 | rs57703465 | 10 | 69881312 | 0.22230576 | 3.81E-05 | 0.879948067 |
| APOBEC3F | 22 | rs455021 | 3 | 11904333 | 0.137951151 | 4.09E-05 | 0.896458005 |
| APOBEC3F | 22 | rs10047182 | 1 | 4474261 | -0.149403287 | 4.10E-05 | 0.896458005 |
| APOBEC3F | 22 | rs34499282 | 16 | 5405669 | 0.14356251 | 4.36E-05 | 0.905224842 |
| APOBEC3F | 22 | rs10754875 | 1 | 14967535 | 0.116032798 | 4.67E-05 | 0.912369094 |
| APOBEC3F | 22 | rs1540493 | 2 | 128427292 | -0.178492722 | 5.00E-05 | 0.914201928 |
| APOBEC3F | 22 | rs4516307 | 18 | 6759891 | 0.112414462 | 5.01E-05 | 0.914201928 |
| APOBEC3F | 22 | rs72705269 | 1 | 109785565 | 0.145938327 | 5.15E-05 | 0.914201928 |
| APOBEC3F | 22 | rs113926657 | 10 | 131691681 | -0.190386388 | 5.16E-05 | 0.914201928 |
| APOBEC3F | 22 | rs34700798 | 1 | 230397087 | 0.202458625 | 5.37E-05 | 0.91575718 |
| APOBEC3F | 22 | rs4704855 | 5 | 157201237 | -0.146376344 | 5.64E-05 | 0.918110223 |
| APOBEC3F | 22 | rs4798491 | 18 | 6758922 | 0.111662737 | 5.65E-05 | 0.918110223 |
| APOBEC3F | 22 | rs4821143 | 22 | 33355081 | 0.181077872 | 6.11E-05 | 0.918110223 |
| APOBEC3F | 22 | rs12677662 | 8 | 133986847 | -0.145836944 | 6.32E-05 | 0.924166007 |
| APOBEC3F | 22 | rs29551 | 5 | 133371835 | 0.124137383 | 6.52E-05 | 0.924166007 |
| APOBEC3F | 22 | rs7762168 | 6 | 133202641 | -0.107459472 | 6.69E-05 | 0.924166007 |
| APOBEC3F | 22 | rs7172340 | 15 | 68288463 | 0.340447515 | 6.78E-05 | 0.924166007 |
| APOBEC3F | 22 | rs2601777 | 16 | 3985067 | -0.101086677 | 6.88E-05 | 0.924166007 |

**Supplementary Table 2g.** *Trans*-eQTL identified for the *APOBEC3F* in non-involved lung tissue of 408 Italian lung adenocarcinoma patients ($P < 1.0 \times 10^{-4}$), listed in order of *P*-value.

| gene | gene chr | SNP | SNP chr | SNP position | beta | p-value | FDR |
|---|---|---|---|---|---|---|---|
| APOBEC3F | 22 | rs809367 | 10 | 87982049 | 0.151334534 | 6.97E-05 | 0.924166007 |
| APOBEC3F | 22 | rs884656 | 6 | 82057693 | 0.136167341 | 7.16E-05 | 0.924166007 |
| APOBEC3F | 22 | rs17102318 | 14 | 64914853 | 0.180619868 | 7.19E-05 | 0.924166007 |
| APOBEC3F | 22 | rs75157479 | 4 | 108351341 | 0.329386359 | 7.39E-05 | 0.924166007 |
| APOBEC3F | 22 | rs34782392 | 4 | 182837006 | -0.100566081 | 7.63E-05 | 0.924166007 |
| APOBEC3F | 22 | rs56363627 | 10 | 11019327 | -0.133460743 | 7.70E-05 | 0.924166007 |
| APOBEC3F | 22 | rs3864338 | 7 | 157700887 | 0.25766933 | 7.71E-05 | 0.924166007 |
| APOBEC3F | 22 | rs12679902 | 8 | 89845761 | 0.118111279 | 8.06E-05 | 0.924166007 |
| APOBEC3F | 22 | rs9419619 | 10 | 131686021 | -0.186069949 | 8.49E-05 | 0.924166007 |
| APOBEC3F | 22 | rs56143914 | 11 | 11113017 | 0.173755319 | 8.55E-05 | 0.924166007 |
| APOBEC3F | 22 | rs58132943 | 11 | 11113219 | 0.173755319 | 8.55E-05 | 0.924166007 |
| APOBEC3F | 22 | rs6770843 | 3 | 185898873 | 0.105909591 | 8.81E-05 | 0.924166007 |
| APOBEC3F | 22 | rs16936996 | 10 | 36425900 | 0.311419552 | 8.88E-05 | 0.924166007 |
| APOBEC3F | 22 | rs75778065 | 6 | 41468219 | 0.246745445 | 8.92E-05 | 0.924166007 |
| APOBEC3F | 22 | rs113554045 | 8 | 107073456 | -0.09764344 | 9.14E-05 | 0.924166007 |
| APOBEC3F | 22 | rs78935380 | 11 | 11100832 | 0.171231518 | 9.17E-05 | 0.924166007 |
| APOBEC3F | 22 | rs4686399 | 3 | 185900478 | 0.106928141 | 9.24E-05 | 0.924166007 |
| APOBEC3F | 22 | rs4726277 | 7 | 153336681 | 0.108271683 | 9.79E-05 | 0.924166007 |
| APOBEC3F | 22 | rs77003492 | 17 | 73735432 | 0.295518353 | 9.94E-05 | 0.924166007 |

**Supplementary Table 2h.** *Trans*-eQTL identified for the *APOBEC3G* in non-involved lung tissue of 408 Italian lung adenocarcinoma patients ($P < 1.0 \times 10^{-4}$), listed in order of *P*-value.

| gene | gene chr | SNP | SNP chr | SNP position | beta | p-value | FDR |
|---|---|---|---|---|---|---|---|
| APOBEC3G | 22 | rs77028530 | 3 | 184631800 | 0.374333327 | 5.14E-07 | 0.65901749 |
| APOBEC3G | 22 | rs66832033 | 1 | 81558380 | 0.235953193 | 3.77E-06 | 0.84453778 |
| APOBEC3G | 22 | rs669918 | 11 | 97651098 | -0.209405774 | 4.41E-06 | 0.84453778 |
| APOBEC3G | 22 | rs1560834 | 2 | 173347944 | 0.153388645 | 4.76E-06 | 0.84453778 |
| APOBEC3G | 22 | rs7796629 | 7 | 17635469 | -0.218344418 | 6.27E-06 | 0.84453778 |
| APOBEC3G | 22 | rs3923767 | 2 | 238890906 | -0.147910427 | 8.45E-06 | 0.84453778 |
| APOBEC3G | 22 | rs10484047 | 14 | 94582891 | 0.230219963 | 9.00E-06 | 0.84453778 |
| APOBEC3G | 22 | rs17693059 | 2 | 173342529 | 0.213930775 | 9.17E-06 | 0.84453778 |
| APOBEC3G | 22 | rs11947432 | 4 | 131269925 | 0.484360665 | 1.10E-05 | 0.84453778 |
| APOBEC3G | 22 | rs12598790 | 16 | 66128551 | 0.470843572 | 1.12E-05 | 0.84453778 |
| APOBEC3G | 22 | rs2514726 | 8 | 118005646 | 0.361901512 | 1.18E-05 | 0.84453778 |
| APOBEC3G | 22 | rs61772303 | 1 | 14812872 | 0.35489717 | 1.99E-05 | 0.863488543 |
| APOBEC3G | 22 | rs814526 | 19 | 40502142 | 0.141562782 | 2.20E-05 | 0.863488543 |
| APOBEC3G | 22 | rs12170237 | 22 | 36963448 | 0.337311425 | 2.23E-05 | 0.863488543 |
| APOBEC3G | 22 | rs11590793 | 1 | 14801369 | 0.340161218 | 2.51E-05 | 0.863488543 |
| APOBEC3G | 22 | rs80231304 | 6 | 6313254 | -0.207570569 | 2.64E-05 | 0.863488543 |
| APOBEC3G | 22 | rs17727707 | 12 | 7695606 | -0.377330416 | 2.69E-05 | 0.868532829 |
| APOBEC3G | 22 | rs3826575 | 18 | 80108244 | 0.172068476 | 2.82E-05 | 0.875040765 |
| APOBEC3G | 22 | rs9643022 | 8 | 105752523 | 0.340822034 | 2.94E-05 | 0.875040765 |
| APOBEC3G | 22 | rs76796320 | 5 | 116394200 | 0.381687167 | 3.00E-05 | 0.875040765 |
| APOBEC3G | 22 | rs1383543 | 15 | 100063068 | 0.159671894 | 3.05E-05 | 0.875040765 |
| APOBEC3G | 22 | rs61733121 | 8 | 48729798 | -0.418740965 | 3.11E-05 | 0.875040765 |
| APOBEC3G | 22 | rs2200442 | 16 | 77019881 | -0.13967843 | 3.11E-05 | 0.875040765 |
| APOBEC3G | 22 | rs62313477 | 4 | 117301840 | 0.327801075 | 3.19E-05 | 0.875040765 |
| APOBEC3G | 22 | rs10250001 | 7 | 157970118 | -0.144809374 | 3.35E-05 | 0.875040765 |
| APOBEC3G | 22 | rs11777456 | 8 | 102085378 | 0.150934221 | 3.43E-05 | 0.875040765 |
| APOBEC3G | 22 | rs2986753 | 1 | 6470375 | -0.188847549 | 3.48E-05 | 0.875040765 |
| APOBEC3G | 22 | rs56140076 | 20 | 58393505 | 0.297625524 | 3.53E-05 | 0.875040765 |
| APOBEC3G | 22 | rs17544336 | 20 | 58432538 | 0.297625524 | 3.53E-05 | 0.875040765 |
| APOBEC3G | 22 | rs199923819 | 20 | 58444435 | 0.297625524 | 3.53E-05 | 0.875040765 |
| APOBEC3G | 22 | rs12783734 | 10 | 5363302 | -0.1840232 | 3.75E-05 | 0.879948067 |
| APOBEC3G | 22 | rs7781652 | 7 | 17663792 | -0.147844688 | 3.81E-05 | 0.879948067 |
| APOBEC3G | 22 | rs6098105 | 20 | 54606388 | -0.201187325 | 3.90E-05 | 0.89286517 |
| APOBEC3G | 22 | rs16869054 | 8 | 102080282 | 0.148740315 | 4.16E-05 | 0.896458005 |
| APOBEC3G | 22 | rs12445265 | 16 | 4874159 | -0.170778227 | 4.32E-05 | 0.901579715 |
| APOBEC3G | 22 | rs4484508 | 6 | 66937497 | -0.148023563 | 4.52E-05 | 0.909284759 |
| APOBEC3G | 22 | rs11018197 | 10 | 127553144 | -0.308855966 | 4.57E-05 | 0.909284759 |
| APOBEC3G | 22 | rs4481415 | 6 | 66937020 | -0.147981563 | 4.57E-05 | 0.909284759 |
| APOBEC3G | 22 | rs7407477 | 18 | 80142946 | 0.167595378 | 4.71E-05 | 0.912369094 |
| APOBEC3G | 22 | rs12906090 | 15 | 46897199 | -0.31595055 | 5.08E-05 | 0.914201928 |
| APOBEC3G | 22 | rs12506110 | 4 | 100663327 | 0.13638305 | 5.45E-05 | 0.91575718 |
| APOBEC3G | 22 | rs12213790 | 6 | 88192845 | 0.136371533 | 5.79E-05 | 0.918110223 |
| APOBEC3G | 22 | rs10432142 | 18 | 7199533 | -0.252463802 | 6.09E-05 | 0.918110223 |
| APOBEC3G | 22 | rs17865908 | 4 | 117330264 | 0.322585192 | 6.40E-05 | 0.924166007 |
| APOBEC3G | 22 | rs1994010 | 15 | 100062266 | 0.153193326 | 6.43E-05 | 0.924166007 |
| APOBEC3G | 22 | rs12133065 | 1 | 70428194 | 0.16131414 | 6.44E-05 | 0.924166007 |
| APOBEC3G | 22 | rs9552726 | 13 | 22824961 | 0.158253182 | 6.56E-05 | 0.924166007 |
| APOBEC3G | 22 | rs5985 | 6 | 6318562 | -0.175732243 | 6.65E-05 | 0.924166007 |

**Supplementary Table 2h.** *Trans*-eQTL identified for the *APOBEC3G* in non-involved lung tissue of 408 Italian lung adenocarcinoma patients ($P < 1.0 \times 10^{-4}$), listed in order of *P*-value.

| gene | gene chr | SNP | SNP chr | SNP position | beta | p-value | FDR |
|---|---|---|---|---|---|---|---|
| APOBEC3G | 22 | rs2156153 | 18 | 80115768 | 0.16414261 | 6.67E-05 | 0.924166007 |
| APOBEC3G | 22 | rs2415847 | 14 | 44221873 | 0.142736562 | 6.86E-05 | 0.924166007 |
| APOBEC3G | 22 | rs1964240 | 7 | 28812369 | -0.151132654 | 6.96E-05 | 0.924166007 |
| APOBEC3G | 22 | rs34563639 | 2 | 238891107 | 0.13507495 | 7.08E-05 | 0.924166007 |
| APOBEC3G | 22 | rs11018183 | 10 | 127540509 | -0.316177988 | 7.28E-05 | 0.924166007 |
| APOBEC3G | 22 | rs7671093 | 4 | 100594296 | 0.13064022 | 7.31E-05 | 0.924166007 |
| APOBEC3G | 22 | rs7092714 | 10 | 5359082 | -0.177470444 | 7.31E-05 | 0.924166007 |
| APOBEC3G | 22 | rs4440298 | 4 | 117300937 | 0.382814113 | 7.33E-05 | 0.924166007 |
| APOBEC3G | 22 | rs9957682 | 18 | 74431368 | 0.367953534 | 7.75E-05 | 0.924166007 |
| APOBEC3G | 22 | rs6943781 | 7 | 41832013 | -0.130953213 | 7.89E-05 | 0.924166007 |
| APOBEC3G | 22 | rs10517026 | 4 | 23955631 | -0.16750575 | 7.94E-05 | 0.924166007 |
| APOBEC3G | 22 | rs763821 | 1 | 15341780 | 0.132391287 | 7.95E-05 | 0.924166007 |
| APOBEC3G | 22 | rs884779,rs884779 | 18 | 80181018 | 0.169649345 | 8.07E-05 | 0.924166007 |
| APOBEC3G | 22 | rs35667410,rs35667410 | 18 | 80194850 | 0.169649345 | 8.07E-05 | 0.924166007 |
| APOBEC3G | 22 | rs59049410 | 1 | 13521537 | 0.360863486 | 8.57E-05 | 0.924166007 |
| APOBEC3G | 22 | rs11669354 | 19 | 53632280 | -0.234492762 | 8.62E-05 | 0.924166007 |
| APOBEC3G | 22 | rs62101568 | 18 | 80176788 | 0.168194484 | 9.11E-05 | 0.924166007 |
| APOBEC3G | 22 | rs2230721 | 4 | 6302117 | -0.233612554 | 9.25E-05 | 0.924166007 |
| APOBEC3G | 22 | rs2282912 | 7 | 28805450 | -0.151149481 | 9.57E-05 | 0.924166007 |
| APOBEC3G | 22 | rs11758028 | 6 | 66920680 | -0.140229184 | 9.59E-05 | 0.924166007 |
| APOBEC3G | 22 | rs7843587 | 8 | 134965190 | -0.129499525 | 9.87E-05 | 0.924166007 |

**Supplementary Table 2i.** *Trans*-eQTL identified for the *APOBEC3H* in non-involved lung tissue of 408 Italian lung adenocarcinoma patients ($P < 1.0 \times 10^{-4}$), listed in order of *P*-value.

| gene | gene chr | SNP | SNP chr | SNP position | beta | p-value | FDR |
|---|---|---|---|---|---|---|---|
| APOBEC3H | 22 | rs56363627 | 10 | 11019327 | -0.152167269 | 9.54E-07 | 0.65901749 |
| APOBEC3H | 22 | rs1687898 | 3 | 4371747 | 0.331394184 | 1.21E-06 | 0.667197673 |
| APOBEC3H | 22 | rs75794433 | 2 | 53462777 | 0.201629684 | 1.90E-06 | 0.805920633 |
| APOBEC3H | 22 | rs12035162 | 1 | 156693333 | 0.280345446 | 2.16E-06 | 0.80896024 |
| APOBEC3H | 22 | rs7996321 | 13 | 105346891 | 0.292825595 | 4.95E-06 | 0.84453778 |
| APOBEC3H | 22 | rs12819125 | 12 | 91992272 | 0.165405483 | 5.16E-06 | 0.84453778 |
| APOBEC3H | 22 | rs7897986 | 10 | 68453411 | -0.203000312 | 5.85E-06 | 0.84453778 |
| APOBEC3H | 22 | rs2127247 | 15 | 94712673 | 0.167816985 | 6.38E-06 | 0.84453778 |
| APOBEC3H | 22 | rs10905883 | 10 | 11025109 | -0.123920002 | 6.49E-06 | 0.84453778 |
| APOBEC3H | 22 | rs72631374 | 17 | 74938730 | 0.205121303 | 7.93E-06 | 0.84453778 |
| APOBEC3H | 22 | rs78367470 | 12 | 119020882 | 0.222672695 | 8.46E-06 | 0.84453778 |
| APOBEC3H | 22 | rs4862131 | 4 | 178769724 | 0.101347875 | 1.03E-05 | 0.84453778 |
| APOBEC3H | 22 | rs1469746 | 18 | 40488775 | 0.218694965 | 1.10E-05 | 0.84453778 |
| APOBEC3H | 22 | rs7108831 | 11 | 76193339 | 0.12088188 | 1.10E-05 | 0.84453778 |
| APOBEC3H | 22 | rs10455823 | 6 | 158508591 | 0.252211499 | 1.13E-05 | 0.84453778 |
| APOBEC3H | 22 | rs73875443 | 3 | 161016846 | 0.295506226 | 1.14E-05 | 0.84453778 |
| APOBEC3H | 22 | rs77028530 | 3 | 184631800 | 0.229478211 | 1.17E-05 | 0.84453778 |
| APOBEC3H | 22 | rs11834727 | 12 | 91992437 | 0.159642085 | 1.20E-05 | 0.84453778 |
| APOBEC3H | 22 | rs201078 | 10 | 10960789 | -0.118883005 | 1.22E-05 | 0.84453778 |
| APOBEC3H | 22 | rs12612847 | 2 | 84122891 | 0.102021223 | 1.28E-05 | 0.848481351 |
| APOBEC3H | 22 | rs12826769 | 12 | 92002100 | 0.155615834 | 1.32E-05 | 0.848481351 |
| APOBEC3H | 22 | rs56703600 | 1 | 69697871 | 0.1535727 | 1.40E-05 | 0.848481351 |
| APOBEC3H | 22 | rs17693059 | 2 | 173342529 | 0.145748071 | 1.59E-05 | 0.853417593 |
| APOBEC3H | 22 | rs1500305 | 9 | 8279817 | 0.132307649 | 1.60E-05 | 0.853417593 |
| APOBEC3H | 22 | rs2565761 | 2 | 124385524 | -0.108114827 | 1.61E-05 | 0.853417593 |
| APOBEC3H | 22 | rs7872167 | 9 | 8284955 | 0.133590031 | 1.70E-05 | 0.863488543 |
| APOBEC3H | 22 | rs34838816 | 2 | 34395066 | 0.370454222 | 1.99E-05 | 0.863488543 |
| APOBEC3H | 22 | rs73203033 | 7 | 111542695 | 0.313098581 | 2.00E-05 | 0.863488543 |
| APOBEC3H | 22 | rs11674417 | 2 | 84121504 | -0.10038238 | 2.05E-05 | 0.863488543 |
| APOBEC3H | 22 | rs11674417 | 2 | 84121504 | -0.10038238 | 2.05E-05 | 0.863488543 |
| APOBEC3H | 22 | rs11674417 | 2 | 84121504 | -0.10038238 | 2.05E-05 | 0.863488543 |
| APOBEC3H | 22 | rs11674417 | 2 | 84121504 | -0.10038238 | 2.05E-05 | 0.863488543 |
| APOBEC3H | 22 | rs72631372 | 17 | 74938619 | 0.193603446 | 2.20E-05 | 0.863488543 |
| APOBEC3H | 22 | rs117620699 | 9 | 94556291 | -0.270514971 | 2.38E-05 | 0.863488543 |
| APOBEC3H | 22 | rs72772071 | 10 | 10969858 | -0.115600378 | 2.54E-05 | 0.863488543 |
| APOBEC3H | 22 | rs74849557 | 9 | 5927828 | 0.330572291 | 2.79E-05 | 0.875040765 |
| APOBEC3H | 22 | rs814526 | 19 | 40502142 | 0.097824547 | 2.80E-05 | 0.875040765 |
| APOBEC3H | 22 | rs11879852 | 19 | 31312276 | -0.129233364 | 2.87E-05 | 0.875040765 |
| APOBEC3H | 22 | rs12688189 | X | 103734516 | 0.180516899 | 2.96E-05 | 0.875040765 |
| APOBEC3H | 22 | rs11120116 | 1 | 213380922 | -0.10196488 | 3.05E-05 | 0.875040765 |
| APOBEC3H | 22 | rs7862136 | 9 | 8285325 | 0.128639418 | 3.26E-05 | 0.875040765 |
| APOBEC3H | 22 | rs1843120 | 5 | 162541036 | 0.290202228 | 3.37E-05 | 0.875040765 |
| APOBEC3H | 22 | rs74332186 | 5 | 162584433 | 0.290202228 | 3.37E-05 | 0.875040765 |
| APOBEC3H | 22 | rs7306999 | 12 | 69131766 | 0.101101684 | 3.38E-05 | 0.875040765 |
| APOBEC3H | 22 | rs2571147 | 19 | 44401885 | -0.114666383 | 3.56E-05 | 0.875040765 |
| APOBEC3H | 22 | rs201489090 | 5 | 116580638 | 0.110204635 | 4.17E-05 | 0.896458005 |
| APOBEC3H | 22 | rs63750072 | 17 | 45983493 | 0.179704719 | 4.20E-05 | 0.896458005 |
| APOBEC3H | 22 | rs117844251 | 13 | 38936446 | 0.205845963 | 4.26E-05 | 0.896890823 |

**Supplementary Table 2i.** *Trans*-eQTL identified for the *APOBEC3H* in non-involved lung tissue of 408 Italian lung adenocarcinoma patients ($P < 1.0 \times 10^{-4}$), listed in order of *P*-value.

| gene | gene chr | SNP | SNP chr | SNP position | beta | p-value | FDR |
|---|---|---|---|---|---|---|---|
| APOBEC3H | 22 | rs12899535 | 15 | 66727015 | 0.191480318 | 4.32E-05 | 0.901579715 |
| APOBEC3H | 22 | rs375807 | 2 | 203825857 | 0.136680964 | 4.34E-05 | 0.903928114 |
| APOBEC3H | 22 | rs77417763 | 12 | 118970443 | 0.21534529 | 4.37E-05 | 0.906462302 |
| APOBEC3H | 22 | rs4862147 | 4 | 178777205 | 0.093750391 | 4.55E-05 | 0.909284759 |
| APOBEC3H | 22 | rs2215053 | 17 | 12810099 | 0.202667623 | 4.75E-05 | 0.912369094 |
| APOBEC3H | 22 | rs7952005 | 11 | 76160246 | 0.133178834 | 4.90E-05 | 0.914201928 |
| APOBEC3H | 22 | rs7499860 | 16 | 78946504 | 0.138789768 | 4.97E-05 | 0.914201928 |
| APOBEC3H | 22 | rs67051757 | 16 | 78946521 | 0.138789768 | 4.97E-05 | 0.914201928 |
| APOBEC3H | 22 | rs3939465 | 13 | 89784191 | -0.088336259 | 5.02E-05 | 0.914201928 |
| APOBEC3H | 22 | rs11563848 | 7 | 90519058 | -0.11809843 | 5.11E-05 | 0.914201928 |
| APOBEC3H | 22 | rs7146456 | 14 | 68572264 | -0.090617494 | 5.11E-05 | 0.914201928 |
| APOBEC3H | 22 | rs2830196 | 21 | 26398526 | 0.22641799 | 5.12E-05 | 0.914201928 |
| APOBEC3H | 22 | rs10851813 | 15 | 70139853 | 0.110262302 | 5.18E-05 | 0.914201928 |
| APOBEC3H | 22 | rs12170237 | 22 | 36963448 | 0.225464062 | 5.20E-05 | 0.914201928 |
| APOBEC3H | 22 | rs62253780 | 3 | 63727863 | -0.117679818 | 5.31E-05 | 0.91575718 |
| APOBEC3H | 22 | rs2158061 | 7 | 24795863 | 0.303821235 | 5.41E-05 | 0.91575718 |
| APOBEC3H | 22 | rs58182085 | 18 | 23057516 | -0.122971675 | 5.42E-05 | 0.91575718 |
| APOBEC3H | 22 | rs6845787 | 4 | 150014701 | 0.121268565 | 5.43E-05 | 0.91575718 |
| APOBEC3H | 22 | rs28460874 | 16 | 56592128 | 0.116534129 | 5.49E-05 | 0.91575718 |
| APOBEC3H | 22 | rs80024341 | 16 | 89178366 | -0.154429903 | 5.92E-05 | 0.918110223 |
| APOBEC3H | 22 | rs62171016 | 2 | 124352699 | -0.09495219 | 6.07E-05 | 0.918110223 |
| APOBEC3H | 22 | rs9327012 | 5 | 116586973 | 0.107661433 | 6.29E-05 | 0.923619639 |
| APOBEC3H | 22 | rs17664293 | 15 | 94719978 | 0.13534457 | 6.34E-05 | 0.924166007 |
| APOBEC3H | 22 | rs298991 | 4 | 118724845 | -0.09713263 | 6.56E-05 | 0.924166007 |
| APOBEC3H | 22 | rs17528236 | 13 | 39142947 | 0.225543535 | 6.67E-05 | 0.924166007 |
| APOBEC3H | 22 | rs9560441 | 13 | 89789504 | -0.100891952 | 6.71E-05 | 0.924166007 |
| APOBEC3H | 22 | rs1932310 | 13 | 89791354 | -0.100891952 | 6.71E-05 | 0.924166007 |
| APOBEC3H | 22 | rs780010 | 2 | 124309174 | -0.101395504 | 6.98E-05 | 0.924166007 |
| APOBEC3H | 22 | rs80079773 | 14 | 78576028 | 0.217505181 | 6.99E-05 | 0.924166007 |
| APOBEC3H | 22 | rs537001 | 11 | 64019639 | 0.090753972 | 7.08E-05 | 0.924166007 |
| APOBEC3H | 22 | rs9522714 | 13 | 89791476 | -0.086566982 | 7.13E-05 | 0.924166007 |
| APOBEC3H | 22 | rs729990 | 1 | 32977665 | 0.183538074 | 7.18E-05 | 0.924166007 |
| APOBEC3H | 22 | rs56140076 | 20 | 58393505 | 0.200056665 | 7.18E-05 | 0.924166007 |
| APOBEC3H | 22 | rs17544336 | 20 | 58432538 | 0.200056665 | 7.18E-05 | 0.924166007 |
| APOBEC3H | 22 | rs199923819 | 20 | 58444435 | 0.200056665 | 7.18E-05 | 0.924166007 |
| APOBEC3H | 22 | rs60614524 | 10 | 61440747 | 0.298356102 | 7.24E-05 | 0.924166007 |
| APOBEC3H | 22 | rs34684289 | 11 | 25370585 | 0.142099639 | 7.70E-05 | 0.924166007 |
| APOBEC3H | 22 | rs7687229 | 4 | 157140712 | 0.276594511 | 7.71E-05 | 0.924166007 |
| APOBEC3H | 22 | rs77036899 | 6 | 52224141 | 0.195005107 | 7.80E-05 | 0.924166007 |
| APOBEC3H | 22 | rs59973928 | 4 | 100604881 | 0.272027223 | 7.89E-05 | 0.924166007 |
| APOBEC3H | 22 | rs7143214 | 14 | 83627876 | -0.120280498 | 8.29E-05 | 0.924166007 |
| APOBEC3H | 22 | rs2722669 | 19 | 44409065 | -0.125006625 | 8.46E-05 | 0.924166007 |
| APOBEC3H | 22 | rs682769 | 8 | 101615727 | 0.099383366 | 8.61E-05 | 0.924166007 |
| APOBEC3H | 22 | rs7150510 | 14 | 83629522 | -0.119673586 | 8.73E-05 | 0.924166007 |
| APOBEC3H | 22 | rs12456875 | 18 | 22830208 | -0.12539394 | 8.81E-05 | 0.924166007 |
| APOBEC3H | 22 | rs11082173 | 18 | 22858048 | -0.12539394 | 8.81E-05 | 0.924166007 |
| APOBEC3H | 22 | rs4243474 | 6 | 144775204 | 0.099729606 | 8.83E-05 | 0.924166007 |
| APOBEC3H | 22 | rs4577027 | 15 | 94711948 | 0.131133703 | 9.01E-05 | 0.924166007 |

**Supplementary Table 2i.** *Trans*-eQTL identified for the *APOBEC3H* in non-involved lung tissue of 408 Italian lung adenocarcinoma patients ($P < 1.0 \times 10^{-4}$), listed in order of *P*-value.

| gene | gene chr | SNP | SNP chr | SNP position | beta | p-value | FDR |
|---|---|---|---|---|---|---|---|
| APOBEC3H | 22 | rs1308154 | 1 | 151716352 | 0.350179984 | 9.03E-05 | 0.924166007 |
| APOBEC3H | 22 | rs11974998 | 7 | 9654040 | 0.176723571 | 9.20E-05 | 0.924166007 |
| APOBEC3H | 22 | rs73440428 | 12 | 129826180 | 0.166507679 | 9.20E-05 | 0.924166007 |
| APOBEC3H | 22 | rs12649030 | 4 | 166118943 | 0.095609952 | 9.22E-05 | 0.924166007 |
| APOBEC3H | 22 | rs79800383 | 3 | 59415067 | 0.121321263 | 9.44E-05 | 0.924166007 |
| APOBEC3H | 22 | rs62443340 | 7 | 38221616 | 0.186630546 | 9.47E-05 | 0.924166007 |
| APOBEC3H | 22 | rs4517886 | 18 | 22982922 | -0.124340405 | 9.48E-05 | 0.924166007 |
| APOBEC3H | 22 | rs1027582 | 9 | 8267854 | 0.102924224 | 9.63E-05 | 0.924166007 |
| APOBEC3H | 22 | rs73368368 | 10 | 122024489 | 0.2337274 | 9.82E-05 | 0.924166007 |
| APOBEC3H | 22 | rs77188729 | 2 | 128516313 | 0.334838285 | 9.86E-05 | 0.924166007 |
| APOBEC3H | 22 | rs4550540 | 18 | 25311667 | -0.093879924 | 9.88E-05 | 0.924166007 |
| APOBEC3H | 22 | rs12650979 | 4 | 166114449 | 0.089995843 | 9.96E-05 | 0.924166007 |

**Supplementary Table 2j.** *Trans*-eQTL identified for the *IFITM2* in non-involved lung tissue of 408 Italian lung adenocarcinoma patients ($P < 1.0 \times 10^{-4}$), listed in order of *P*-value.

| gene | gene chr | SNP | SNP chr | SNP position | beta | p-value | FDR |
|---|---|---|---|---|---|---|---|
| IFITM2 | 11 | rs62552897 | 9 | 74448907 | -0.144462897 | 1.77E-07 | 0.493509947 |
| IFITM2 | 11 | rs17612218 | 9 | 74573969 | -0.18852787 | 1.79E-07 | 0.493509947 |
| IFITM2 | 11 | rs11792875 | 9 | 74389326 | -0.138082086 | 4.81E-07 | 0.65901749 |
| IFITM2 | 11 | rs1872826 | 15 | 79615435 | 0.1197724 | 3.80E-06 | 0.84453778 |
| IFITM2 | 11 | rs8055622 | 16 | 26754164 | -0.374261254 | 4.26E-06 | 0.84453778 |
| IFITM2 | 11 | rs116722052 | 4 | 35371319 | 0.258359698 | 6.44E-06 | 0.84453778 |
| IFITM2 | 11 | rs7163832 | 15 | 79614002 | 0.116973744 | 6.60E-06 | 0.84453778 |
| IFITM2 | 11 | rs3769567 | 2 | 32808990 | -0.113544826 | 7.21E-06 | 0.84453778 |
| IFITM2 | 11 | rs10173352 | 2 | 32832322 | -0.115588209 | 8.74E-06 | 0.84453778 |
| IFITM2 | 11 | rs2829425 | 21 | 24828862 | -0.13310997 | 9.77E-06 | 0.84453778 |
| IFITM2 | 11 | rs200811796 | 21 | 24831419 | -0.13310997 | 9.77E-06 | 0.84453778 |
| IFITM2 | 11 | rs6495429 | 15 | 79607526 | 0.114892586 | 9.84E-06 | 0.84453778 |
| IFITM2 | 11 | rs704341 | 3 | 61963062 | 0.170460342 | 1.04E-05 | 0.84453778 |
| IFITM2 | 11 | rs916288 | 3 | 50475726 | -0.116543536 | 1.16E-05 | 0.84453778 |
| IFITM2 | 11 | rs2829429 | 21 | 24832383 | -0.126827208 | 1.24E-05 | 0.848481351 |
| IFITM2 | 11 | rs13144677 | 4 | 5839922 | 0.120242004 | 1.31E-05 | 0.848481351 |
| IFITM2 | 11 | rs3816385 | 15 | 64126456 | 0.126485819 | 1.34E-05 | 0.848481351 |
| IFITM2 | 11 | rs3848182 | 15 | 79626369 | 0.111728493 | 1.44E-05 | 0.848481351 |
| IFITM2 | 11 | rs2865194 | 15 | 79616482 | 0.110342269 | 1.48E-05 | 0.848481351 |
| IFITM2 | 11 | rs3769556 | 2 | 32820507 | -0.111186236 | 1.61E-05 | 0.853417593 |
| IFITM2 | 11 | rs7952778 | 12 | 24055375 | 0.111786247 | 1.70E-05 | 0.863488543 |
| IFITM2 | 11 | rs28366555 | 15 | 79594813 | 0.226728199 | 2.07E-05 | 0.863488543 |
| IFITM2 | 11 | rs11853292 | 15 | 64085609 | 0.123529916 | 2.14E-05 | 0.863488543 |
| IFITM2 | 11 | rs73489804 | 18 | 78045734 | -0.159980734 | 2.26E-05 | 0.863488543 |
| IFITM2 | 11 | rs7418357 | 1 | 5411457 | -0.137860294 | 2.35E-05 | 0.863488543 |
| IFITM2 | 11 | rs13211009 | 6 | 12515735 | -0.119619396 | 2.39E-05 | 0.863488543 |
| IFITM2 | 11 | rs7027790 | 9 | 20319777 | -0.108531794 | 2.45E-05 | 0.863488543 |
| IFITM2 | 11 | rs9305744 | 21 | 41471061 | -0.136065904 | 2.47E-05 | 0.863488543 |
| IFITM2 | 11 | rs13048709 | 21 | 42605776 | -0.201486892 | 2.62E-05 | 0.863488543 |
| IFITM2 | 11 | rs10236407 | 7 | 8996185 | -0.2151301 | 2.79E-05 | 0.875040765 |
| IFITM2 | 11 | rs176400 | 2 | 32408501 | -0.113034207 | 3.00E-05 | 0.875040765 |
| IFITM2 | 11 | rs8097909 | 18 | 78028508 | -0.147608725 | 3.27E-05 | 0.875040765 |
| IFITM2 | 11 | rs4952302 | 2 | 32826462 | -0.107813512 | 3.31E-05 | 0.875040765 |
| IFITM2 | 11 | rs80067748 | 3 | 149142962 | 0.299731701 | 3.36E-05 | 0.875040765 |
| IFITM2 | 11 | rs9920588 | 15 | 79600050 | 0.194834852 | 3.46E-05 | 0.875040765 |
| IFITM2 | 11 | rs1317866 | 15 | 79604698 | 0.194834852 | 3.46E-05 | 0.875040765 |
| IFITM2 | 11 | rs2109001 | 16 | 5661290 | 0.120053692 | 3.63E-05 | 0.875040765 |
| IFITM2 | 11 | rs4811694 | 20 | 56454332 | 0.101149442 | 4.14E-05 | 0.896458005 |
| IFITM2 | 11 | rs10088747 | 8 | 30009325 | 0.163566857 | 4.21E-05 | 0.896458005 |
| IFITM2 | 11 | rs9886212 | 7 | 9006246 | -0.242867215 | 4.23E-05 | 0.896458005 |
| IFITM2 | 11 | rs2829408 | 21 | 24817021 | -0.113761835 | 4.31E-05 | 0.901453344 |
| IFITM2 | 11 | rs2418156 | 12 | 24035664 | 0.10789654 | 4.51E-05 | 0.909284759 |
| IFITM2 | 11 | rs9308929 | 2 | 32806291 | 0.101379753 | 4.77E-05 | 0.912369094 |
| IFITM2 | 11 | rs13046013 | 21 | 42605757 | -0.195767979 | 4.99E-05 | 0.914201928 |
| IFITM2 | 11 | rs8109957 | 19 | 7192582 | -0.171938274 | 5.10E-05 | 0.914201928 |
| IFITM2 | 11 | rs10809863 | 9 | 12956313 | -0.105895232 | 5.18E-05 | 0.914201928 |
| IFITM2 | 11 | rs8052167 | 16 | 83681944 | -0.102712536 | 5.38E-05 | 0.91575718 |
| IFITM2 | 11 | rs16875859 | 5 | 78746392 | 0.180549964 | 5.73E-05 | 0.918110223 |

**Supplementary Table 2j.** *Trans*-eQTL identified for the *IFITM2* in non-involved lung tissue of 408 Italian lung adenocarcinoma patients ($P < 1.0 \times 10^{-4}$), listed in order of *P*-value.

| gene | gene chr | SNP | SNP chr | SNP position | beta | p-value | FDR |
|---|---|---|---|---|---|---|---|
| IFITM2 | 11 | rs4259413 | 8 | 83677570 | 0.100449954 | 5.83E-05 | 0.918110223 |
| IFITM2 | 11 | rs6487369 | 12 | 24017085 | 0.10269733 | 5.90E-05 | 0.918110223 |
| IFITM2 | 11 | rs9904568 | 17 | 73157532 | 0.127912408 | 5.99E-05 | 0.918110223 |
| IFITM2 | 11 | rs2576267 | 1 | 216886589 | -0.1338449 | 6.00E-05 | 0.918110223 |
| IFITM2 | 11 | rs76377903 | 16 | 57664510 | -0.28986676 | 6.08E-05 | 0.918110223 |
| IFITM2 | 11 | rs2710628 | 2 | 32533286 | -0.099555151 | 6.11E-05 | 0.918110223 |
| IFITM2 | 11 | rs4928190 | 3 | 99102894 | -0.225647819 | 6.15E-05 | 0.919202897 |
| IFITM2 | 11 | rs75079377 | 18 | 78028154 | -0.148370608 | 6.17E-05 | 0.919202897 |
| IFITM2 | 11 | rs11588081 | 1 | 48532770 | -0.111556992 | 6.24E-05 | 0.920563722 |
| IFITM2 | 11 | rs1536788 | 9 | 12963198 | -0.104918134 | 6.39E-05 | 0.924166007 |
| IFITM2 | 11 | rs11143958 | 9 | 74395765 | -0.103613968 | 6.66E-05 | 0.924166007 |
| IFITM2 | 11 | rs12437511 | 15 | 79613586 | 0.181272685 | 6.66E-05 | 0.924166007 |
| IFITM2 | 11 | rs61737062 | 21 | 43749042 | 0.334093306 | 6.70E-05 | 0.924166007 |
| IFITM2 | 11 | rs4696652 | 4 | 151069620 | -0.169713431 | 6.82E-05 | 0.924166007 |
| IFITM2 | 11 | rs155394 | 3 | 1381950 | -0.162075372 | 7.28E-05 | 0.924166007 |
| IFITM2 | 11 | rs7692068 | 4 | 42108378 | -0.279421273 | 7.31E-05 | 0.924166007 |
| IFITM2 | 11 | rs13145829 | 4 | 151093501 | -0.166901753 | 7.73E-05 | 0.924166007 |
| IFITM2 | 11 | rs62016096 | 15 | 94327332 | 0.189577474 | 7.93E-05 | 0.924166007 |
| IFITM2 | 11 | rs55776030 | 17 | 8692127 | -0.10844712 | 8.09E-05 | 0.924166007 |
| IFITM2 | 11 | rs1179726 | 2 | 216866585 | 0.098717854 | 8.26E-05 | 0.924166007 |
| IFITM2 | 11 | rs342231 | 7 | 106692000 | -0.237900214 | 8.34E-05 | 0.924166007 |
| IFITM2 | 11 | rs28703682 | 3 | 53453024 | -0.101483204 | 8.39E-05 | 0.924166007 |
| IFITM2 | 11 | rs12779337 | 10 | 53492627 | 0.240628254 | 8.42E-05 | 0.924166007 |
| IFITM2 | 11 | rs41280357 | 4 | 52605030 | 0.322736659 | 8.49E-05 | 0.924166007 |
| IFITM2 | 11 | rs117446301 | 10 | 649961 | -0.305180569 | 8.57E-05 | 0.924166007 |
| IFITM2 | 11 | rs342238 | 7 | 106695815 | -0.237269176 | 8.73E-05 | 0.924166007 |
| IFITM2 | 11 | rs4952312 | 2 | 32876841 | 0.164357956 | 9.22E-05 | 0.924166007 |
| IFITM2 | 11 | rs1202222 | 7 | 3616433 | -0.099706321 | 9.25E-05 | 0.924166007 |
| IFITM2 | 11 | rs6968401 | 7 | 44330732 | 0.098140155 | 9.35E-05 | 0.924166007 |
| IFITM2 | 11 | rs73602872 | 8 | 53281710 | -0.211527691 | 9.36E-05 | 0.924166007 |
| IFITM2 | 11 | rs4315885 | 5 | 69028656 | -0.098637042 | 9.38E-05 | 0.924166007 |
| IFITM2 | 11 | rs11761910 | 7 | 32094526 | 0.1102379 | 9.56E-05 | 0.924166007 |
| IFITM2 | 11 | rs10216516 | 8 | 1973423 | 0.112434082 | 9.61E-05 | 0.924166007 |

**Supplementary Table 2k.** *Trans*-eQTL identified for the *IFITM3* in non-involved lung tissue of 408 Italian lung adenocarcinoma patients ($P < 1.0 \times 10^{-4}$), listed in order of *P*-value.

| gene | gene chr | SNP | SNP chr | SNP position | beta | p-value | FDR |
|---|---|---|---|---|---|---|---|
| IFITM3 | 11 | rs62552897 | 9 | 74448907 | -0.140092057 | 1.47E-07 | 0.493509947 |
| IFITM3 | 11 | rs11792875 | 9 | 74389326 | -0.130215938 | 8.54E-07 | 0.65901749 |
| IFITM3 | 11 | rs17612218 | 9 | 74573969 | -0.173922598 | 6.03E-07 | 0.65901749 |
| IFITM3 | 11 | rs6487369 | 12 | 24017085 | 0.115459352 | 2.56E-06 | 0.812807829 |
| IFITM3 | 11 | rs3732341 | 2 | 240716127 | 0.10779723 | 1.09E-05 | 0.84453778 |
| IFITM3 | 11 | rs292182 | 5 | 36954710 | 0.105156005 | 1.20E-05 | 0.84453778 |
| IFITM3 | 11 | rs4259413 | 8 | 83677570 | 0.110444579 | 4.21E-06 | 0.84453778 |
| IFITM3 | 11 | rs4379441 | 8 | 83704465 | 0.108744906 | 6.01E-06 | 0.84453778 |
| IFITM3 | 11 | rs11143958 | 9 | 74395765 | -0.109394221 | 1.20E-05 | 0.84453778 |
| IFITM3 | 11 | rs1021671 | 12 | 24023878 | -0.109207421 | 1.11E-05 | 0.84453778 |
| IFITM3 | 11 | rs34252042 | 13 | 42884542 | 0.106607796 | 1.09E-05 | 0.84453778 |
| IFITM3 | 11 | rs13170811 | 5 | 36795628 | 0.104014985 | 1.45E-05 | 0.848481351 |
| IFITM3 | 11 | rs7846815 | 9 | 74432686 | -0.10849838 | 1.49E-05 | 0.848481351 |
| IFITM3 | 11 | rs9315970 | 13 | 42890159 | -0.108359073 | 1.29E-05 | 0.848481351 |
| IFITM3 | 11 | rs201290424 | 13 | 94936686 | -0.108401412 | 1.50E-05 | 0.848481351 |
| IFITM3 | 11 | rs8055622 | 16 | 26754164 | -0.34218642 | 1.32E-05 | 0.848481351 |
| IFITM3 | 11 | rs10173297 | 2 | 3666539 | 0.152340552 | 2.65E-05 | 0.863488543 |
| IFITM3 | 11 | rs4555902 | 6 | 66924950 | -0.102805941 | 2.22E-05 | 0.863488543 |
| IFITM3 | 11 | rs72686244 | 8 | 83544178 | 0.103937791 | 2.64E-05 | 0.863488543 |
| IFITM3 | 11 | rs7845183 | 8 | 83669463 | 0.097960825 | 2.30E-05 | 0.863488543 |
| IFITM3 | 11 | rs7952778 | 12 | 24055375 | 0.1058574 | 2.39E-05 | 0.863488543 |
| IFITM3 | 11 | rs7983005 | 13 | 42888893 | -0.103196505 | 2.38E-05 | 0.863488543 |
| IFITM3 | 11 | rs13046013 | 21 | 42605757 | -0.195542463 | 2.58E-05 | 0.863488543 |
| IFITM3 | 11 | rs13048709 | 21 | 42605776 | -0.196267334 | 2.13E-05 | 0.863488543 |
| IFITM3 | 11 | rs2065708 | 1 | 45512013 | 0.125177427 | 3.16E-05 | 0.875040765 |
| IFITM3 | 11 | rs12757540 | 1 | 205918261 | -0.144314652 | 3.00E-05 | 0.875040765 |
| IFITM3 | 11 | rs116722052 | 4 | 35371319 | 0.231838875 | 2.74E-05 | 0.875040765 |
| IFITM3 | 11 | rs200606028 | 7 | 151899983 | -0.103218044 | 3.58E-05 | 0.875040765 |
| IFITM3 | 11 | rs1586563 | 8 | 83624772 | 0.100269281 | 3.07E-05 | 0.875040765 |
| IFITM3 | 11 | rs2418156 | 12 | 24035664 | 0.106010345 | 3.16E-05 | 0.875040765 |
| IFITM3 | 11 | rs9676648 | 19 | 46235736 | -0.216827527 | 3.45E-05 | 0.875040765 |
| IFITM3 | 11 | rs1797590 | 4 | 74462824 | 0.101505897 | 3.96E-05 | 0.894226918 |
| IFITM3 | 11 | rs4481415 | 6 | 66937020 | -0.108990491 | 4.24E-05 | 0.896458005 |
| IFITM3 | 11 | rs4484508 | 6 | 66937497 | -0.108937256 | 4.25E-05 | 0.896458005 |
| IFITM3 | 11 | rs17560636 | 10 | 52043995 | 0.107957764 | 4.05E-05 | 0.896458005 |
| IFITM3 | 11 | rs202012889 | 10 | 52048802 | 0.108319923 | 4.21E-05 | 0.896458005 |
| IFITM3 | 11 | rs202206233 | 1 | 205918320 | -0.141238018 | 4.29E-05 | 0.901453344 |
| IFITM3 | 11 | rs73219736 | 8 | 14395555 | -0.14375506 | 4.55E-05 | 0.909284759 |
| IFITM3 | 11 | rs11211133 | 1 | 45525407 | 0.12264303 | 4.63E-05 | 0.912369094 |
| IFITM3 | 11 | rs117764140 | 6 | 150530977 | 0.269489955 | 4.76E-05 | 0.912369094 |
| IFITM3 | 11 | rs12461351 | 19 | 46193684 | -0.181957598 | 4.66E-05 | 0.912369094 |
| IFITM3 | 11 | rs117446301 | 10 | 649961 | -0.304063348 | 4.83E-05 | 0.914201928 |
| IFITM3 | 11 | rs11633509 | 15 | 93153235 | -0.105360897 | 5.49E-05 | 0.91575718 |
| IFITM3 | 11 | rs724749 | 4 | 124804199 | -0.096761462 | 5.97E-05 | 0.918110223 |
| IFITM3 | 11 | rs2112298 | 5 | 72469712 | -0.120990935 | 5.62E-05 | 0.918110223 |
| IFITM3 | 11 | rs2761025 | 9 | 113397580 | 0.220378839 | 5.79E-05 | 0.918110223 |
| IFITM3 | 11 | rs1044856 | 13 | 42888286 | 0.098226339 | 5.84E-05 | 0.918110223 |
| IFITM3 | 11 | rs34138846 | 19 | 46139343 | -0.172472877 | 6.01E-05 | 0.918110223 |

**Supplementary Table 2k.** *Trans*-eQTL identified for the *IFITM3* in non-involved lung tissue of 408 Italian lung adenocarcinoma patients ($P < 1.0 \times 10^{-4}$), listed in order of *P*-value.

| gene | gene chr | SNP | SNP chr | SNP position | beta | p-value | FDR |
|---|---|---|---|---|---|---|---|
| IFITM3 | 11 | rs7990517 | 13 | 94537841 | 0.23394787 | 6.17E-05 | 0.919202897 |
| IFITM3 | 11 | rs9590032 | 13 | 94553370 | 0.23394787 | 6.17E-05 | 0.919202897 |
| IFITM3 | 11 | rs11184332 | 1 | 104872034 | -0.320766229 | 6.97E-05 | 0.924166007 |
| IFITM3 | 11 | rs6794523 | 3 | 3066007 | 0.097025157 | 8.95E-05 | 0.924166007 |
| IFITM3 | 11 | rs17182624 | 4 | 74406532 | 0.097487575 | 8.62E-05 | 0.924166007 |
| IFITM3 | 11 | rs9308191 | 4 | 145665062 | -0.09611998 | 9.91E-05 | 0.924166007 |
| IFITM3 | 11 | rs13109257 | 4 | 145681821 | -0.096204787 | 8.22E-05 | 0.924166007 |
| IFITM3 | 11 | rs6860727 | 5 | 37179239 | 0.094975089 | 8.58E-05 | 0.924166007 |
| IFITM3 | 11 | rs1890311 | 6 | 76124025 | -0.134428944 | 9.68E-05 | 0.924166007 |
| IFITM3 | 11 | rs6968401 | 7 | 44330732 | 0.094966554 | 8.73E-05 | 0.924166007 |
| IFITM3 | 11 | rs6987971 | 8 | 141432438 | -0.099692421 | 7.89E-05 | 0.924166007 |
| IFITM3 | 11 | rs10809863 | 9 | 12956313 | -0.099599252 | 7.87E-05 | 0.924166007 |
| IFITM3 | 11 | rs1536788 | 9 | 12963198 | -0.0992138 | 8.79E-05 | 0.924166007 |
| IFITM3 | 11 | rs7023422 | 9 | 20318799 | -0.099637854 | 8.57E-05 | 0.924166007 |
| IFITM3 | 11 | rs75988114 | 9 | 113607851 | 0.165668522 | 8.10E-05 | 0.924166007 |
| IFITM3 | 11 | rs716781 | 10 | 89650064 | -0.113764123 | 9.87E-05 | 0.924166007 |
| IFITM3 | 11 | rs7076259 | 10 | 89652845 | -0.113764123 | 9.87E-05 | 0.924166007 |
| IFITM3 | 11 | rs3908479 | 10 | 89670424 | -0.115225037 | 7.94E-05 | 0.924166007 |
| IFITM3 | 11 | rs10744263 | 12 | 126480833 | -0.126109454 | 7.20E-05 | 0.924166007 |
| IFITM3 | 11 | rs2138125 | 17 | 80814689 | 0.093801513 | 9.89E-05 | 0.924166007 |
| IFITM3 | 11 | rs35968328 | 18 | 49590837 | -0.138347238 | 7.54E-05 | 0.924166007 |

**Supplementary Table 2l.** *Trans*-eQTL identified for the *PARP12* in non-involved lung tissue of 408 Italian lung adenocarcinoma patients ($P < 1.0 \times 10^{-4}$), listed in order of *P*-value.

| gene | gene chr | SNP | SNP chr | SNP position | beta | p-value | FDR |
|---|---|---|---|---|---|---|---|
| PARP12 | 7 | rs2941816 | 18 | 78748001 | 0.1231411 | 2.67E-06 | 0.819496588 |
| PARP12 | 7 | rs8012626 | 14 | 30283902 | -0.114884904 | 4.77E-06 | 0.84453778 |
| PARP12 | 7 | rs764200 | 1 | 187790468 | 0.115329051 | 1.28E-05 | 0.848481351 |
| PARP12 | 7 | rs12661851 | 6 | 12175965 | -0.115478588 | 1.44E-05 | 0.848481351 |
| PARP12 | 7 | rs829914 | 6 | 45737831 | 0.315005606 | 1.33E-05 | 0.848481351 |
| PARP12 | 7 | rs12581308 | 12 | 128532856 | -0.176608224 | 1.47E-05 | 0.848481351 |
| PARP12 | 7 | rs12321655 | 12 | 128534256 | -0.176608224 | 1.47E-05 | 0.848481351 |
| PARP12 | 7 | rs9949091 | 18 | 78747259 | 0.121364584 | 1.61E-05 | 0.853417593 |
| PARP12 | 7 | rs9582 | 19 | 17212911 | -0.105074409 | 1.59E-05 | 0.853417593 |
| PARP12 | 7 | rs10242548 | 7 | 107821704 | 0.138576815 | 1.65E-05 | 0.857061646 |
| PARP12 | 7 | rs72910161 | 2 | 21406549 | -0.197900548 | 2.61E-05 | 0.863488543 |
| PARP12 | 7 | rs10033074 | 4 | 183534161 | 0.107329831 | 1.96E-05 | 0.863488543 |
| PARP12 | 7 | rs10471588 | 5 | 63616918 | 0.120038006 | 2.08E-05 | 0.863488543 |
| PARP12 | 7 | rs200580497 | 5 | 146668191 | 0.133449452 | 1.68E-05 | 0.863488543 |
| PARP12 | 7 | rs458725 | 5 | 146673325 | 0.131464134 | 2.28E-05 | 0.863488543 |
| PARP12 | 7 | rs76329448 | 7 | 21793971 | -0.210159338 | 2.21E-05 | 0.863488543 |
| PARP12 | 7 | rs72671217 | 14 | 20247310 | -0.106096792 | 1.80E-05 | 0.863488543 |
| PARP12 | 7 | rs704391 | 3 | 64140445 | -0.102669685 | 3.54E-05 | 0.875040765 |
| PARP12 | 7 | rs11847357 | 14 | 48559103 | 0.113953206 | 3.10E-05 | 0.875040765 |
| PARP12 | 7 | rs4620786 | 12 | 94707626 | -0.109909978 | 3.72E-05 | 0.879948067 |
| PARP12 | 7 | rs67354843 | 4 | 180829124 | 0.15384017 | 3.98E-05 | 0.894226918 |
| PARP12 | 7 | rs746034 | 12 | 104598102 | -0.101463582 | 4.03E-05 | 0.896023321 |
| PARP12 | 7 | rs200799751 | 14 | 36652319 | 0.102993503 | 4.07E-05 | 0.896458005 |
| PARP12 | 7 | rs1320207 | 15 | 69954410 | -0.116047041 | 4.15E-05 | 0.896458005 |
| PARP12 | 7 | rs11124257 | 2 | 31305043 | -0.209775736 | 4.43E-05 | 0.90811064 |
| PARP12 | 7 | rs17757714 | 15 | 69965615 | -0.122129745 | 4.41E-05 | 0.90811064 |
| PARP12 | 7 | rs73165334 | 13 | 36124586 | -0.119377258 | 4.59E-05 | 0.911360524 |
| PARP12 | 7 | rs4701655 | 5 | 16237127 | -0.14670339 | 4.62E-05 | 0.912369094 |
| PARP12 | 7 | rs10195902 | 2 | 121668910 | 0.170850932 | 5.43E-05 | 0.91575718 |
| PARP12 | 7 | rs17362588 | 2 | 178856319 | -0.172391388 | 5.42E-05 | 0.91575718 |
| PARP12 | 7 | rs7379819 | 5 | 63607930 | 0.113339099 | 5.42E-05 | 0.91575718 |
| PARP12 | 7 | rs10078495 | 5 | 178455628 | -0.101159497 | 5.46E-05 | 0.91575718 |
| PARP12 | 7 | rs4784157 | 16 | 61739818 | -0.102741318 | 5.27E-05 | 0.91575718 |
| PARP12 | 7 | rs79516966 | 5 | 79711954 | -0.201146039 | 5.55E-05 | 0.9168243 |
| PARP12 | 7 | rs77374532 | 4 | 69934659 | 0.171606604 | 5.80E-05 | 0.918110223 |
| PARP12 | 7 | rs714582 | 7 | 107832881 | 0.12715379 | 5.99E-05 | 0.918110223 |
| PARP12 | 7 | rs11235533 | 11 | 72557668 | 0.184782718 | 5.85E-05 | 0.918110223 |
| PARP12 | 7 | rs10777868 | 12 | 97463359 | -0.100360296 | 5.83E-05 | 0.918110223 |
| PARP12 | 7 | rs12105498 | 2 | 35680928 | -0.163184792 | 9.80E-05 | 0.924166007 |
| PARP12 | 7 | rs2028135 | 2 | 80711329 | 0.165433688 | 8.07E-05 | 0.924166007 |
| PARP12 | 7 | rs4848429 | 2 | 116969469 | 0.101393231 | 7.10E-05 | 0.924166007 |
| PARP12 | 7 | rs6756225 | 2 | 121341180 | 0.166048768 | 8.71E-05 | 0.924166007 |
| PARP12 | 7 | rs17006394 | 2 | 121360536 | 0.165305727 | 8.80E-05 | 0.924166007 |
| PARP12 | 7 | rs2304560 | 2 | 121458843 | 0.164615123 | 9.43E-05 | 0.924166007 |
| PARP12 | 7 | rs6704587 | 2 | 121508348 | 0.164615123 | 9.43E-05 | 0.924166007 |
| PARP12 | 7 | rs10209628 | 2 | 121608436 | 0.16871894 | 7.46E-05 | 0.924166007 |
| PARP12 | 7 | rs12513188 | 4 | 70135074 | 0.114227021 | 7.13E-05 | 0.924166007 |
| PARP12 | 7 | rs75646154 | 4 | 180807979 | 0.179178754 | 8.45E-05 | 0.924166007 |

**Supplementary Table 2l.** *Trans*-eQTL identified for the *PARP12* in non-involved lung tissue of 408 Italian lung adenocarcinoma patients ($P < 1.0 \times 10^{-4}$), listed in order of *P*-value.

| gene | gene chr | SNP | SNP chr | SNP position | beta | p-value | FDR |
|---|---|---|---|---|---|---|---|
| PARP12 | 7 | rs7657018 | 4 | 180836269 | 0.149587527 | 6.81E-05 | 0.924166007 |
| PARP12 | 7 | rs116027742 | 5 | 60539619 | -0.147236323 | 6.66E-05 | 0.924166007 |
| PARP12 | 7 | rs12656371 | 5 | 79711572 | -0.194275918 | 8.90E-05 | 0.924166007 |
| PARP12 | 7 | rs28386947 | 5 | 138642474 | 0.100623661 | 7.90E-05 | 0.924166007 |
| PARP12 | 7 | rs17424324 | 7 | 133111563 | -0.12856203 | 9.75E-05 | 0.924166007 |
| PARP12 | 7 | rs55778057 | 8 | 104772249 | 0.24701984 | 7.19E-05 | 0.924166007 |
| PARP12 | 7 | rs79127267 | 8 | 105492381 | 0.299573699 | 7.11E-05 | 0.924166007 |
| PARP12 | 7 | rs7465467 | 8 | 134974649 | -0.16879437 | 6.55E-05 | 0.924166007 |
| PARP12 | 7 | rs28567030 | 8 | 134983030 | -0.166755988 | 8.07E-05 | 0.924166007 |
| PARP12 | 7 | rs74445624 | 8 | 134983498 | -0.166755988 | 8.07E-05 | 0.924166007 |
| PARP12 | 7 | rs10826467 | 10 | 28411008 | -0.185856152 | 7.45E-05 | 0.924166007 |
| PARP12 | 7 | rs77960080 | 11 | 77431260 | 0.213905548 | 9.06E-05 | 0.924166007 |
| PARP12 | 7 | rs10860211 | 12 | 97460264 | -0.097609653 | 9.64E-05 | 0.924166007 |
| PARP12 | 7 | rs73170494 | 13 | 37919884 | -0.134719671 | 9.96E-05 | 0.924166007 |
| PARP12 | 7 | rs7208415 | 17 | 37132758 | -0.117676531 | 7.79E-05 | 0.924166007 |
| PARP12 | 7 | rs2577075 | 18 | 31142076 | -0.114671112 | 9.57E-05 | 0.924166007 |
| PARP12 | 7 | rs9613255 | 22 | 26708742 | -0.094144135 | 8.09E-05 | 0.924166007 |
| PARP12 | 7 | rs9613258 | 22 | 26717734 | 0.093011463 | 8.05E-05 | 0.924166007 |

**Supplementary Table 2m.** *Trans*-eQTL identified for the *PARP14* in non-involved lung tissue of 408 Italian lung adenocarcinoma patients ($P < 1.0 \times 10^{-4}$), listed in order of *P*-value.

| gene | gene chr | SNP | SNP chr | SNP position | beta | p-value | FDR |
|---|---|---|---|---|---|---|---|
| PARP14 | 3 | rs4615951 | 10 | 106170358 | -0.201495709 | 6.69E-07 | 0.65901749 |
| PARP14 | 3 | rs77963749 | 2 | 203055178 | -0.391780659 | 7.27E-07 | 0.65901749 |
| PARP14 | 3 | rs17820810 | 10 | 106156255 | -0.19894357 | 1.07E-06 | 0.65901749 |
| PARP14 | 3 | rs10491355 | 5 | 109683579 | -0.149273217 | 6.10E-06 | 0.84453778 |
| PARP14 | 3 | rs2113296 | 16 | 83226676 | 0.15884773 | 9.10E-06 | 0.84453778 |
| PARP14 | 3 | rs1021318 | 4 | 168419488 | 0.102671601 | 9.30E-06 | 0.84453778 |
| PARP14 | 3 | rs4916266 | 1 | 172481655 | 0.10183178 | 9.50E-06 | 0.84453778 |
| PARP14 | 3 | rs1411827 | 10 | 106185184 | -0.194044491 | 1.06E-05 | 0.84453778 |
| PARP14 | 3 | rs2310313 | 2 | 102594085 | -0.10592601 | 1.18E-05 | 0.84453778 |
| PARP14 | 3 | rs77488375 | 10 | 88811497 | -0.182960677 | 1.49E-05 | 0.848481351 |
| PARP14 | 3 | rs11621581 | 14 | 92336216 | -0.117309877 | 1.97E-05 | 0.863488543 |
| PARP14 | 3 | rs1076510 | 5 | 109692724 | -0.140321027 | 2.03E-05 | 0.863488543 |
| PARP14 | 3 | rs12187371 | 5 | 26380625 | -0.14810377 | 2.17E-05 | 0.863488543 |
| PARP14 | 3 | rs115755791 | 4 | 76798881 | -0.291533702 | 2.38E-05 | 0.863488543 |
| PARP14 | 3 | rs79732314 | 1 | 237804843 | -0.409487551 | 2.52E-05 | 0.863488543 |
| PARP14 | 3 | rs12474526 | 2 | 100533157 | -0.101071704 | 2.54E-05 | 0.863488543 |
| PARP14 | 3 | rs1452333 | 9 | 28120386 | -0.11345148 | 3.41E-05 | 0.875040765 |
| PARP14 | 3 | rs73442485 | 18 | 40762151 | 0.200533494 | 3.52E-05 | 0.875040765 |
| PARP14 | 3 | rs77469194 | 8 | 123313472 | -0.417084449 | 3.73E-05 | 0.879948067 |
| PARP14 | 3 | rs4448808 | 13 | 38727987 | 0.359146513 | 3.97E-05 | 0.894226918 |
| PARP14 | 3 | rs17691363 | 9 | 74617578 | -0.259735071 | 3.99E-05 | 0.894226918 |
| PARP14 | 3 | rs2147289 | 10 | 7202245 | 0.096769067 | 4.18E-05 | 0.896458005 |
| PARP14 | 3 | rs1933500 | 6 | 104612194 | -0.197316702 | 4.22E-05 | 0.896458005 |
| PARP14 | 3 | rs11170231 | 12 | 52674849 | 0.129969153 | 4.38E-05 | 0.906462302 |
| PARP14 | 3 | rs34961270 | 2 | 100464844 | 0.131127144 | 4.42E-05 | 0.90811064 |
| PARP14 | 3 | rs17650116 | 16 | 78149716 | -0.1575303 | 4.48E-05 | 0.90811064 |
| PARP14 | 3 | rs114473153 | 6 | 11133407 | -0.355354868 | 4.53E-05 | 0.909284759 |
| PARP14 | 3 | rs10825299 | 10 | 54302557 | -0.129168627 | 4.73E-05 | 0.912369094 |
| PARP14 | 3 | rs1543786 | 11 | 128780876 | 0.176738878 | 4.94E-05 | 0.914201928 |
| PARP14 | 3 | rs9499892 | 6 | 104608560 | -0.194106256 | 4.96E-05 | 0.914201928 |
| PARP14 | 3 | rs7662668 | 4 | 16396302 | 0.123098953 | 5.49E-05 | 0.91575718 |
| PARP14 | 3 | rs17057355 | 13 | 38193322 | -0.185381736 | 5.79E-05 | 0.918110223 |
| PARP14 | 3 | rs10506285 | 12 | 49356805 | -0.221998661 | 5.82E-05 | 0.918110223 |
| PARP14 | 3 | rs10506284 | 12 | 49358868 | -0.221998661 | 5.82E-05 | 0.918110223 |
| PARP14 | 3 | rs80306678 | 12 | 49359151 | -0.221998661 | 5.82E-05 | 0.918110223 |
| PARP14 | 3 | rs72962946 | 6 | 118698119 | 0.119310389 | 5.89E-05 | 0.918110223 |
| PARP14 | 3 | rs11737393 | 4 | 137222729 | -0.108162081 | 6.15E-05 | 0.919202897 |
| PARP14 | 3 | rs3944003 | 7 | 117818436 | -0.114155213 | 7.06E-05 | 0.924166007 |
| PARP14 | 3 | rs870638 | 2 | 21261882 | -0.09625758 | 7.16E-05 | 0.924166007 |
| PARP14 | 3 | rs79705514 | 8 | 16770760 | -0.196550064 | 7.22E-05 | 0.924166007 |
| PARP14 | 3 | rs4698481 | 4 | 16409226 | 0.121451101 | 7.47E-05 | 0.924166007 |
| PARP14 | 3 | rs61249850 | 1 | 39524784 | 0.167439201 | 7.61E-05 | 0.924166007 |
| PARP14 | 3 | rs2505098 | 10 | 30144661 | -0.224611071 | 7.69E-05 | 0.924166007 |
| PARP14 | 3 | rs1885411 | 20 | 13274942 | -0.092603795 | 7.70E-05 | 0.924166007 |
| PARP14 | 3 | rs56003946 | 14 | 89556989 | -0.293322245 | 7.73E-05 | 0.924166007 |
| PARP14 | 3 | rs17669793 | 18 | 22413957 | 0.165239962 | 7.88E-05 | 0.924166007 |
| PARP14 | 3 | rs12778129 | 10 | 54346874 | -0.125331526 | 7.97E-05 | 0.924166007 |
| PARP14 | 3 | rs768792 | 5 | 104421304 | 0.092943888 | 8.44E-05 | 0.924166007 |

**Supplementary Table 2m.** *Trans*-eQTL identified for the *PARP14* in non-involved lung tissue of 408 Italian lung adenocarcinoma patients ($P < 1.0 \times 10^{-4}$), listed in order of *P*-value.

| gene | gene chr | SNP | SNP chr | SNP position | beta | p-value | FDR |
|---|---|---|---|---|---|---|---|
| PARP14 | 3 | rs6678558 | 1 | 217109801 | -0.124844083 | 8.44E-05 | 0.924166007 |
| PARP14 | 3 | rs2189791 | 12 | 125631015 | -0.093232479 | 8.95E-05 | 0.924166007 |
| PARP14 | 3 | rs508686 | 20 | 3753334 | -0.09947829 | 9.06E-05 | 0.924166007 |
| PARP14 | 3 | rs488483 | 11 | 60824754 | -0.12272582 | 9.08E-05 | 0.924166007 |
| PARP14 | 3 | rs12570292 | 10 | 81995805 | 0.151646367 | 9.53E-05 | 0.924166007 |
| PARP14 | 3 | rs7746706 | 6 | 105977226 | 0.099911488 | 9.62E-05 | 0.924166007 |
| PARP14 | 3 | rs7308310 | 12 | 20731877 | 0.098835129 | 9.74E-05 | 0.924166007 |
| PARP14 | 3 | rs10825320 | 10 | 54396474 | -0.123361036 | 9.77E-05 | 0.924166007 |

**Supplementary Table 2n.** *Trans*-eQTL identified for the *ST3GAL1* in non-involved lung tissue of 408 Italian lung adenocarcinoma patients ($P < 1.0 \times 10^{-4}$), listed in order of *P*-value.

| gene | gene chr | SNP | SNP chr | SNP position | beta | p-value | FDR |
|---|---|---|---|---|---|---|---|
| ST3GAL1 | 8 | rs75490428 | 1 | 90182646 | 0.299641835 | 1.54E-06 | 0.741191087 |
| ST3GAL1 | 8 | rs4077426 | 3 | 24708623 | 0.173397728 | 1.74E-06 | 0.767785475 |
| ST3GAL1 | 8 | rs1078080 | 10 | 116064228 | -0.142914125 | 2.05E-06 | 0.807441011 |
| ST3GAL1 | 8 | rs946327 | 10 | 101127256 | 0.139803827 | 2.20E-06 | 0.80896024 |
| ST3GAL1 | 8 | rs10095624 | 8 | 27324759 | -0.15094731 | 2.58E-06 | 0.812807829 |
| ST3GAL1 | 8 | rs11598215 | 10 | 116067481 | -0.155031505 | 2.52E-06 | 0.812807829 |
| ST3GAL1 | 8 | rs59979264 | 1 | 90182849 | 0.282659997 | 4.95E-06 | 0.84453778 |
| ST3GAL1 | 8 | rs2359318 | 2 | 174783176 | 0.127651269 | 1.09E-05 | 0.84453778 |
| ST3GAL1 | 8 | rs11993843 | 8 | 27324749 | -0.137542971 | 8.36E-06 | 0.84453778 |
| ST3GAL1 | 8 | rs180620 | 10 | 116027245 | -0.154110188 | 7.92E-06 | 0.84453778 |
| ST3GAL1 | 8 | rs10133034 | 14 | 80798517 | -0.123819392 | 7.17E-06 | 0.84453778 |
| ST3GAL1 | 8 | rs72693011 | 14 | 80849550 | -0.125466161 | 5.74E-06 | 0.84453778 |
| ST3GAL1 | 8 | rs1797671 | 14 | 80855450 | -0.12471443 | 6.79E-06 | 0.84453778 |
| ST3GAL1 | 8 | rs1016699 | 14 | 80922780 | -0.122185067 | 1.04E-05 | 0.84453778 |
| ST3GAL1 | 8 | rs180580 | 10 | 116049647 | -0.152526285 | 1.48E-05 | 0.848481351 |
| ST3GAL1 | 8 | rs79633351 | 5 | 107576953 | 0.205456593 | 1.65E-05 | 0.856932848 |
| ST3GAL1 | 8 | rs12693055 | 2 | 174785941 | 0.125236892 | 2.26E-05 | 0.863488543 |
| ST3GAL1 | 8 | rs6765029 | 3 | 24711178 | 0.121283124 | 2.18E-05 | 0.863488543 |
| ST3GAL1 | 8 | rs10517057 | 4 | 42898545 | -0.126233181 | 2.52E-05 | 0.863488543 |
| ST3GAL1 | 8 | rs7781087 | 7 | 89155042 | -0.125402724 | 1.96E-05 | 0.863488543 |
| ST3GAL1 | 8 | rs592420 | 8 | 9902910 | 0.147954134 | 2.36E-05 | 0.863488543 |
| ST3GAL1 | 8 | rs180561 | 10 | 116053801 | -0.144884099 | 2.37E-05 | 0.863488543 |
| ST3GAL1 | 8 | rs752849 | 11 | 47153776 | 0.140051024 | 2.49E-05 | 0.863488543 |
| ST3GAL1 | 8 | rs7359146 | 14 | 98618265 | -0.16823949 | 1.78E-05 | 0.863488543 |
| ST3GAL1 | 8 | rs58791433 | 14 | 98619997 | -0.16823949 | 1.78E-05 | 0.863488543 |
| ST3GAL1 | 8 | rs78831237 | 16 | 85159288 | 0.332767449 | 2.47E-05 | 0.863488543 |
| ST3GAL1 | 8 | rs10414604 | 19 | 14258950 | 0.216051544 | 2.45E-05 | 0.863488543 |
| ST3GAL1 | 8 | rs36079614 | 5 | 39642405 | 0.136717065 | 2.71E-05 | 0.872704818 |
| ST3GAL1 | 8 | rs73208134 | 3 | 131987176 | 0.218925607 | 3.06E-05 | 0.875040765 |
| ST3GAL1 | 8 | rs76653678 | 3 | 131988000 | 0.218925607 | 3.06E-05 | 0.875040765 |
| ST3GAL1 | 8 | rs57027909 | 3 | 131996780 | 0.209152887 | 2.75E-05 | 0.875040765 |
| ST3GAL1 | 8 | rs13185971 | 5 | 39637783 | 0.135776638 | 3.13E-05 | 0.875040765 |
| ST3GAL1 | 8 | rs234932 | 6 | 2220286 | 0.12083884 | 3.11E-05 | 0.875040765 |
| ST3GAL1 | 8 | rs491364 | 8 | 9900720 | 0.14706962 | 3.29E-05 | 0.875040765 |
| ST3GAL1 | 8 | rs72955584 | 11 | 84954664 | -0.318193509 | 3.27E-05 | 0.875040765 |
| ST3GAL1 | 8 | rs11215458 | 11 | 115269131 | 0.288183357 | 2.92E-05 | 0.875040765 |
| ST3GAL1 | 8 | rs17527836 | 14 | 98648084 | -0.160634914 | 3.50E-05 | 0.875040765 |
| ST3GAL1 | 8 | rs530311 | 19 | 14269255 | 0.223977555 | 3.26E-05 | 0.875040765 |
| ST3GAL1 | 8 | rs492049 | 19 | 14279877 | 0.21970268 | 2.98E-05 | 0.875040765 |
| ST3GAL1 | 8 | rs73100482 | 1 | 234028894 | -0.428864545 | 3.81E-05 | 0.879948067 |
| ST3GAL1 | 8 | rs12677170 | 8 | 84534332 | -0.208209618 | 3.75E-05 | 0.879948067 |
| ST3GAL1 | 8 | rs74826154 | 14 | 98645540 | -0.245230739 | 3.81E-05 | 0.879948067 |
| ST3GAL1 | 8 | rs11972318 | 7 | 32093812 | -0.116495647 | 3.90E-05 | 0.89286517 |
| ST3GAL1 | 8 | rs77974706 | 1 | 61469618 | 0.399892754 | 3.99E-05 | 0.894226918 |
| ST3GAL1 | 8 | rs61507843 | 15 | 88535425 | -0.182195653 | 4.06E-05 | 0.896458005 |
| ST3GAL1 | 8 | rs2044482 | 8 | 27226026 | -0.128145719 | 4.49E-05 | 0.90811064 |
| ST3GAL1 | 8 | rs139887662 | X | 11154416 | 0.431375242 | 4.48E-05 | 0.90811064 |
| ST3GAL1 | 8 | rs6908269 | 6 | 2205572 | 0.14625176 | 4.60E-05 | 0.911360524 |

**Supplementary Table 2n.** *Trans*-eQTL identified for the *ST3GAL1* in non-involved lung tissue of 408 Italian lung adenocarcinoma patients ($P < 1.0 \times 10^{-4}$), listed in order of *P*-value.

| gene | gene chr | SNP | SNP chr | SNP position | beta | p-value | FDR |
|---|---|---|---|---|---|---|---|
| ST3GAL1 | 8 | rs4959171 | 6 | 2362508 | -0.129138484 | 4.75E-05 | 0.912369094 |
| ST3GAL1 | 8 | rs71506796 | 9 | 24241765 | 0.285654177 | 5.03E-05 | 0.914201928 |
| ST3GAL1 | 8 | rs12901500 | 15 | 53519399 | -0.117074213 | 4.94E-05 | 0.914201928 |
| ST3GAL1 | 8 | rs901968 | 15 | 93458891 | 0.123554708 | 5.16E-05 | 0.914201928 |
| ST3GAL1 | 8 | rs7185361 | 16 | 10613718 | 0.124621292 | 5.07E-05 | 0.914201928 |
| ST3GAL1 | 8 | rs4780315 | 16 | 10623710 | 0.123917818 | 5.13E-05 | 0.914201928 |
| ST3GAL1 | 8 | rs9535531 | 13 | 50926440 | -0.113729021 | 5.40E-05 | 0.91575718 |
| ST3GAL1 | 8 | rs56112309 | 15 | 88515844 | -0.183456799 | 5.36E-05 | 0.91575718 |
| ST3GAL1 | 8 | rs1598962 | 1 | 234047233 | -0.329895621 | 5.81E-05 | 0.918110223 |
| ST3GAL1 | 8 | rs17193050 | 3 | 109388064 | 0.155930223 | 5.71E-05 | 0.918110223 |
| ST3GAL1 | 8 | rs234913 | 6 | 2209776 | 0.116539481 | 6.07E-05 | 0.918110223 |
| ST3GAL1 | 8 | rs4752817 | 11 | 46977396 | 0.133751083 | 5.83E-05 | 0.918110223 |
| ST3GAL1 | 8 | rs1725872 | 12 | 77651416 | 0.150483945 | 6.10E-05 | 0.918110223 |
| ST3GAL1 | 8 | rs680705 | 13 | 50869179 | -0.121837698 | 6.15E-05 | 0.919202897 |
| ST3GAL1 | 8 | rs8062377 | 16 | 10613015 | 0.123362737 | 6.17E-05 | 0.919202897 |
| ST3GAL1 | 8 | rs77589855 | 14 | 73664605 | 0.370675493 | 6.21E-05 | 0.919360784 |
| ST3GAL1 | 8 | rs804135 | 1 | 14804374 | 0.113002969 | 9.98E-05 | 0.924166007 |
| ST3GAL1 | 8 | rs115524498 | 2 | 18659661 | 0.370812254 | 7.01E-05 | 0.924166007 |
| ST3GAL1 | 8 | rs199640262 | 3 | 24705779 | 0.153563495 | 6.63E-05 | 0.924166007 |
| ST3GAL1 | 8 | rs4077425 | 3 | 24708470 | 0.11359593 | 9.54E-05 | 0.924166007 |
| ST3GAL1 | 8 | rs769520 | 3 | 109432862 | 0.14900333 | 8.46E-05 | 0.924166007 |
| ST3GAL1 | 8 | rs73881930 | 3 | 176953546 | 0.210670934 | 9.89E-05 | 0.924166007 |
| ST3GAL1 | 8 | rs61432191 | 3 | 181602375 | 0.147994872 | 9.94E-05 | 0.924166007 |
| ST3GAL1 | 8 | rs111836141 | 4 | 67013789 | 0.264505336 | 7.56E-05 | 0.924166007 |
| ST3GAL1 | 8 | rs323691 | 5 | 2557667 | 0.209404827 | 7.88E-05 | 0.924166007 |
| ST3GAL1 | 8 | rs2048010 | 5 | 39675508 | 0.13112361 | 6.81E-05 | 0.924166007 |
| ST3GAL1 | 8 | rs2505673 | 6 | 2355995 | 0.132889726 | 8.41E-05 | 0.924166007 |
| ST3GAL1 | 8 | rs12191002 | 6 | 2361836 | 0.133569759 | 9.61E-05 | 0.924166007 |
| ST3GAL1 | 8 | rs3127661 | 6 | 106694052 | 0.116282153 | 8.03E-05 | 0.924166007 |
| ST3GAL1 | 8 | rs3016538 | 6 | 161815650 | 0.125504766 | 8.53E-05 | 0.924166007 |
| ST3GAL1 | 8 | rs117609876 | 7 | 56251541 | -0.29603543 | 9.37E-05 | 0.924166007 |
| ST3GAL1 | 8 | rs111425321 | 8 | 3194889 | 0.315623022 | 9.16E-05 | 0.924166007 |
| ST3GAL1 | 8 | rs6557990 | 8 | 27288829 | -0.133203512 | 6.78E-05 | 0.924166007 |
| ST3GAL1 | 8 | rs7025985 | 9 | 10572534 | -0.123848259 | 7.84E-05 | 0.924166007 |
| ST3GAL1 | 8 | rs1334044 | 9 | 10995168 | 0.363002417 | 8.77E-05 | 0.924166007 |
| ST3GAL1 | 8 | rs180554 | 10 | 116059049 | -0.119522938 | 9.67E-05 | 0.924166007 |
| ST3GAL1 | 8 | rs180552 | 10 | 116059331 | -0.119073999 | 9.42E-05 | 0.924166007 |
| ST3GAL1 | 8 | rs55805335 | 11 | 36777199 | -0.23655536 | 9.29E-05 | 0.924166007 |
| ST3GAL1 | 8 | rs10891817 | 11 | 115265994 | 0.264675495 | 7.20E-05 | 0.924166007 |
| ST3GAL1 | 8 | rs1940406 | 11 | 132506614 | 0.149914939 | 8.99E-05 | 0.924166007 |
| ST3GAL1 | 8 | rs57898486 | 13 | 113860854 | -0.247137699 | 8.34E-05 | 0.924166007 |
| ST3GAL1 | 8 | rs8020294 | 14 | 22537823 | -0.226647173 | 8.60E-05 | 0.924166007 |
| ST3GAL1 | 8 | rs228125 | 14 | 80871724 | 0.110985239 | 9.37E-05 | 0.924166007 |
| ST3GAL1 | 8 | rs62001352 | 15 | 27388527 | -0.150865971 | 7.71E-05 | 0.924166007 |
| ST3GAL1 | 8 | rs739921 | 17 | 50642229 | 0.123345819 | 9.30E-05 | 0.924166007 |
| ST3GAL1 | 8 | rs12453935 | 17 | 61820094 | 0.139608318 | 9.56E-05 | 0.924166007 |
| ST3GAL1 | 8 | rs61733889 | 18 | 5956382 | 0.26192113 | 8.22E-05 | 0.924166007 |
| ST3GAL1 | 8 | rs7234193 | 18 | 73800665 | 0.17092308 | 9.19E-05 | 0.924166007 |

**Supplementary Table 2n.** *Trans*-eQTL identified for the *ST3GAL1* in non-involved lung tissue of 408 Italian lung adenocarcinoma patients ($P < 1.0 \times 10^{-4}$), listed in order of *P*-value.

| gene | gene chr | SNP | SNP chr | SNP position | beta | p-value | FDR |
|---|---|---|---|---|---|---|---|
| ST3GAL1 | 8 | rs12858390 | X | 84014298 | 0.265634908 | 8.36E-05 | 0.924166007 |
| ST3GAL1 | 8 | rs12689966 | X | 84193110 | 0.121486058 | 8.74E-05 | 0.924166007 |
| ST3GAL1 | 8 | rs12689966 | X | 84193110 | 0.121486058 | 8.74E-05 | 0.924166007 |
| ST3GAL1 | 8 | rs12689966 | X | 84193110 | 0.121486058 | 8.74E-05 | 0.924166007 |
| ST3GAL1 | 8 | rs12689966 | X | 84193110 | 0.121486058 | 8.74E-05 | 0.924166007 |

**Supplementary Table 2o.** *Trans*-eQTL identified for the *TMPRSS2* in non-involved lung tissue of 408 Italian lung adenocarcinoma patients ($P < 1.0 \times 10^{-4}$), listed in order of *P*-value.

| gene | gene chr | SNP | SNP chr | SNP position | beta | p-value | FDR |
|---|---|---|---|---|---|---|---|
| TMPRSS2 | 21 | rs28732378 | 3 | 85354742 | -0.173014218 | 3.63E-06 | 0.84453778 |
| TMPRSS2 | 21 | rs2946394 | 4 | 23906305 | 0.151215242 | 7.67E-06 | 0.84453778 |
| TMPRSS2 | 21 | rs11134868 | 5 | 174711703 | -0.152119702 | 8.79E-06 | 0.84453778 |
| TMPRSS2 | 21 | rs11055449 | 12 | 13450669 | 0.341797055 | 9.17E-06 | 0.84453778 |
| TMPRSS2 | 21 | rs9513827 | 13 | 100949908 | 0.169340347 | 2.99E-06 | 0.84453778 |
| TMPRSS2 | 21 | rs2803213 | 13 | 100952196 | 0.163242161 | 7.43E-06 | 0.84453778 |
| TMPRSS2 | 21 | rs2765328 | 13 | 100953160 | 0.158696104 | 8.04E-06 | 0.84453778 |
| TMPRSS2 | 21 | rs17236868 | 15 | 34356446 | 0.331289393 | 7.09E-06 | 0.84453778 |
| TMPRSS2 | 21 | rs569233 | 19 | 40163445 | 0.158011305 | 3.68E-06 | 0.84453778 |
| TMPRSS2 | 21 | rs6982161 | 8 | 101972051 | 0.144420642 | 1.26E-05 | 0.848481351 |
| TMPRSS2 | 21 | rs72719385 | 9 | 29050491 | 0.34035853 | 1.45E-05 | 0.848481351 |
| TMPRSS2 | 21 | rs2218343 | 17 | 13698368 | 0.357347965 | 1.61E-05 | 0.853417593 |
| TMPRSS2 | 21 | rs656709 | 1 | 107530965 | 0.177912079 | 1.97E-05 | 0.863488543 |
| TMPRSS2 | 21 | rs4953722 | 2 | 109664198 | 0.149261347 | 2.20E-05 | 0.863488543 |
| TMPRSS2 | 21 | rs11941221 | 4 | 5979277 | 0.198427576 | 2.20E-05 | 0.863488543 |
| TMPRSS2 | 21 | rs1425576 | 4 | 38078886 | -0.143700014 | 2.66E-05 | 0.863488543 |
| TMPRSS2 | 21 | rs12549425 | 8 | 726191 | -0.164154961 | 2.58E-05 | 0.863488543 |
| TMPRSS2 | 21 | rs7922735 | 10 | 117412160 | -0.239407486 | 2.21E-05 | 0.863488543 |
| TMPRSS2 | 21 | rs11055458 | 12 | 13463601 | 0.338520358 | 1.74E-05 | 0.863488543 |
| TMPRSS2 | 21 | rs7302344 | 12 | 26335249 | 0.17114423 | 2.59E-05 | 0.863488543 |
| TMPRSS2 | 21 | rs4931255 | 12 | 30094011 | -0.162191111 | 2.46E-05 | 0.863488543 |
| TMPRSS2 | 21 | rs1021707 | 12 | 124309734 | 0.185990302 | 2.60E-05 | 0.863488543 |
| TMPRSS2 | 21 | rs78970981 | 17 | 13656955 | 0.347808154 | 2.12E-05 | 0.863488543 |
| TMPRSS2 | 21 | rs4804561 | 19 | 11029468 | -0.151627353 | 2.30E-05 | 0.863488543 |
| TMPRSS2 | 21 | rs106082 | 19 | 40089121 | 0.145758319 | 2.20E-05 | 0.863488543 |
| TMPRSS2 | 21 | rs36121075 | 20 | 44489376 | -0.20729138 | 1.96E-05 | 0.863488543 |
| TMPRSS2 | 21 | rs4403590 | 1 | 188313587 | -0.535511257 | 3.55E-05 | 0.875040765 |
| TMPRSS2 | 21 | rs4361414 | 4 | 167681992 | 0.300256351 | 3.25E-05 | 0.875040765 |
| TMPRSS2 | 21 | rs9365344 | 6 | 161898113 | -0.134396239 | 2.97E-05 | 0.875040765 |
| TMPRSS2 | 21 | rs1332810 | 9 | 9454258 | 0.191310506 | 3.26E-05 | 0.875040765 |
| TMPRSS2 | 21 | rs10832169 | 11 | 14044939 | -0.140323677 | 3.07E-05 | 0.875040765 |
| TMPRSS2 | 21 | rs2618512 | 11 | 14054400 | -0.139140425 | 3.34E-05 | 0.875040765 |
| TMPRSS2 | 21 | rs16909124 | 12 | 13465015 | 0.330101386 | 2.83E-05 | 0.875040765 |
| TMPRSS2 | 21 | rs2290044 | 12 | 18496375 | -0.173451799 | 2.99E-05 | 0.875040765 |
| TMPRSS2 | 21 | rs7961432 | 12 | 54372607 | -0.234199023 | 3.04E-05 | 0.875040765 |
| TMPRSS2 | 21 | rs80210275 | 18 | 30280296 | -0.310931076 | 3.64E-05 | 0.875040765 |
| TMPRSS2 | 21 | rs73286333 | 12 | 13449176 | 0.332822024 | 3.72E-05 | 0.879948067 |
| TMPRSS2 | 21 | rs1491066 | 5 | 125469534 | -0.139510385 | 3.89E-05 | 0.89286517 |
| TMPRSS2 | 21 | rs35350746 | 2 | 233078998 | -0.145114525 | 3.95E-05 | 0.894226918 |
| TMPRSS2 | 21 | rs818215 | 3 | 85322250 | -0.138250611 | 4.01E-05 | 0.894226918 |
| TMPRSS2 | 21 | rs7829974 | 8 | 101992630 | 0.135824829 | 4.14E-05 | 0.896458005 |
| TMPRSS2 | 21 | rs12949732 | 17 | 41701367 | 0.470789197 | 4.22E-05 | 0.896458005 |
| TMPRSS2 | 21 | rs76476899 | 18 | 30298583 | -0.394001037 | 4.49E-05 | 0.90811064 |
| TMPRSS2 | 21 | rs12376687 | 9 | 9442794 | 0.190269485 | 4.72E-05 | 0.912369094 |
| TMPRSS2 | 21 | rs8192474 | 17 | 32488663 | -0.144073703 | 4.77E-05 | 0.912369094 |
| TMPRSS2 | 21 | rs6103662 | 20 | 44164045 | -0.15560559 | 4.67E-05 | 0.912369094 |
| TMPRSS2 | 21 | rs115457413 | 2 | 59035335 | -0.241959813 | 5.08E-05 | 0.914201928 |
| TMPRSS2 | 21 | rs67469048 | 5 | 174697681 | -0.155490183 | 5.46E-05 | 0.91575718 |

**Supplementary Table 2o.** *Trans*-eQTL identified for the *TMPRSS2* in non-involved lung tissue of 408 Italian lung adenocarcinoma patients ($P < 1.0 \times 10^{-4}$), listed in order of *P*-value.

| gene | gene chr | SNP | SNP chr | SNP position | beta | p-value | FDR |
|---|---|---|---|---|---|---|---|
| TMPRSS2 | 21 | rs1486343 | 12 | 38648610 | -0.206755459 | 5.28E-05 | 0.91575718 |
| TMPRSS2 | 21 | rs74384381 | 17 | 13669076 | 0.326353683 | 5.41E-05 | 0.91575718 |
| TMPRSS2 | 21 | rs1882535 | 12 | 81619319 | 0.374704375 | 5.54E-05 | 0.916497699 |
| TMPRSS2 | 21 | rs201434136 | 12 | 81647065 | 0.374704375 | 5.54E-05 | 0.916497699 |
| TMPRSS2 | 21 | rs12506469 | 4 | 124450032 | 0.145723972 | 5.80E-05 | 0.918110223 |
| TMPRSS2 | 21 | rs10484888 | 6 | 10837021 | -0.297796922 | 5.74E-05 | 0.918110223 |
| TMPRSS2 | 21 | rs62432531 | 6 | 138617688 | -0.279562791 | 6.01E-05 | 0.918110223 |
| TMPRSS2 | 21 | rs11786361 | 8 | 101986859 | -0.137727175 | 5.84E-05 | 0.918110223 |
| TMPRSS2 | 21 | rs11023056 | 11 | 14030572 | -0.135548359 | 5.96E-05 | 0.918110223 |
| TMPRSS2 | 21 | rs115829889 | 14 | 42343809 | -0.489482426 | 5.74E-05 | 0.918110223 |
| TMPRSS2 | 21 | rs8024987 | 15 | 32044027 | 0.157875324 | 6.10E-05 | 0.918110223 |
| TMPRSS2 | 21 | rs59602121 | 2 | 59578303 | -0.319845835 | 6.66E-05 | 0.924166007 |
| TMPRSS2 | 21 | rs61676395 | 2 | 59582067 | -0.319845835 | 6.66E-05 | 0.924166007 |
| TMPRSS2 | 21 | rs10185001 | 2 | 59599494 | -0.321192688 | 8.29E-05 | 0.924166007 |
| TMPRSS2 | 21 | rs13419857 | 2 | 157945509 | 0.181953565 | 6.37E-05 | 0.924166007 |
| TMPRSS2 | 21 | rs6707765 | 2 | 190175107 | 0.186606882 | 8.16E-05 | 0.924166007 |
| TMPRSS2 | 21 | rs60805365 | 2 | 205989042 | -0.142697044 | 9.80E-05 | 0.924166007 |
| TMPRSS2 | 21 | rs40160 | 3 | 142976959 | 0.139159685 | 8.67E-05 | 0.924166007 |
| TMPRSS2 | 21 | rs4832980 | 4 | 38008672 | -0.137273231 | 7.51E-05 | 0.924166007 |
| TMPRSS2 | 21 | rs7698019 | 4 | 136887804 | -0.339865126 | 7.29E-05 | 0.924166007 |
| TMPRSS2 | 21 | rs13147308 | 4 | 174672336 | -0.174786998 | 7.01E-05 | 0.924166007 |
| TMPRSS2 | 21 | rs12648678 | 4 | 174677129 | -0.174786998 | 7.01E-05 | 0.924166007 |
| TMPRSS2 | 21 | rs200250231 | 5 | 174707349 | -0.132089505 | 7.76E-05 | 0.924166007 |
| TMPRSS2 | 21 | rs2715140 | 7 | 50625014 | -0.20513089 | 9.50E-05 | 0.924166007 |
| TMPRSS2 | 21 | rs13262331 | 8 | 5010531 | -0.160082415 | 8.51E-05 | 0.924166007 |
| TMPRSS2 | 21 | rs72719371 | 9 | 29002266 | 0.251684787 | 7.60E-05 | 0.924166007 |
| TMPRSS2 | 21 | rs7096762 | 10 | 51933907 | -0.156604698 | 9.47E-05 | 0.924166007 |
| TMPRSS2 | 21 | rs11000191 | 10 | 51935207 | -0.156604698 | 9.47E-05 | 0.924166007 |
| TMPRSS2 | 21 | rs12282656 | 11 | 69248261 | 0.147476376 | 8.69E-05 | 0.924166007 |
| TMPRSS2 | 21 | rs57906526 | 12 | 2430594 | 0.235350909 | 8.53E-05 | 0.924166007 |
| TMPRSS2 | 21 | rs12228908 | 12 | 38373369 | -0.204990671 | 7.74E-05 | 0.924166007 |
| TMPRSS2 | 21 | rs12582802 | 12 | 39591404 | 0.22365248 | 8.05E-05 | 0.924166007 |
| TMPRSS2 | 21 | rs11066827 | 12 | 113935240 | 0.205429148 | 7.66E-05 | 0.924166007 |
| TMPRSS2 | 21 | rs8022905 | 14 | 75316498 | -0.169073882 | 6.98E-05 | 0.924166007 |
| TMPRSS2 | 21 | rs710962 | 17 | 31612357 | 0.134479468 | 6.82E-05 | 0.924166007 |
| TMPRSS2 | 21 | rs79036712 | 18 | 30442690 | -0.26516445 | 6.71E-05 | 0.924166007 |
| TMPRSS2 | 21 | rs58387531 | 18 | 51798655 | 0.328314173 | 9.19E-05 | 0.924166007 |
| TMPRSS2 | 21 | rs947202 | 20 | 43150303 | 0.287381665 | 9.28E-05 | 0.924166007 |
| TMPRSS2 | 21 | rs135277 | 22 | 48070053 | 0.13875338 | 7.38E-05 | 0.924166007 |
