## Supplemental Table 3 for "Lung expression of genes encoding SARS-CoV-2 cell entry molecules and antiviral restriction factors: interindividual differences are associated with age and germline variants"

**Supplementary Table 3**. Minor allele frequencies of the top 16 *cis*-eQTL SNPs in different populations (according to the 1000 Genomes Project Phase 3 [39], as reported in Ensembl genome browser) and in this study

| SNP | Minor allele | Minor allele frequency (%) | | | | | |
|---|---|---|---|---|---|---|---|
| | | African | American | East Asian | South Asian | European | This study |
| **APOBEC3D** | | | | | | | |
| rs139296 | A | 31 | 28 | 25 | 40 | 34 | 32 |
| rs9611092 | A | 32 | 28 | 25 | 39 | 34 | 32 |
| rs139331 | A | 32 | 24 | 25 | 39 | 34 | 32 |
| rs5757715 | G | 8 | 32 | 77 | 33 | 27 | 24 |
| rs5750849 | A | 7 | 32 | 77 | 34 | 27 | 24 |
| rs9607655 | A | 2 | 30 | 74 | 27 | 28 | 24 |
| **APOBEC3G** | | | | | | | |
| rs8177832 | G | 43 | 7 | 7 | 1 | 3 | 4 |
| rs17537581 | A | 55 | 12 | 10 | 2 | 8 | 8 |
| rs61362448 | A | 33 | 6 | 2 | 2 | 7 | 9 |
| rs11914001 | A | 51 | 10 | 8 | 4 | 12 | 13 |
| rs139331 | A | 32 | 24 | 25 | 39 | 34 | 32 |
| rs59728386 | A | 47 | 10 | 7 | 4 | 12 | 12 |
| rs738469 | G | 47 | 7 | 1 | 2 | 8 | 9 |
| rs139382 | A | 0 | 1 | 0 | 0 | 2 | 2 |
| rs17209173 | G | 6 | 13 | 1 | 12 | 25 | 21 |
| rs430915 | A | 12 | 52 | 42 | 46 | 49 | 43 |
